# Adaptation in outbred sexual yeast is repeatable, polygenic, and favors rare haplotypes

**DOI:** 10.1101/2021.08.27.457900

**Authors:** Robert A. Linder, Behzad Zabanavar, Arundhati Majumder, Hannah Chiao-Shyan Hoang, Vanessa Genesaret Delgado, Ryan Tran, Vy Thoai La, Simon William Leemans, Anthony D Long

## Abstract

We describe the results of a 200 generation Evolve and Resequence (E&R) study initiated from an outbred dipliod recombined synthetic base population derived from 18 genetically diverse founders. Replicate populations were maintained at large effective population sizes (>10^5^ individuals), exposed to several different chemical challenges over 12 weeks of evolution, and whole-genome resequenced. Weekly forced outcrossing implies a per gene per cell-division recombination rate higher than that achieved in Drosophila E&R studies. In 55 sexual populations we observe large fitness gains and highly repeatable patterns of genome-wide haplotype change within each chemical challenge. There was little evidence for pervasive pleiotropy, as evidenced by patterns of haplotype change between drug treatments. Within treatment adaptation appears highly polygenic with almost the entire genome showing significant consistent haplotype change. Finally, adaptation was almost always associated with only one of the 18 founder alleles, suggesting selection primarily acts on rare variants private to a founder or haplotype blocks harboring multiple mutations. This observation contradicts the notion that adaptation is often due to subtle frequency shifts at intermediate frequency variants.

## Introduction

A detailed understanding of the genetic mechanisms through which organisms adapt to stressful environments has important implications for a wide range of issues such as climate change, antibiotic resistance, and bioengineering. Studying adaptation in a controlled laboratory environment through experimental evolution has become a mainstay of evolutionary genetics (Garland & Rose, 2009). Such studies have furthered our understanding of the dynamics of evolution on both a phenotypic and genotypic level. A powerful approach to studying evolution in a laboratory setting is the evolve and resequence (E&R) paradigm (Turner et al., 2011), in which populations in a controlled setting are sequenced before and during the course of adaptation to a novel environment to better understand the dynamics of adaptation (reviewed in (Long et al., 2015; Schlotterer et al., 2016)).

Advances made in understanding mechanisms of adaptation have come from E&R studies in microbes, flies, and, more recently, worms. Studies in microbes have been instrumental in furthering our understanding of adaptation in a purely asexual context, which has important implications for the evolution of antibiotic resistance (Barlow & Hall, 2003; Chevereau et al., 2015; Lamrabet et al., 2019; Sousa et al., 2012; Toprak et al., 2011), cancer tumorigenesis (Sprouffske et al., 2012; Taylor et al., 2013, 2017), and the acquisition of novel phenotypes (López-Malo et al., 2015; Wannier et al., 2018; Wildenberg & Murray, 2014). The majority of such studies are initiated from an isogenic haploid asexual population that evolves in a novel environment; thus, adaptation occurs solely via *de novo* mutations and adaptation is dominated by selective sweeps and clonal interference between beneficial mutations on different genetic backgrounds (Good et al., 2017; Lang et al., 2013; Toprak et al., 2011). Under this asexual isogenic paradigm forces such as historical contingency (Toprak et al., 2011), genetic parallelism at the level of genes but not (except for rare exceptions, see (Toprak et al., 2011)) specific mutations (Tenaillon et al., 2012; Toprak et al., 2011), and diminishing returns epistasis (Toprak et al., 2011; Wang et al., 2016) matter a great deal to the evolutionary process. Similar studies in asexual isogenic diploids have shown that ploidy can influence the dynamics of adaptation, with diploids often adapting more slowly than haploids, likely due to the effects of Haldane’s sieve (Fisher et al., 2018; Gerstein et al., 2014; Johnson et al., 2021; Marad et al., 2018; Sellis et al., 2016; Sellis et al., 2011; Zeyl et al., 2003). Diploids also appear to accumulate more potentially deleterious mutations (increased mutational load), and continue to adapt longer than haploids, likely due to the effects of mitotic recombination (Forche et al., 2011; Gerstein et al., 2014; Johnson et al., 2021). Despite these differences, general aspects of adaptation are shared with asexual haploids, including clonal interference and the fixation of beneficial mutations.

However, the vast majority of higher eukaryotes are obligately sexual, thus evolution takes place in the presence of standing variation and recombination. As a result, evolution in asexual isogenics may serve as a poor model for the dynamics of evolution in sexual outbreds. In the presence of standing variation and recombination, adaptation appears to proceed more deterministically, with selection acting on alleles already present in the population (Burke et al., 2010; Burke et al., 2014; Hernandez et al., 2011; Phillips et al., 2016), and *de novo* mutations generally having a minor role in adaptation over short time-scales (Burke et al., 2014). A frequent observation is that sexual populations maintain high levels of heterozygosity (Barghi et al., 2019; Burke et al., 2010; O’Connor et al., 2021; Phillips et al., 2016), even after hundreds of generations of evolution in a constant environment. Further, studies in budding yeast have shown that adaptation with even modest amounts of recombination is capable of making evolution more efficient, both by decoupling deleterious hitchhiking mutations from beneficial mutations and by combining multiple beneficial mutations onto a single genetic background (Gray & Goddard, 2012; McDonald et al., 2016; Zeyl & Bell, 1997). An emerging pattern from Drosophila E&R experiments is that adaptation appears to be fairly polygenic with much of the genome responding to selection (Barghi et al., 2019; Barghi & Schlötterer, 2020; Burke et al., 2010; Phillips et al., 2016).

Despite progress, many questions remain unanswered. E&R experiments in Drosophila often are limited in terms of the number of replicates examined, the number of generations over which an experiment is carried out, and the effective population size that can be efficiently handled (*N_e_* is typically <<1000). Furthermore, base populations are typically initiated from hundreds of uncharacterized founders, so the tracking of haplotypes in an evolved population is a difficult undertaking. Finally, although flies are obligate sexuals they are recombinationally depauperate. Recombination only occurs in females and the number of recombination events per female gamete per generation is ∼5. Since both the recombination rate and *N_e_* is small in fly E&R experiments, the selection coefficients associated with adaptive variants must in turn be fairly large (*s*>>*1/N_e_*) and as a result difficult to characterize haplotype blocks several megabases in size could be responding to selection, especially if rare alleles are being favored. Although arguments have been made that observed patterns of allele frequency change support polygenic adaptation (c.f. Barghi et al., 2020), it is difficult to disentangle polygenic adaptation from “traffic”, the phenomena of several long strongly selected haplotypes trying to simultaneously increase in frequency. It is finally of interest to confirm that patterns of adaptation observed in populations with *N_e_*<<1000 match those seen in populations with *N_e_* approaching those of natural populations.

Budding yeast is a model system ideally suited to help resolve these fundamental questions, due to their facultatively sexual life cycle and the ease of maintaining very large populations with a high degree of replication for many hundreds of generations. Previous work has been instrumental in describing the dynamics of adaptation in outcrossing populations of budding yeast ( Burke et al., 2014; Leu et al., 2020; McDonald et al., 2016; Kosheleva & Desai, 2018), although these studies all employed four or fewer founder haplotypes and did not ask populations to evolve in the presence of several different novel stressors. We extend previous E&R experiments that used an outcrossing population of budding yeast derived from four highly diverged founder strains (the SGRP-4X) carried through twelve rounds of meiosis followed by random mating to break up linkage blocks (Burke et al., 2014; Cubillos et al., 2013; Parts et al., 2011), but instead employ a base population derived from 18 different founders (Linder et al., 2020). This population harbors considerably more standing variation than previously studied budding yeast populations, and allows us to examine patterns of haplotype change over the 18 different starting alleles. Unlike previous work in outbred sexual yeast, we evolve this base population with replication in the presence of different chemical challenges, modeling how short term evolution occurs when an organism encounters a novel environment. Weekly outcrossing, at a rate four times higher per cell division per gene than in Drosophila E&R experiments, shuffles evolving haplotypes throughout the course of evolution, and allows us to better distinguish between polygenic adaptation versus linked hitch-hiking. This study has several advantages relative to previous E&R experiments in sexual outbreds: a large effective population size is maintained (>10^5^), the experiment has a high degree of replication, a wide-range of cellular processes are perturbed, a highly outcrossed base population is employed, and founder haplotypes are known and can be tracked. These features of our model system allow for general inferences about the nature of adaptation in outbred sexual populations.

## Results

### Experimental evolution of budding yeast

Yeast were evolved under a set of conditions chosen to mimic a ∼200 generation burst of adaptation to a novel environment in a recombining diploid outbred population maintaining a large population size. A highly outbred diploid population was used to initiate a sixteen-fold replicated evolve-and-resequence (E&R) experiment that explored the response to 16 different chemical challenges (Table S1) over 12 weeks. A schematic of the weekly evolution regimen is depicted in Figure 1.

**Figure 1.**
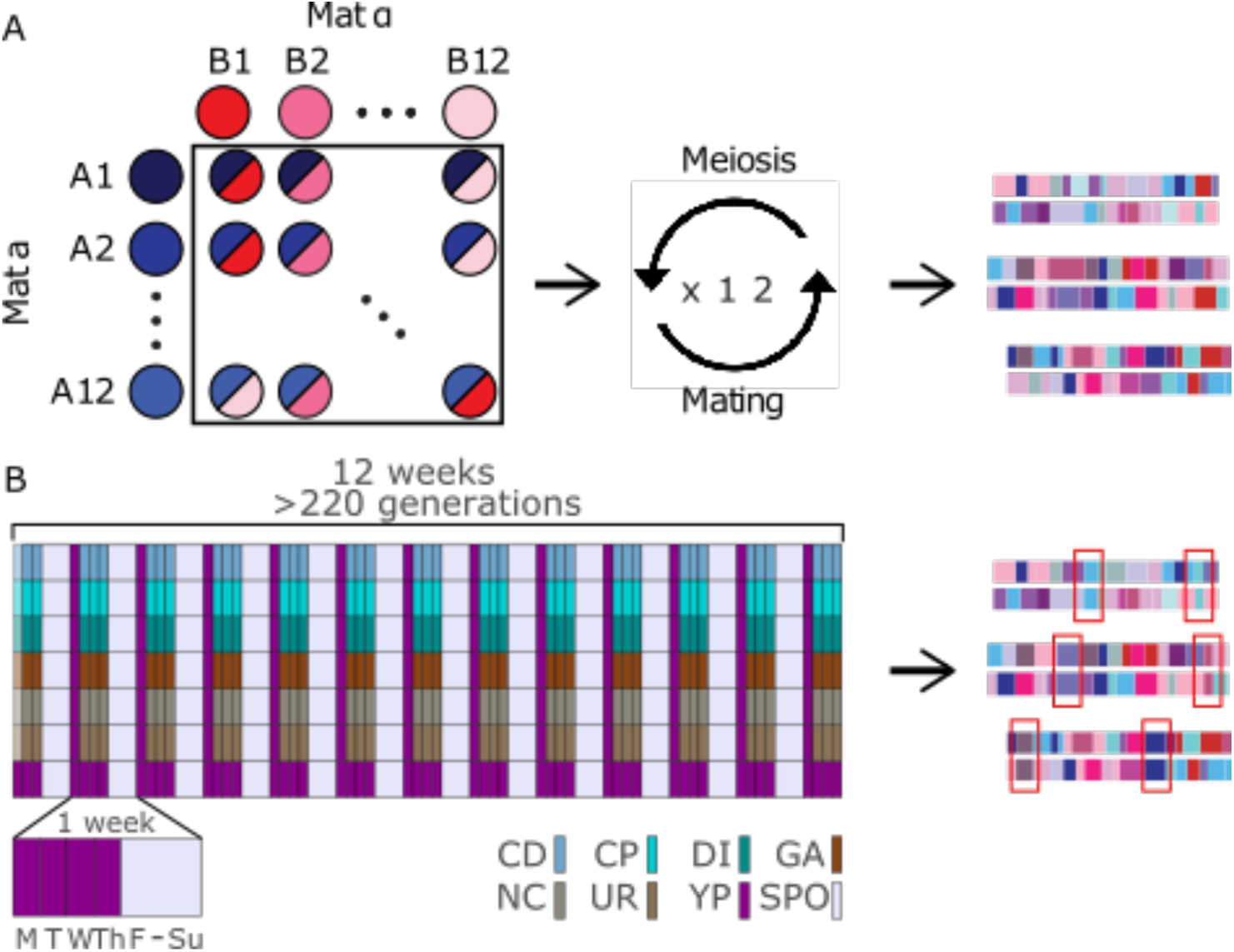
Schematic of the long-term evolution experiment. Panel (A) depicts the creation of the base population used in this study. In total, 11 Mat a and 11 Mat @ strains were crossed in a full diallel, followed by 12 rounds of forced outcrossing to create a highly diverse and mosaic population of diploids. Panel (B) depicts the long-term evolution experiment. Details of the regimen are in the methods. The first day of the experiment, a different, normally lower dose of the chemical stressor was used to acclimate cells to the chemical.

Each week populations experienced daily serial passages Tuesday through Friday in media supplemented with the different chemicals. Every Friday populations were sporulated, spores were recovered on Mondays, mated, and diploids recovered on Tuesday mornings (in total, from Friday to Tuesday, populations underwent one meiotic and six mitotic generations; Figure 1- figure supplement 1). We estimate that the populations were going through roughly 12 cell divisions per week in the presence of the chemical challenges (Figure 1- figure supplement 1B). Since budding yeast experience ∼90 cross-overs per meiosis (Mortimer et al. 1991), the average recombination rate per gene per cell division is estimated to be roughly 0.0008. This recombination rate is ∼4-folder higher than the per gene recombination rate experienced in obligately sexual *Drosophila melanogaster* during experimental evolution. Each evolved population was initiated from >1M cells, the serial transfer bottlenecks were estimated to contain ∼10-20M cells, we recovered ∼3M haploid spores per population and ∼750,000 diploids after mating each Tuesday (Figure 1- figure supplement 1C). Although the census size can be orders of magnitude higher, the effective population size of the experiment is likely close to 750,000. This is a population size orders of magnitude larger than typical outbred sexual E&R experiments, and is perhaps even larger than the effective population size of the human species (Henn et al., 2012).

To examine the genomic response to adaptation we carried out short-read Illumina pooled sequencing of each population, with the base population sequenced to 2226X and the evolved populations to an average of 54X (range 5X to 145X, Figure 1- figure supplement 2). Since the gDNA used to create the libraries was obtained from ∼10^8^ cells, coverage at the 182,460 queried SNPs estimates the frequency of that SNP in the population with a binomial error almost entirely associated with sampling due to sequencing coverage, as opposed to sequencing coverage plus error due to a finite pool of individuals (Burke et al., 2010). We estimated founder haplotype frequencies for the base population and each evolved population at 11,574 loci spaced every 1 kb throughout the genome. There does not appear to be a need to estimate haplotype frequencies at a resolution greater than one per 1kb, as the mean haplotype frequency change between adjacent positions in founder AB3 is very close to zero (estimated at ∼5×10^-7^, Figure 2- figure supplement 1; also c.f. Figure 4 and Figure 9). Previous work further suggests that the average absolute error in the individual haplotype frequency estimates is ∼0.01 at the sequencing coverages we employ. That is, haplotype frequency estimates are associated with much lower errors than would be predicted based solely on coverage and binomial sampling, consistent with other studies (Burke et al., 2014; Kessner et al., 2013; Linder et al., 2020; Long et al., 2011; Tilk et al., 2019).

Since the populations were evolved under conditions that forced outcrossing every 18 cell divisions, at the molecular level we expected the population to maintain high levels of diversity and only display localized changes in allele frequencies and losses of heterozygosity associated with adaptation, a prediction of evolutionary models in outbred sexuals (reviewed in Long et al., 2015). But Manhattan plots of per population heterozygosity and haplotype frequency change suggested that in many replicates, for many drugs, asexual “cheaters” emerged and took over those populations (Figure 2). It appears that a combination of selection, forced meiosis, and aggressive spore recovery using chemical and mechanical shearing can favor the emergence of genotypes that can avoid our desired life-cycle. Despite the emergence of cheaters being an interesting observation, to understand evolution in sexual outbred populations, we identified a subset of chemicals and populations in which cheaters were unable to gain a foothold and sweep to high frequency using a classification metric (see Methods) that allowed us to classify populations into one of three non-overlapping types: aneuploid haploids, clonal diploids, or outbred sexuals (for examples see Figure 2). We observed 105 drug/population combinations out of 221 (∼48%) that appeared to remain sexual throughout our experiment (Table S2). Different chemical challenges are associated with different rates of cheater emergence. Lower 12-week average OD630 (a measure of growth rate) is associated with lower week 12 heterozygosity measures across different chemical treatments (Figure 2- figure supplement 2A), a pattern that similarly manifests as a smaller fraction of populations classified as outbred sexuals as a function of decreasing growth rate (Figure 2- figure supplement 2B). These observations are consistent with the idea that the subset of chemical treatments that presented greater initial selective challenges, as measured by slower growth and lower cell densities, were more prone to invasion by a cheater. We hypothesize that given a very strong selective pressure (or an absence of standing variation) mutations/aneuploidies of large effect can be favored irrespective of deleterious side-effects. A corollary is that weaker selective pressure favored adaptation via standing variation.

**Figure 2.**
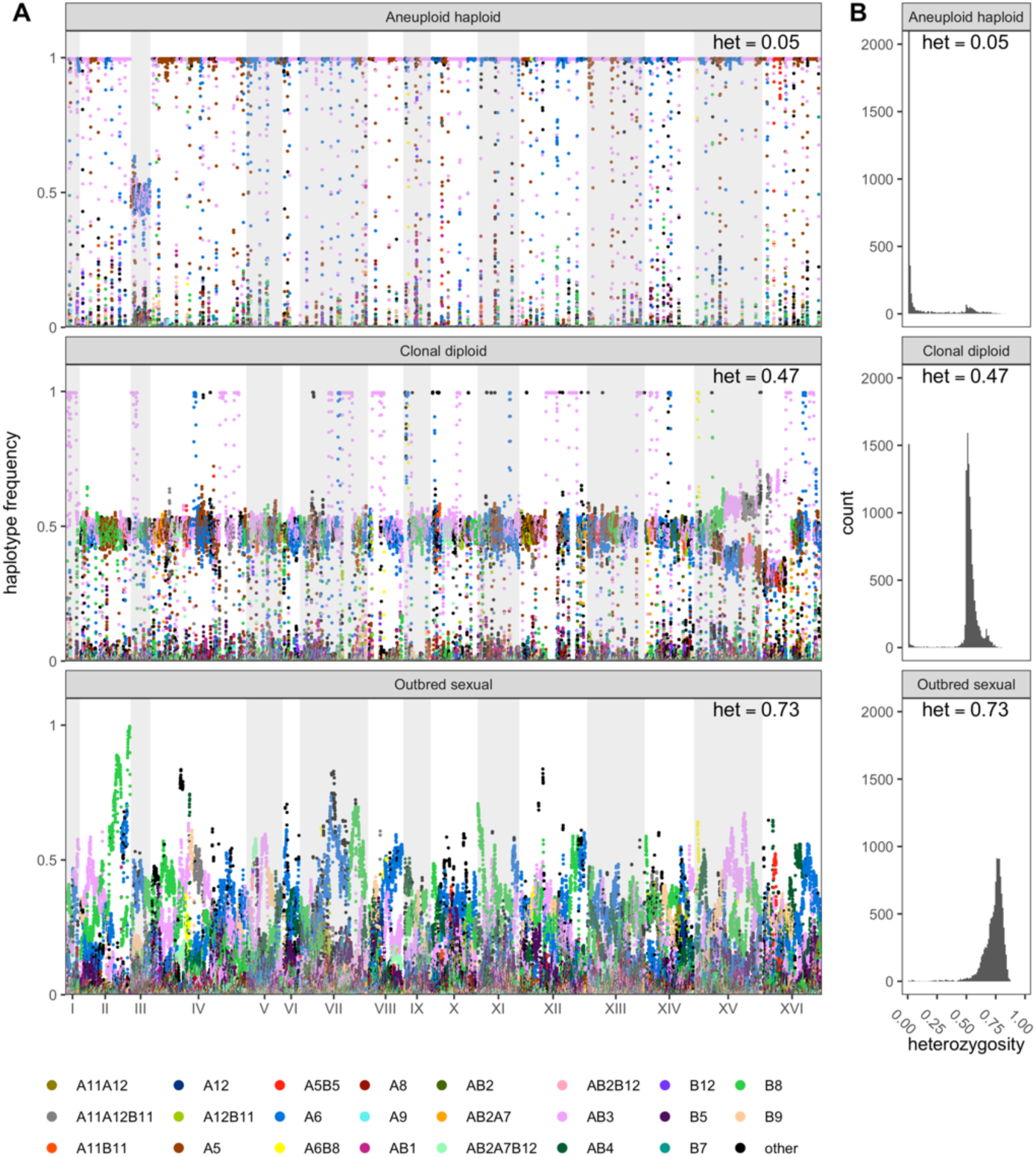
Different classes of evolved populations. In (A), raw haplotype frequency plots are shown of (from top to bottom) a caffeine replicate, a glacial acetic acid replicate, and a cadmium chloride replicate classified as aneuploid haploid, heterozygous clonal, and outbred sexual, respectively. In (B) are shown histograms of the per-site heterozygosity of the corresponding replicates. The average per-site heterozygosity is shown at the top of each panel. **Figure 2- source data 1** **Plots of genome-wide haplotype frequencies.** This zip archive contains all genome-wide raw haplotype frequency plots from each chemical and replicate (same format as Figure 2A). **Figure 2- source data 2** **Plots of per-site heterozygosity.** This zip archive contains all per-site heterozygosity histograms (same format as Figure 2B), with the first subplot being a histogram of the per-site heterozygosity of the base population included in all files as a baseline comparison.

To enable statistical analyses of patterns of evolutionary change in the subset of populations that remained sexual we additionally required that each chemical treatment retain at least 6 replicate sexual populations (excluding the control YPD treatment) and that genome-wide haplotype frequency estimates for populations within a chemical treatment clustered together relative to different treatments in a distance-based cladogram (Figure 2- figure supplement 3). This clustering filter, which removed 23 populations, was an attempt to control for possible cross-contamination events (that cannot be completely eliminated due to the complex weekly transfer regimen). After this aggressive filtering seven chemicals and 55 total replicates (∼25%) remained, and we focus the remainder of the manuscript on these populations (Figure 2- figure supplement 4 and Table 1).

**Table 1.**
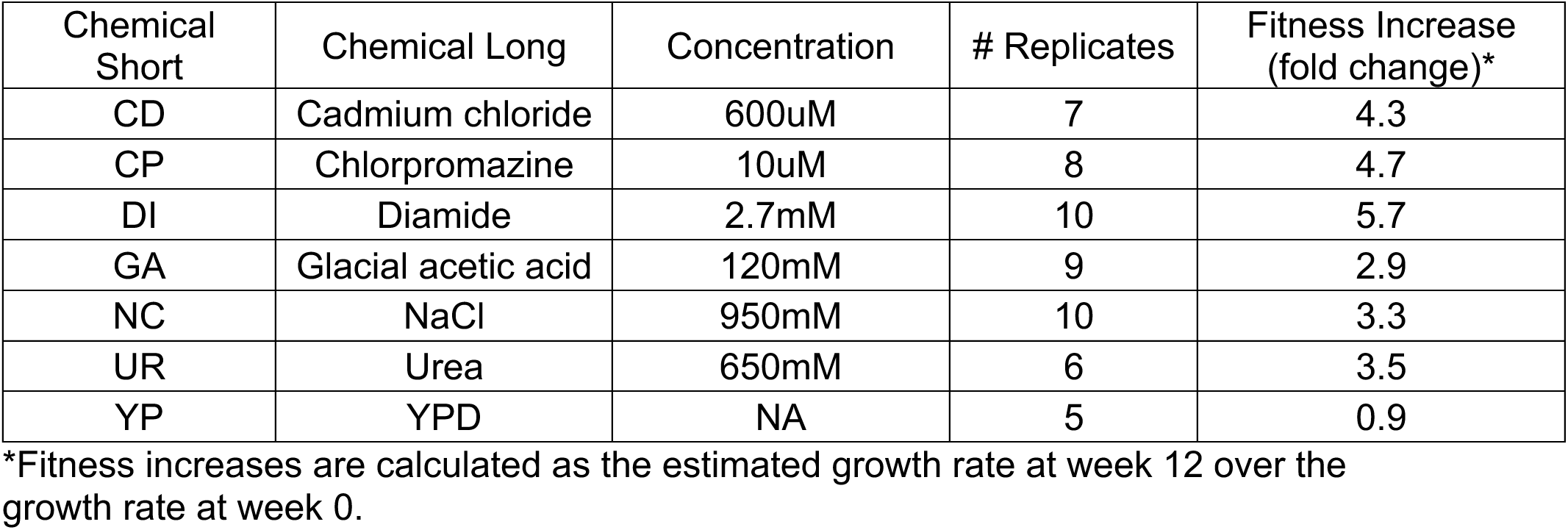
Chemicals and replicates used in this study.

### Fitness increased substantially after 12 weeks of evolution

Like previous experimental evolution studies, we observe considerable increases in fitness over the 12 weeks of the experiment, with the exception of the YPD-only condition, which slightly lost fitness at week 12 (Table 1), potentially due to a fitness trade-off between mitotic growth and mating efficiency, a phenomenon that has been documented previously (Lang et al., 2009) (also see (Li et al., 2019; Zeyl & Bell, 1997), in which there was no observable fitness gain in mitotic growth in the YPD-only condition). Over the six chemical stressors we observe a week twelve increase in fitness ranging from ∼3-fold to slightly less than 6-fold (see methods for details on measuring growth rate). This compares favorably to an experiment in diploid budding yeast carried out for a similar number of generations with recombination, in which fitness was found to increase by ∼1.5-fold (Kosheleva & Desai, 2018) in YPD-only media in a 2-way outcrossed population. Other studies where 2- and 4-way populations of outbred budding yeast were evolved asexually for 50 generations in two different chemicals (rapamycin and hydroxyurea) found fitness gains of ∼1.4 and ∼1.5 respectively (Li et al., 2019). Studies in haploid budding yeast have shown small overall fitness gains after >100 generations of purely asexual evolution from an isogenic base population (Fisher et al., 2018; Kryazhimskiy et al., 2014; Lang et al., 2011; Levy et al., 2015). Further, after 50,000 generations of purely asexual evolution, (Tenaillon et al., 2016) found an average fitness increase of ∼1.7-fold amongst all replicate *Escherichia coli* populations during adaptation to nutrient limitation. These comparisons suggest that the presence of recombination and high levels of standing variation lead to much larger fitness gains than are normally seen in experiments with less variation and/or no recombination, at least over short bursts of evolution in a novel environment, consistent with early studies that have more directly made this observation (Johnson et al., 2021).

### De novo mutations and aneuploidies

Across the seven chemicals used, we detect several different aneuploidies (Table S3). Of note, the entirety of chromosome II was duplicated in all seven cadmium chloride replicates- the well-known cadmium transporter, *PCA1*, is present on this chromosome, which is a suggestive explanation for this particular aneuploidy. Additionally, a single sodium chloride replicate has the entirety of chromosomes III and V duplicated, while three urea replicates have an additional copy of chromosome V. Partial aneuploidies were also detected, including the end of chromosome XVI in half of the chlorpromazine replicates and one at the beginning of chromosome XVI in 6 of 9 glacial acetic acid replicates, as well as a duplication of part of chromosome VI that occurred in all 10 diamide replicates. This last partial duplication had 2-fold increased relative coverage in evolved populations as compared to the base, implying that an additional 2 copies of this region are present in all individuals. Of note, a single partial duplication of the end of chromosome XVI was detected in all conditions with the exception of cadmium chloride. Several founders that increased in frequency at the end of chromosome XVI in the aneuploid populations have naturally occurring duplications that fall within this region, some of which include the multidrug transporter, *SGE1*, a potential candidate driving this particular aneuploidy. Only a single YPD replicate had any detectable aneuploidies (at the end of chromosome XVI), suggesting that the other aneuploidies detected in the six chemical treatments are likely specifically beneficial for the chemical(s) they are detected in. Despite the existence of several presumably adaptive aneuploidies, it is unlikely these aneuploidies account for the bulk of adaptation as they tend to involve a small number of chromosomes or replicates, and we see strong genome-wide responses to selection.

We also detected 219 single nucleotide variants (SNVs) that reached frequencies of 20% or higher in the populations they were detected in that likely spontaneously arose during the course of evolution (Table S4; see Methods for how *de novo* mutations were detected). An average of 31 *de novo* SNVs were detected per condition, from 14 in cadmium chloride to 56 in sodium chloride. These mutations reached an average frequency of 28% over the 12 weeks of evolution, with 20 total SNVs at a frequency at or greater than 50% and 3 total SNVs reaching a frequency of 75% or greater. The presence of multi-hit genes within a single condition are strongly suggestive of mutations that affect the chemicals they are detected in. In total, we found only one gene which was hit in 2 different chlorpromazine replicates (*DSF2*, a gene of unknown function). As with aneuploidies, although a subset of the *de novo* mutations detected are likely adaptive, they are unlikely to explain a large fraction of adaptive change since they generally only occur in a subset of genes and/or replicates. Furthermore, if *de novo* mutations dominated the selective response we would not expect to see strong convergence of haplotype frequencies over replicates as described below. Nonetheless, in our large effective population size E&R experiment, newly arising mutations seem to play a larger role in adaptation compared to past experiments with much smaller values of *N_e_μ*.

### A genome-wide scan for haplotype frequency change suggests highly polygenic adaptation

A genome-wide scan for genomic regions contributing to adaptation was carried out for each of the seven drug treatments to detect regions showing consistent haplotype change relative to the base population. Figure 3, Manhattan plots of genome-wide LOD scores for all chemical treatments, shows that across all treatments the vast majority of the genome is above the significance cutoff. Across all treatments, an average of 95.2**%** of the genome was above the significance threshold (Table S5). A simple linear model regressing the number of replicates per treatment onto the genome-wide average LOD shows that there is no correlation between the number of replicate populations and the average genome-wide significance (Figure 3- figure supplement 1A), suggesting that our test statistic is not strongly impacted by the number of replicates. Similarly, a regression of the average LOD score on the number of haplotypes contributing to a test (Figure 3- figure supplement 1B) suggests that our test statistic is not strongly impacted by the number of haplotypes contributing to a test. Overall patterns of haplotype frequency change suggest that adaptation is highly polygenic, with nearly all regions of the genome being either directly under selection or in linkage disequilibrium (LD) with selected regions.

**Figure 3.**
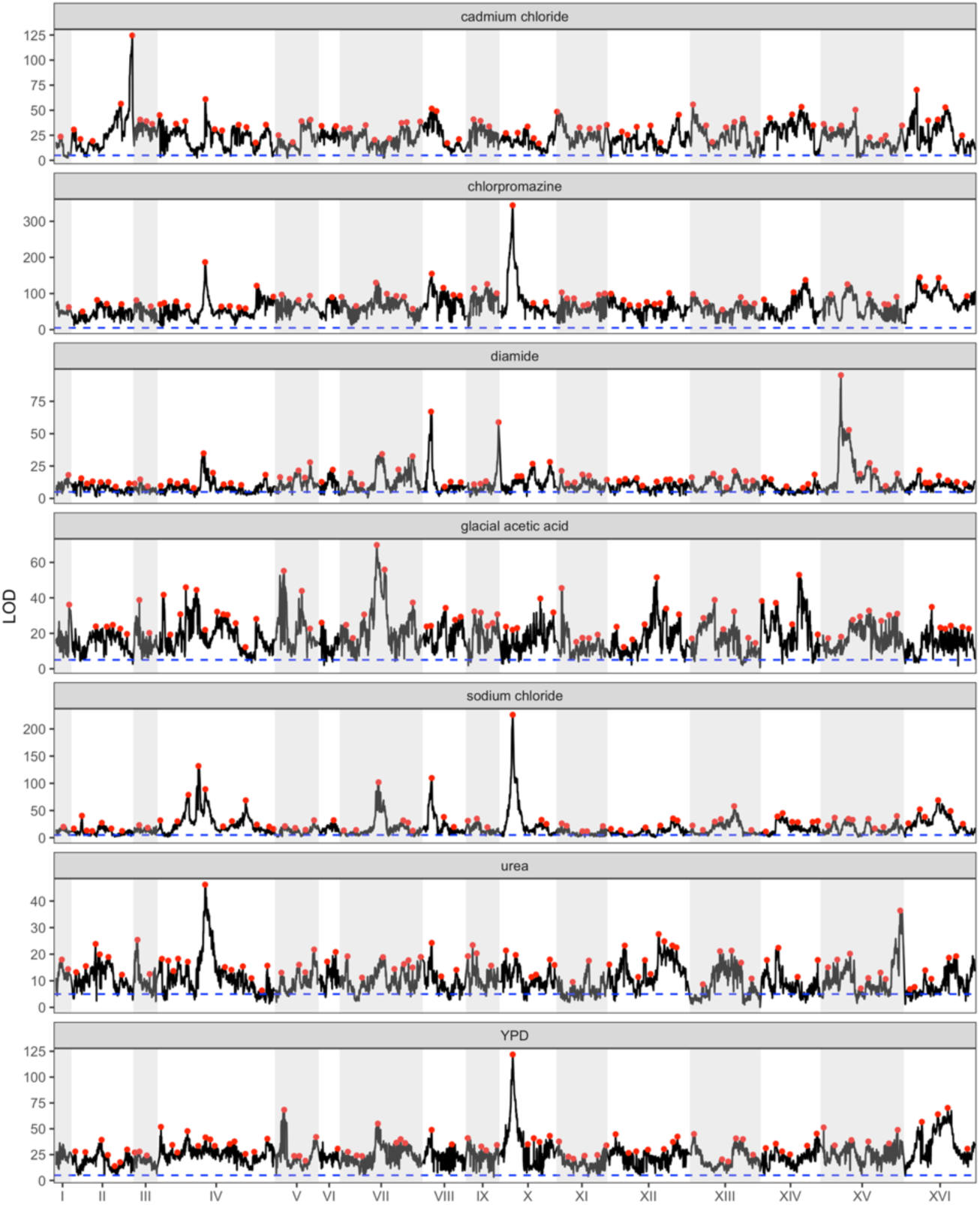
Genome-wide Manhattan plots of the LOD score for all chemicals. Red dots represent putative local peaks. The dashed blue horizontal line marks an LOD score of 5.

### Identifying candidate major effect genes

Despite the vast majority of the genome exhibiting haplotype change, there are clearly sub-regions of the genome not inconsistent with major genes (or gene regions) playing a role in adaptation. Due to the highly recombined and diverse nature of the base population, as well as the number of replicates and population sizes maintained throughout evolution, we are often able to resolve potentially major effect loci to a small number of genes. In the Supplemental Results, we describe candidate genes and/or potentially causative variants under the top three peaks from each condition that suggest targets for future functional validation. In several cases, there are strong *a priori* or *a posteriori* candidate genes contained within the detected interval. Unlike a typical E&R experiment in sexual outbreds, we can track changes in frequency at founder haplotypes and observe that adaptation is often due to the action of a single haplotype (below). As a result, candidate variants unique to that haplotype are strong candidate causative variants. Further, by virtue of the founders being *de novo* sequenced through both long-read and short-read technology (Linder et al., 2020), we are able to identify both candidate SNPs as well as candidate structural variants private to a particular founder. Figure 4 exemplifies a peak with a single haplotype increasing in frequency identified in cadmium chloride treatment. Panel A depicts the genome-wide LOD score, while panel B zooms in on chromosome XVI, and panel C shows the LOD scores and the sum of the squared haplotype change for a peak on this chromosome, as well as the genes under the peak, the absolute SNP differences, and the average haplotype change over all cadmium chloride replicates. A red box in panel C highlights a candidate causative SNP private to the most increased haplotype (MIH) that creates a binding site for Yap1p, a transcription factor known to be involved in cadmium chloride tolerance (Wemmie et al., 1994). Figures S8 and S9 show examples of peaks underlying chlorpromazine, diamide, sodium chloride, and urea tolerance.

**Figure 4.**
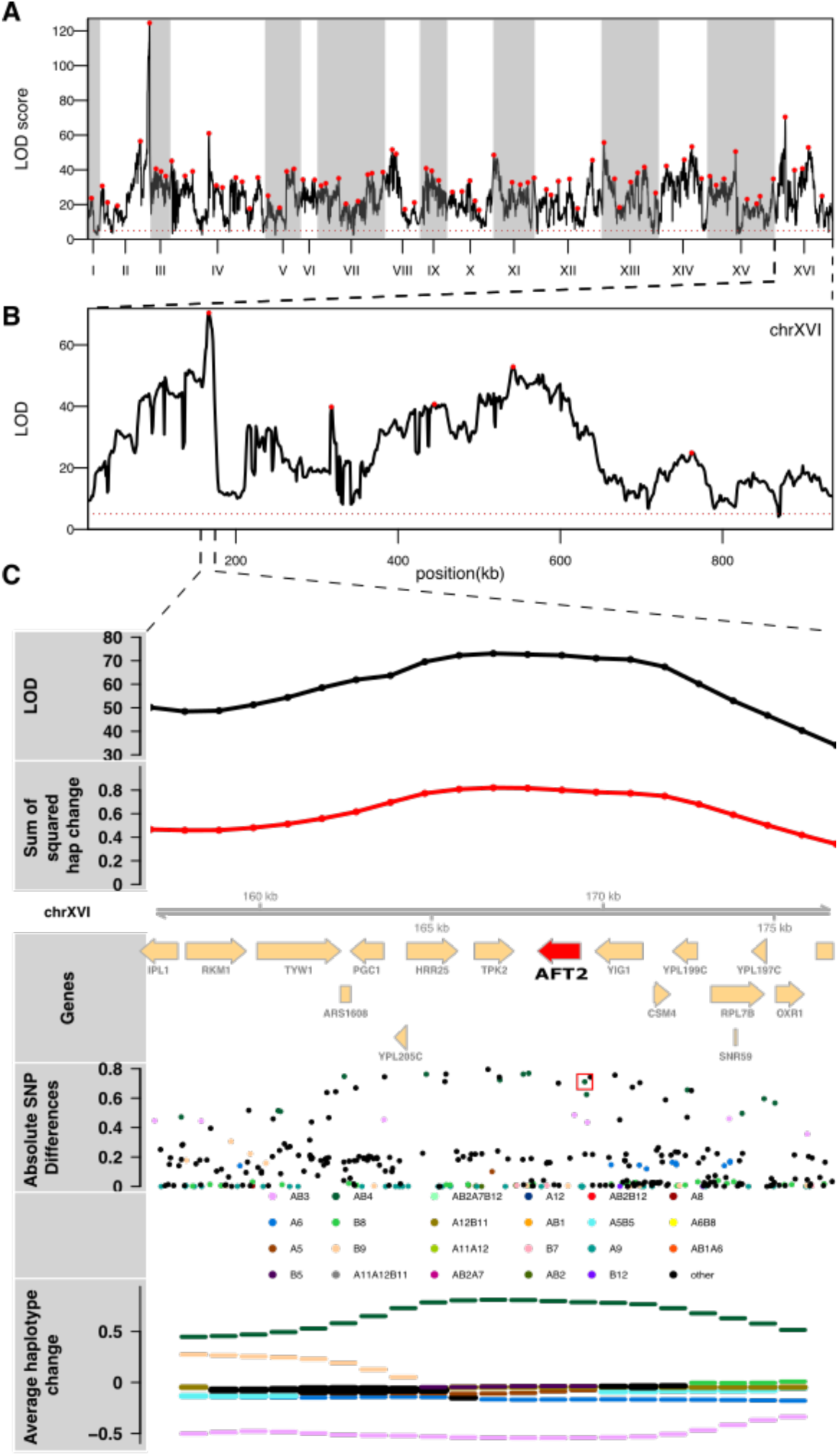
The results of a genome-wide scan for potentially causal genomic regions for cadmium chloride treatment is shown (panel A). Panel (B) depicts a close-up of chromosome XVI, where one of the highest peaks was detected and which emphasizes the polygenic nature of the adaptive response. Panel (C) zooms in under the peak on Chromosome XVI, where there is a candidate causative SNP (red box) just upstream of the candidate gene *AFT2* (highlighted in red) in the most changed haplotype, AB4, that creates a Yap1p binding site (see main text).

### Adaptation appears to be most often due to the action of a single haplotype

By virtue of our quantifying the dynamics of 18 founder haplotypes, we can identify the founder haplotype(s) showing the most evolutionary change over replicate evolved populations and correlate variation unique to that haplotype (or haplotypes) with the change. Figure 5, which compares the average change in frequency of the most increased haplotype (MIH) vs the next most increased haplotype per site across the 21 peaks discussed above, reveals that, across all chemical treatments, the most increased haplotype shows much larger gains than the next most increased haplotype. On average, the MIH increases by about 64.6%, while the next MIH increases by about 5.2%, a more than 12-fold difference. Furthermore, this disparity is consistent when comparing the MIH against the next MIH at each site across the genome, with the MIH increasing by an average of ∼23.6% and the next MIH increasing by an average of ∼9.4%, a ∼2.5 fold difference (Figure 5- figure supplement 1A). To show that this effect is not solely due to the initial frequency of the founder haplotypes in the population, Figure 5- figure supplement 1B shows the average change of the MIH vs the next MIH genome-wide only for sites at which the next MIH starts out at a higher initial frequency. The average increase of the MIH at these sites is ∼23%, while the average increase of the next MIH is ∼10%, a slightly more than 2-fold difference. These results show that adaptation across the conditions tested is primarily due to a single haplotype increasing in frequency, suggesting that adaptation is often either due to rare variants private to a single founder or due to a block of several mutations defining a local haplotype. There is little evidence that intermediate frequency SNPs, present on multiple haplotypes, individually drive adaptation. For intermediate frequency SNPs to be important in adaptation in this system, at a minimum, the idea of considerable traffic between closely linked factors needs to be invoked.

**Figure 5.**
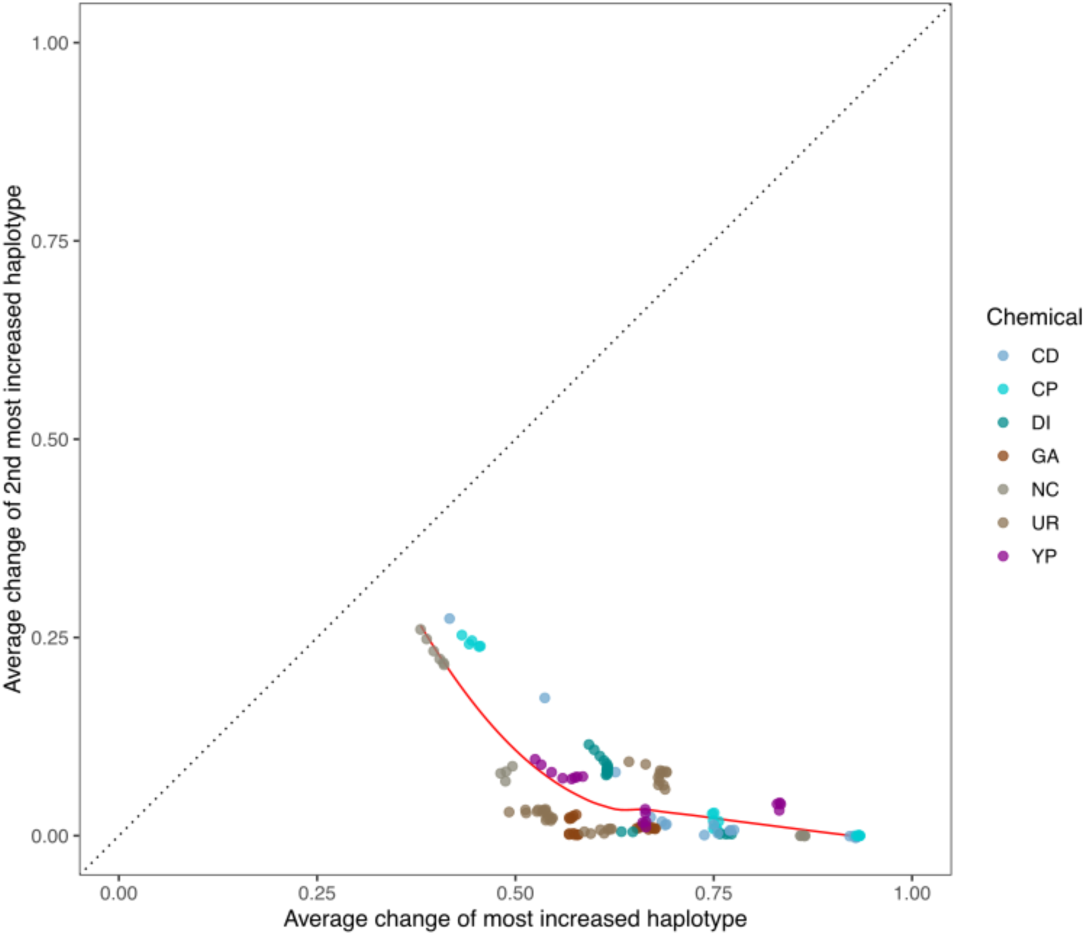
Average change of the most increased haplotype vs that of the next most increased haplotype at each of the 21 major peaks detected.

### The repeatability of evolution in a large-sized, outcrossed, sexual population

We determined the counts of minor alleles for all private SNPs in our base population as a proxy for the counts of all segregating sites (Figure 6- figure supplement 1). About 23% of private SNPs are not observed in the base population, suggesting they were either lost during the 12 rounds of intercrossing or present at an extremely low frequency. We also calculated the minor allele frequency (MAF) for all private SNPs in our base population present at a detectable frequency as a proxy for the MAF of all segregating sites and determined that the average MAF is ∼4.4%, with ∼79% of all MAFs greater than 0.4% and 100% of all MAFs at a frequency of greater than 0.04%. At these minor allele frequencies in the base population and given *N_e_*=750,000, all variants should be visible to selection and evolution should be essentially deterministic (given a primarily additive genetic architecture). To verify this, we calculated Spearman’s rho between haplotype frequencies for all pairs of replicates within a treatment at each locus in the genome. Figure 6- figure supplement 2 depicts the average such value for all pairs of populations within treatment and shows that the extent to which adaptive strategies are correlated is dependent on the specific chemical stressor employed. Figure 6A looks at the correlation in LOD scores between typical replicates across all loci and shows that regions of the genome showing more dramatic changes often share those changes across evolutionary replicates, with more subtle changes being less consistent. Chlorpromazine, sodium chloride, glacial acetic acid, YPD, and cadmium chloride all exhibit highly correlated genomic responses, while diamide and urea exhibit moderately correlated responses. To examine spatial patterns in correlated responses we transformed the LOD scores to z-scores (to somewhat mute the impact of major genes) for two groups of 3 different cadmium chloride replicates that typify responses and display those scores as a Manhattan plot (Figure 6B). The figure suggests that evolution is extremely replicable over much of the genome, with only a small number of regions showing noticeable differences in adaptive outcomes. A final metric that seems to indicate that the within treatment replicates are behaving similarly is to calculate the mean per-haplotype deviation at each position across the genome for each replicate. We then averaged these deviations over all replicates per chemical (Figure 7). The averaged deviations are similar in magnitude across all chemicals, for the most part falling between 0.05 to 0.1, again suggesting that changes in replicate haplotype frequency are similar across most of the genome; although a handful of local peaks can indicate regions of the genome where the responses were less coordinated.

**Figure 6.**
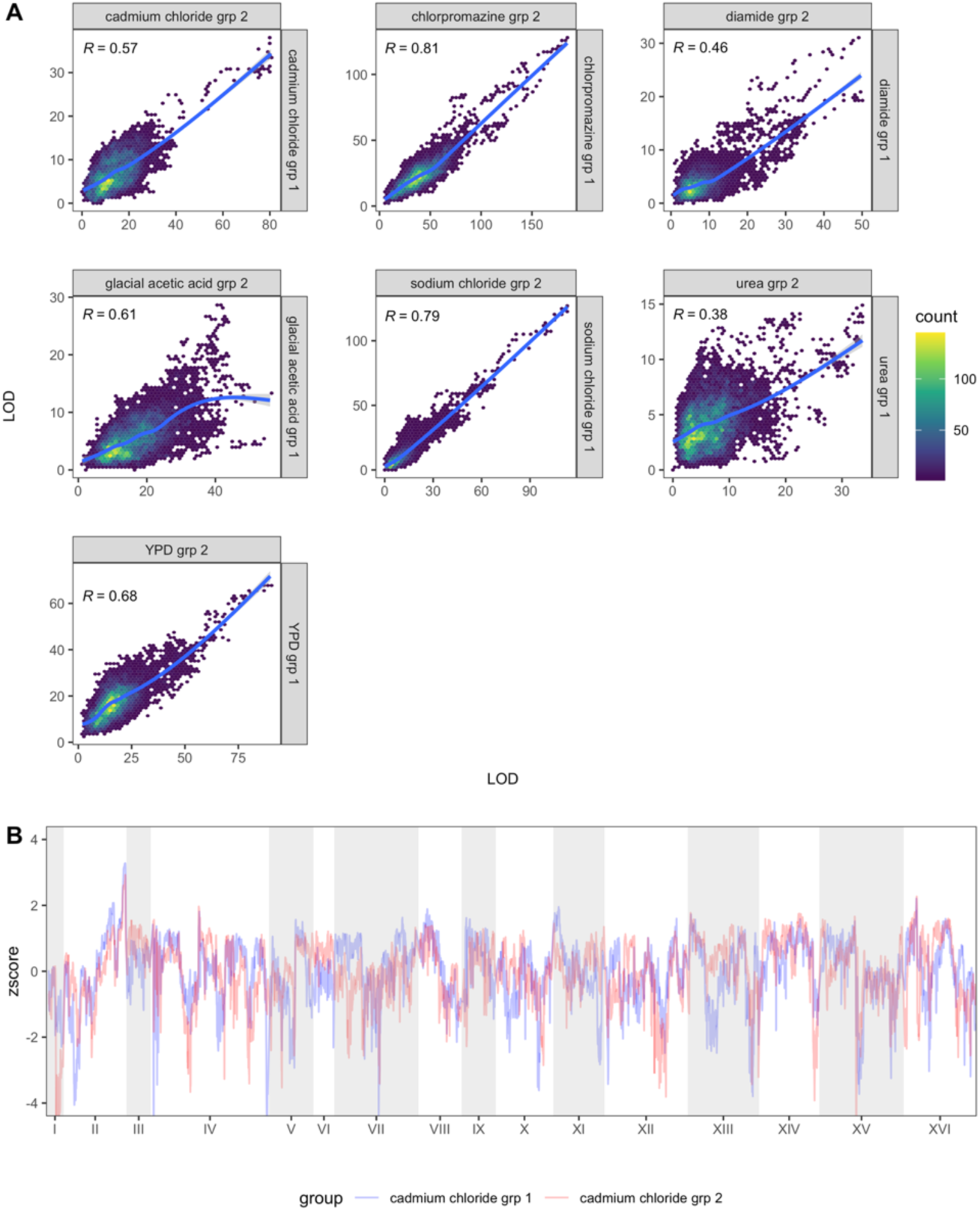
Repeatability of evolution amongst replicate populations. Panel (A) depicts the Spearman correlation between the LOD scores of randomly grouped replicates from each chemical treatment. Panel (B) shows the genome-wide z-score of each cadmium chloride group superimposed on one another.

**Figure 7.**
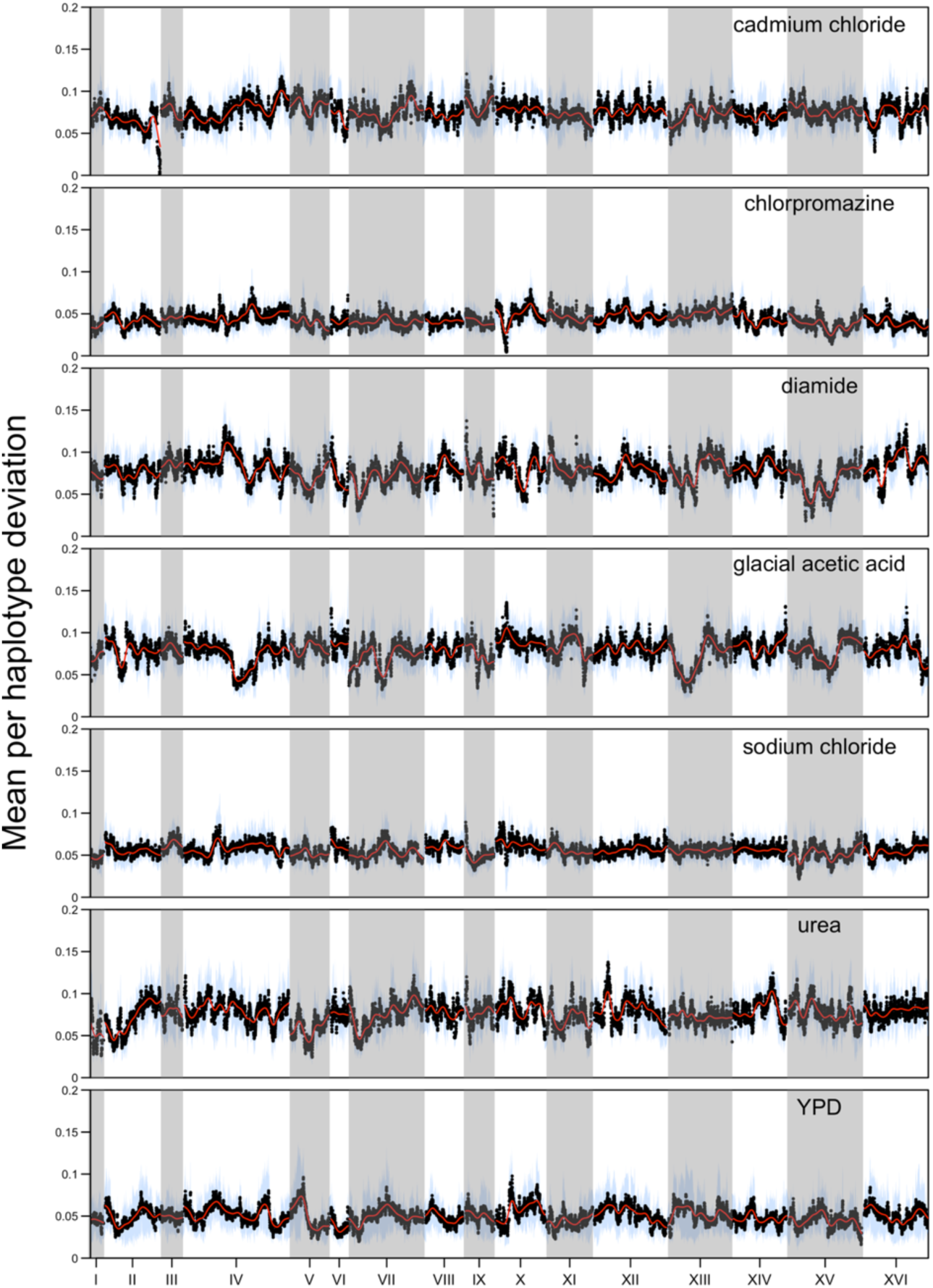
The mean per-haplotype deviation genome-wide averaged amongst replicates for each chemical. The red line represents a kernel regression run using the ksmooth() function in R with kernel set at ‘normal’ and bandwidth set at 100000. The blue lines represent one standard deviation from the mean.

A strong predictor of within treatment repeatability over drugs is heterozygosity, as average heterozygosity at 12 weeks is highly positively correlated with the average within treatment correlation in haplotype change (Figure 6- figure supplement 3). That is, evolution appears more deterministic in populations that maintain higher amounts of genetic variation and presumably experience softer selection over the course of evolution. Figure 6- figure supplement 4 shows that although populations maintain high levels of heterozygosity overall, heterozygosity does become depleted at regions that experience the largest amount of change across the different chemical treatments.

### The deterministic trajectory of evolution in large, outcrossed populations

Due to the large population sizes and relatively large amount of recombination experienced by our populations, we are in a unique position to examine how deterministic allele frequency changes are under the different selection regimens imposed. Figure 6- figure supplement 5A shows that, for the 21 leading factor peaks examined, there is no correlation between the initial frequency of the MIH and the average change in haplotype frequency, while Figure 6- figure supplement 5B shows that there is a strong relationship between the initial haplotype frequency and the average change in the next MIH. However, when looking at the entire genome (Figure 6- figure supplement 6A and B), there appears to be no relationship between the initial haplotype frequency and average haplotype change in both the MIH and the next MIH. This suggests that the selective pressure is high enough that evolution is highly deterministic across a wide range of initial frequencies in the base population.

### We observe some instances of strong pleiotropy, despite it being rare overall

By virtue of evolving our population to several different stressors with replication with all populations initiated at the same time from the same base population we can test for pleiotropy by identifying regions of the genome showing similar responses between treatments relative to pure within treatment replicates. Although an examination of the average LOD scores by chemical treatment (Figure 3) suggests the existence of several regions responding to multiple stressors, and hence exhibiting pleiotropy (discussed below), pleiotropy is not a general feature of the experiment. Figure 8 shows the correlation between average LOD scores for each pairwise combination of drugs.

**Figure 8.**
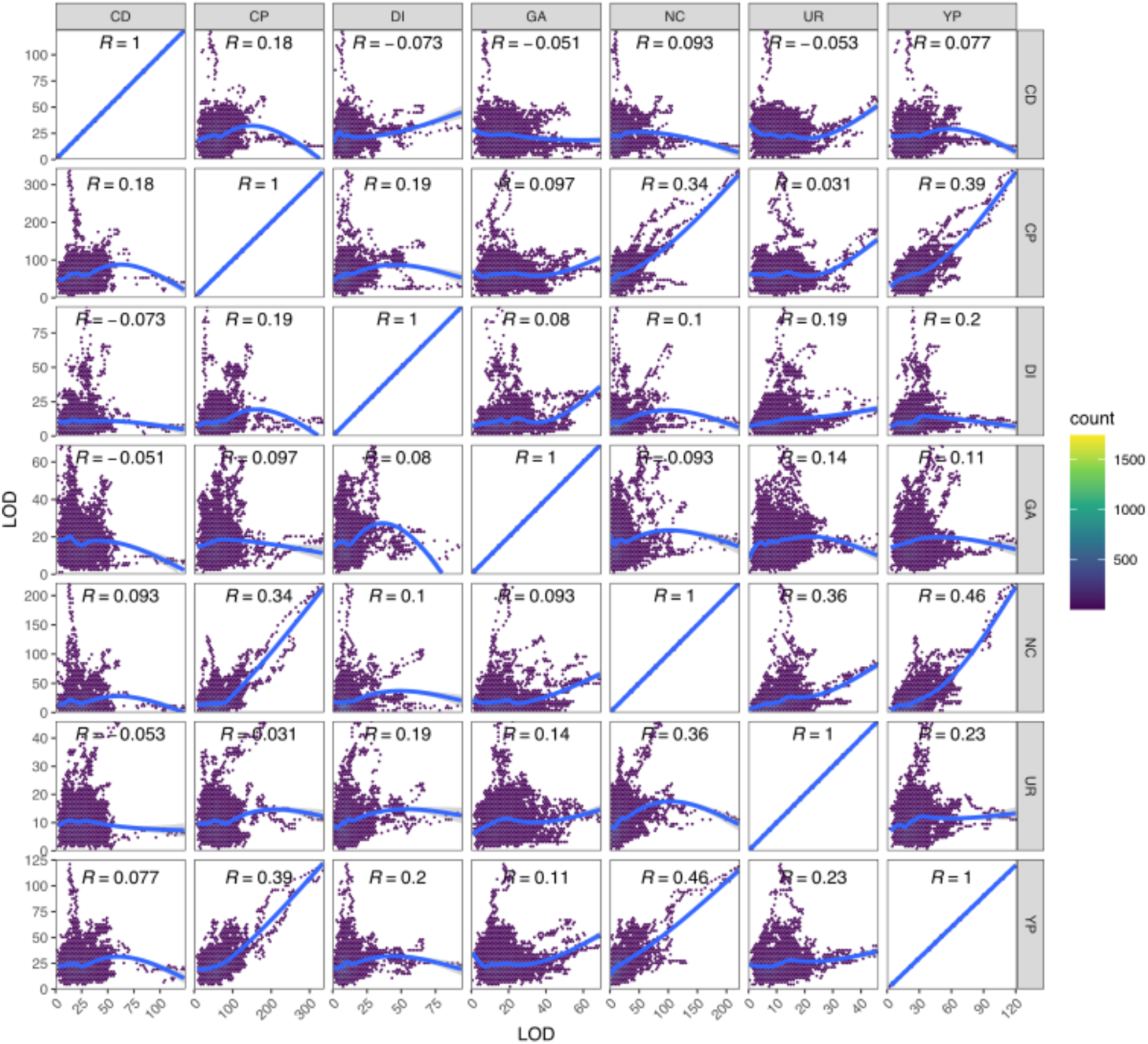
Detecting pleiotropy. Shown are the Spearman correlations of the genome-wide LOD scores for each pair of chemical treatments.

Although pure replicates are highly correlated in their evolutionary response (c.f. Figure 6A) this is not the case for between treatment comparisons in general, with most loci showing a response specific to the stressor they were exposed to. Several chemicals appear to elicit moderately correlated responses with one another, including YPD, sodium chloride, chlorpromazine, and urea, whereas cadmium chloride, diamide, and glacial acetic acid all exhibited more unique genome-wide patterns of selection. As detailed above, the parameters of this experiment enable us to identify relatively narrow intervals containing potentially pleiotropic gene regions, which are discussed in detail in the Supplemental Results. One striking example of pleiotropy occurs on chromosome X for a subset of three chemicals (chlorpromazine, sodium chloride, and YPD). All exhibit very similar evolutionary dynamics (Figure 9A and B) and a single founder (AB4) shows a large increase in frequency over the course of the experiment (Figure 9D). The peak change occurs at the three genes *TRK1, URA2,* and *PBS2* (Figure 9C), all three harbor SNPs private to AB4, and all are potential functional candidates based on annotation. It is notable that our base population is a *URA3* null (Linder et al., 2020), and perhaps *URA2* compensates for some subtle lost function in non-minimal media.

**Figure 9.**
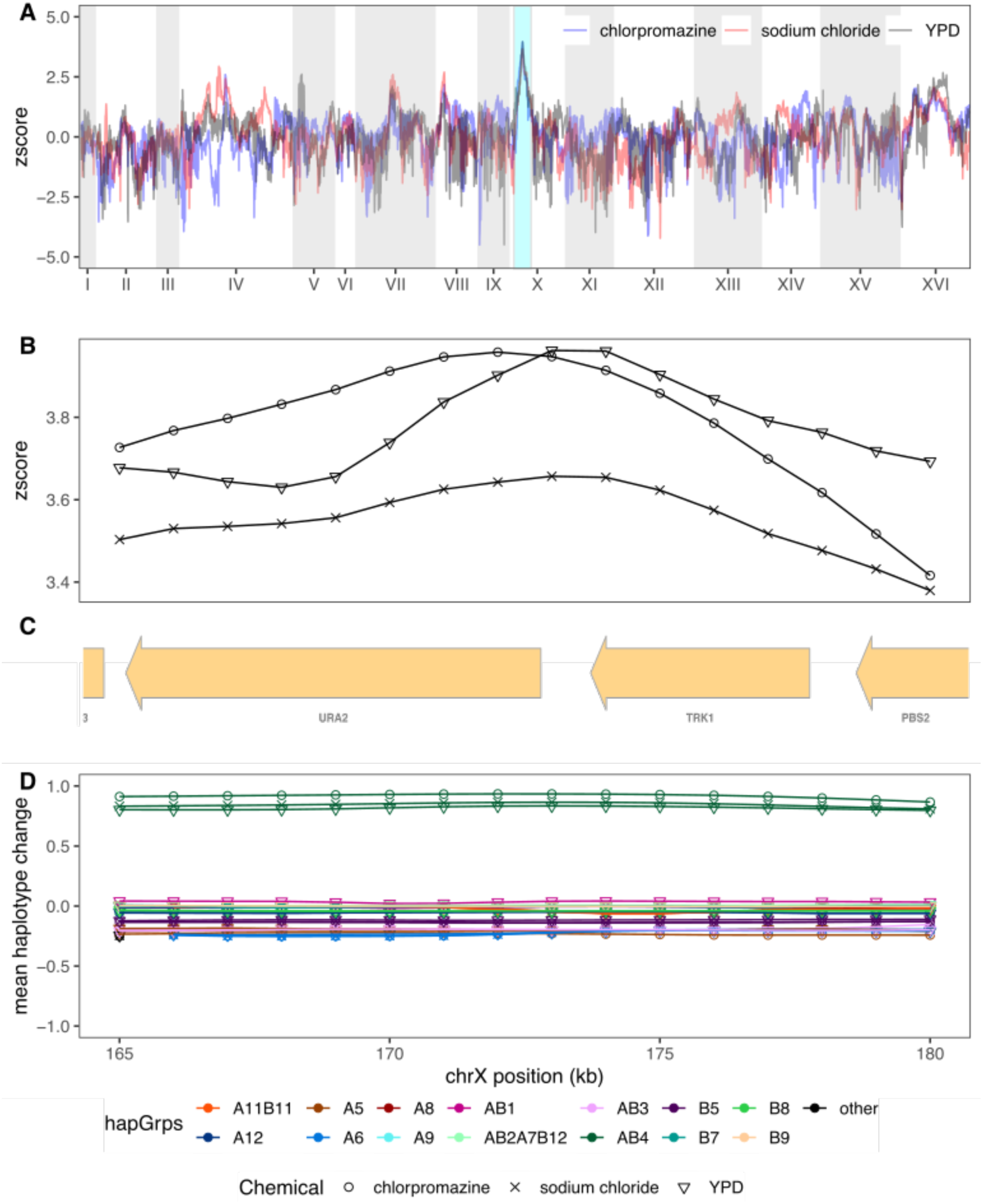
The most striking example of pleiotropy was detected on chromosome X. Panel (A) depicts the superimposed genome-wide z-scores of chlorpromazine, sodium chloride, and YPD, with the shared peak on Chromosome X highlighted in teal. Panel (B) zooms in on z-scores at the shared peak, with panel (C) showing the genes present in this region and panel (D) depicting the mean haplotype change amongst the three different chemicals.

## Discussion

### Yeast as a model for evolution in outbred sexual species

Evolve and Resequence (E&R) experiments have emerged as a paradigm for understanding evolutionary change. Under this paradigm, replicate populations are lab evolved and next generation sequenced, often as a pooled sample, to test evolutionary models. E&R experiments have been carried out in a variety of systems, including: 1. Evolving microbes asexually initiated from a single isogenic background (no sex, no variation), 2) evolving facultatively sexual microbes asexually starting from an outbred base population (no sex, with variation), 3) evolving facultatively sexual microbes with interspersed rounds of sex starting from a single isogenic background (sex, no variation), or 4) evolving either facultative or obligate sexual populations with sex starting from an outbred base population (sex, with variation). The last case of sex with variation is of special interest to evolutionary biology, since virtually all multicellular eukaryotes, including humans, fall into this category. Evolution in sexual outbreds has been commonly modeled using *Drosophila melanogaster*, but logistics dictate a modest number of replicate populations, tens as opposed to hundreds of generations of evolution, and effective population sizes (*N_e_*) generally less than one thousand individuals. In E&R experiments selected sites (or haplotypes) must have selection coefficients larger than ∼1/*N_e_* for selection to efficiently act on them. The result is that regions responding to selection in Drosophila E&R experiments are likely associated with selection coefficients of between 0.1 and 1%. Conversely, highly replicated large *N_e_* experiments can be carried out in microbes lacking sex and/or variation (Tenaillon et al., 2012), but evolution is then due entirely to newly arising mutations and clonal interference dominates evolutionary dynamics (Lang et al., 2013). It is not clear asexual systems effectively model evolution in outbred sexuals, and previous work has shown that adaptation can occur more efficiently in the presence of standing variation and recombination (Gray & Goddard, 2012; McDonald et al., 2016; Zeyl & Bell, 1997).

In a previous paper we manipulated yeast to create an outbred diploid highly recombined base population (Linder et al., 2020), and here we evolve that population under a set of conditions that mimic a burst of adaptation to a novel environment in a recombining diploid outbred population. We evolved populations at a large effective population size (*N_e_* = 750000) for 216 generations under a regime where sex is forced roughly once every 18 generations. Such a scheme results in a per gene per generation recombination rate ∼4X that achieved in *Drosophila*. The large effective population size approximates that of some natural populations and allows for intermediate frequency alleles with selection coefficients approaching 0.0001% to be visible to natural selection, and the high rate of recombination reduces the hitch-hiking footprint length associated with alleles changing in frequency. We finally adapted the yeast to several different chemical challenges in parallel. Despite some limitations of the system (discussed below), it is a powerful model for understanding evolution in large populations of diploid outbred eukaryotes.

### Limitations to using budding yeast as a model of obligate sexual evolution

Unlike obligate sexual species (such as *D. melanogaster* and humans), *S. cerevisiae* is a facultatively sexual organism, capable of reproducing both asexually and sexually. In nature, budding yeast predominantly divide asexually, with meiosis induced under harsh conditions, most likely to increase chances of survival (Ruderfer et al., 2006; Tsai et al., 2008). We manipulated yeast to create a large outbred diploid population which we forced to adapt to several environmental stressors, with adaptation occurring at large population sizes with frequent recombination. This artificial situation was designed to mimic adaptation in outbred sexuals, but clearly is not natural for yeast. Despite stringent selection for sporulation followed by random mating, ∼50% of our populations experienced the emergence and dominance of what appear to be asexual ‘cheaters’ (*i.e.*, individuals capable of making it through the evolution regimen without sporulating/mating). The propensity of cheaters to emerge and dominate a population correlated with the intensity of the selective pressure, with populations experiencing more intense selection being more likely to evolve cheaters. One potential explanation is that mutations of large effect are favored under these conditions, and the background in which such mutations occur can very quickly reach a high frequency in a population. Furthermore, the forced outcrossing procedure itself imparts a large fitness cost on the adapting populations, as evidenced by the nearly 600-fold drop in viable cell numbers from the beginning of sporulation to just after random mating. This substantial fitness cost associated with sporulation, spore isolation and dispersal, and mating results in strong selective pressure for individuals to evolve strategies that ‘game the system’. One potential means by which this could take place is through the construction and maintenance of fortified cell walls capable of withstanding the chemical and mechanical forces spores face during isolation and dispersal. It may even be possible that vegetative cells evolve that have the 4-layered cell walls of haploid spores without going through the process of meiosis. However, confirming the exact mechanism(s) of how asexual ‘cheaters’ survive is beyond the scope of this paper. Despite the emergence of cheaters, after aggressive filtering we were able to identify 55 replicate populations across seven chemical conditions that appear to have remained sexual throughout the experiment. These populations can serve as a model of evolution in sexual outbreds.

### Comparison to other long-term evolution experiments in facultatively sexual microbes

Some of the earliest experiments to document the advantages of recombination with standing variation used the facultatively sexual algae *Chlamydomonas reinhardtii (Colegrave et al., 2002; da Silva & Bell, 1996)*. These ground-breaking studies found that even a small amount of recombination can lead to long-term fitness gains compared to populations maintained purely asexually. However, only a few (up to 3) rounds of recombination were carried out and the molecular mechanisms behind the adaptive gains remain unknown. With the advent of next-generation sequencing technologies, the ability to resequence the genomes of evolved populations emerged as a powerful means to uncover the genetic changes underlying adaptation. Budding yeast have become a powerful model system to interrogate the genetic underpinnings of adaptation in the presence of sex and/or standing variation. Initial studies used highly outcrossed populations (that had gone through 12 rounds of recombination), generated from two or four founder strains (Cubillos et al., 2013, 2017; Li et al., 2019; Parts et al., 2011; Vázquez-García et al., 2017). Very large, replicated diploid populations were exposed to high temperature, hydroxyurea, or rapamycin and were propagated solely asexually for between 25-54 generations before whole genome resequencing of evolved and ancestral populations. These studies were pivotal in dissecting the role of standing variation on initial adaptation in outbred diploids but were carried out for a small number of generations without recombination during the selective regimen (no sex + variation). In contrast, two studies evolved large, replicate populations of budding yeast (on the order of ∼10^5^ cells, with between 3-4 replicates sequenced per condition) with regular rounds of recombination interspersed between periods of mitotic growth (Leu et al., 2020; McDonald et al., 2016). These studies spanned a much larger generational time (>= 1000 generations) with 11 or 24 rounds of recombination followed by random mating and showed that recombination alters the dynamics of adaptation but were initiated from an isogenic background without standing variation present during evolution (sex + no variation). At present, only two studies in budding yeast we are aware of have evolved populations with recombination in the presence of standing variation. One of these varied the frequency of recombination with replicate diploid populations of ∼10^5^ cells for nearly 1000 generations (Kosheleva & Desai, 2018), but evolution was only initiated from two genetically diverged founder strains. The other study took a 4-way population through 540 generations of mitotic evolution with 18 rounds of recombination (Burke et al., 2014) with populations maintained at ∼10^6^ cells. Both studies propagated budding yeast in rich media under a lab domestication regimen. Our study builds off these experiments by evolving populations with large effective population sizes with several replicates per condition for 216 mitotic generations interspersed with 12 rounds of recombination. Unlike previous studies in budding yeast, this study was initiated with much higher levels of standing variation, thus enabling us to better model long-term evolution in an outbred sexual species. And indeed, the higher levels of initial standing variation were associated with large fitness gains. Further, by evolving replicate populations in parallel to many different environments, we can make general inferences about the characteristics of adaptation. Finally, by virtue of the base population of this experiment being initiated from 18 different trackable founder alleles we are able to characterize the starting frequencies of variants important in adaptation.

### Comparison to other long-term evolution experiments in obligate sexuals

Most E&R studies to date have utilized *Drosophila spp.* to model evolution in sexual outbred diploids, with *Caenorhabditis elegans* and/or *remanei* perhaps emerging as an alternate model (Chelo & Teotónio, 2013; O’Connor et al., 2021). A large advantage of these studies is the high levels of genetic diversity segregating in the evolving populations. Drosophila E&R studies are often initiated from >100 founders, in stark contrast to sexual yeast populations initiated from four or fewer founders. Here, we employ 18 founders, still far fewer than fly experiments, but much more variation than previous budding yeast E&R experiments. The bulk of previous E&R work in *Drosophila* has been carried out for fewer than 100 generations (Barghi et al., 2017; Barghi et al., 2019; Franssen et al., 2015; Griffin et al., 2017; Hardy et al., 2018; Huang et al., 2014; Jalvingh et al., 2014; Jha et al., 2015; Kelly & Hughes, 2019; Martins et al., 2014; Michalak et al., 2017; Orozco-terWengel et al., 2012; Remolina et al., 2012; Shahrestani et al., 2021; Sørensen et al., 2020; Tobler et al., 2014; Turner & Miller, 2012), with a handful of studies carried out for between 100 to 200 generations (Hoedjes et al., 2019; Iranmehr et al., 2021; Kawecki et al., 2021; Turner et al., 2011; Zhou et al., 2011) and only a few studies maintained for between 400 to greater than 900 generations (Burke et al., 2010; Phillips et al., 2016, 2018). Minimum census population sizes in these studies have ranged from hundreds to greater than two thousand individuals (Shahrestani et al., 2021). In contrast our study maintained population sizes one to two orders of magnitude greater. The majority of studies in flies have evolved 2-5 replicate populations under 2-3 selection regimens, with the exception of a recent study in *D simulans* which used 10 replicates (Barghi et al., 2019) as well as a recent study in *D melanogaster* which used 6 replicates (Kawecki et al., 2021). The physical size of haplotype blocks changing in concert in fly E&R experiments varies among experiments and is difficult to estimate, but studies have suggested that such blocks can extend over several megabases (Barghi et al., 2019; Franssen et al., 2015), a significant fraction of the 120Mb Drosophila genome. The smaller size of haplotype blocks in our base population, with a median block size in the base population estimated at 66kb (Linder et al., 2020), along with higher rates of recombination per gene per generation, and an ability to track founder haplotypes allow the yeast system to detect adaptive genomic regions at a resolution much higher than in flies.

### Aneuploidies and de novo mutations are part of the adaptive response

Previous studies in budding yeast using populations with high levels of standing variation report conflicting results about the role of *de novo* mutations in adaptation, with two studies (both carried out with recombination) determining that mutations play a negligible role (Burke et al., 2014; Kosheleva & Desai, 2018), while another study showed that the role of mutations can depend strongly on the type of stressor used (Li et al., 2019). Of the 219 total *de novo* mutations detected in this study, only 20 reached a frequency >=50% and only three reached a frequency >=75%. This suggests that *de novo* mutations had a limited role in the adaptive response, which is further supported by the lack of multi-hit genes within a chemical treatment (only one gene was hit twice in a single condition). Aneuploidies have also been shown to have an important role in the adaptive response to a plethora of stressors, especially during the first few hundred generations of adaptation (reviewed in (Gilchrist & Stelkens, 2019)). We detect three different whole-chromosome duplications as well as ten unique segmental duplications, some shared across some replicated within drug treatments. These patterns suggest that aneuploidies can play an important role in adaptation, even in populations undergoing recombination. This is likely because some isolates of budding yeast are known to tolerate aneuploidies well (Hose et al., 2020) and reviewed in (Gilchrist & Stelkens, 2019). Further, the presence of aneuploidies does not seem to have affected our ability to map potentially causal loci, implying that recombination is still active in these regions. Aneuploidies do, however, seem to have a limited role in the adaptive response as all chromosomes showed extensive change in haplotype frequencies, and some replicates do not seem to harbor aneuploidies, yet have evolved.

### Long-term adaptation is highly polygenic

A large number of studies in flies have claimed that adaptation in outbred sexuals is highly polygenic, with many loci throughout the genome responding to selection (Barghi et al., 2017; Barghi et al., 2019; Burke et al., 2010; Chandler, 2014; Clark et al., 2007; Coop et al., 2009; Griffin et al., 2017; Hardy et al., 2018; Hoedjes et al., 2019; Huang et al., 2014; Jha et al., 2015; Kang et al., 2016; Kawecki et al., 2021; Orozco-terWengel et al., 2012; Phillips et al., 2016; Remolina et al., 2012; Shahrestani et al., 2021; Sørensen et al., 2020; Tobler et al., 2014; Turner et al., 2011; Turner & Miller, 2012; Tusso et al., 2021), but to some extent the inference of polygenicity is based on a visual inspection of Manhattan plots. *Very few studies attempt to estimate the minimum number of factors consistent with observed results*. Although hundreds of factors are consistent with observations, so are as few as 20-30 (Kelly & Hughes, 2019). Part of the problem is that E&R experiments start from limited numbers of haplotypes and selection coefficients are perhaps approaching 1%, so physically large haplotypes (>>1Mb) can hitch-hike with a selected site (Barghi et al., 2019; Franssen et al., 2015; Nuzhdin & Turner, 2013). The only other studies that evolved yeast with recombination that started from highly recombined populations are (Burke et al., 2014; Kosheleva & Desai, 2018) and both identified many regions of potentially small-effect throughout the genome, although major factors were also observed in (Burke et al., 2014). In this study, we find that almost the entire genome appears to respond to each of the seven different selective conditions, implying that most of the genome is either directly under selection or in linkage with regions under selection.

The size of unrecombined haplotype blocks centered on a selected site over a short evolutionary burst is proportional to the selection coefficient divided by the recombination rate per basepair per cell division (Kaplan et al., 1989). For a fixed selection coefficient, we thus predict the physical length of unrecombined hitchhiking blocks is ∼20X smaller in yeast than flies. Furthermore, since the effective population size is perhaps two orders of magnitude larger in yeast than flies, many additional factors with much smaller selection coefficients should be visible to selection, and those sites would be associated with smaller unrecombined blocks still (perhaps 200-2000X smaller). The large population size, high recombination rate, and observation of the entire genome responding to selection requires a much larger minimum number of factors to explain the apparent polygenic response in yeast relative to flies. This being said, more explicit simulations would be of value as the demography of the yeast evolution experiment involves phases of asexual growth punctuated by highly sexual episodes and there is some evidence that recombination occurs at hotspots in yeast (Illingworth et al., 2013). Either of these forces could result in larger than expected blocks and fewer factors required to explain the results.

### But also includes genes of large effect

Although there is a clear genome-wide polygenic response to selection, not all regions of the genome are responding equally, suggesting an important role for genes of large effect. We identify several such regions, many of which are associated with excellent candidate genes, and in many cases patterns of haplotype variation suggest candidate causative polymorphisms. Major factors challenge the dichotomy of the genetic architecture of adaptation being highly polygenic versus more oligo-genetic. Perhaps a more useful metric is to ask: how much of the observed change in fitness is due to the largest five factors (or what is the minimum number of factors required to explain half the fitness change)? This was the initial justification for QTL mapping, although thousands of segregating genes may impact a complex trait, it was believed that identifying the leading few was both possible and had practical value (Lander & Botstein, 1989; Thoday & Thompson, 1976). Unfortunately, we cannot estimate the relative contribution of these leading factors to total observed fitness gain. It is possible that high-throughput competition experiments employing thousands of barcoded gene replacements could be used to answer this question (Roy et al., 2018; Sadhu et al., 2018), although these experiments remain challenging if the factors being selected harbor multiple potentially interacting causative sites (see below) or if they ignore private alleles. The complex patterns of evolutionary change observed starting from relatively simple synthetic populations over short evolutionary bursts display complex enough dynamics that they are not trivial to model.

### Selection often seems to act predominantly on a single beneficial haplotype

E&R experiments are typically initiated from an outbred base population and then changes in the frequency of SNPs observed following adaptation (Barghi et al., 2017; Barghi et al., 2019; Burke et al., 2010; Franssen et al., 2015; Griffin et al., 2017; Hardy et al., 2018; Huang et al., 2014; Jalvingh et al., 2014; Jha et al., 2015; Kelly & Hughes, 2019; Martins et al., 2014; Michalak et al., 2017; Orozco-terWengel et al., 2012; Phillips et al., 2016, 2018; Remolina et al., 2012; Shahrestani et al., 2021; Sørensen et al., 2020; Tobler et al., 2014; Turner & Miller, 2012; Hoedjes et al., 2019; Iranmehr et al., 2021; Kawecki et al., 2021; Turner et al., 2011; Zhou et al., 2011). Such studies have several strengths, but it is difficult to distinguish SNPs that are drivers versus responders. Other E&R experiments have been initiated from a small number of founder alleles (Burke et al., 2014; Chandler, 2014; Cubillos et al., 2013; Li et al., 2019; Tusso et al., 2021), in these cases all selection is acting on common variants. In a 4-way cross from which asexual selection was carried out and founder haplotype frequencies tracked there was some evidence that the majority of adaptation was due to one founder versus three (Cubillos et al., 2013). This was a curious observation as it suggested selection was, for the most part, not acting on common variants. In this study, initiated from 18 founders, whose frequencies were tracked, the selective response is almost always associated with a single haplotype that is favored over all the others. This suggests that selection is almost never acting on evolutionary older intermediate frequency SNPs shared among several founders (or we would expect to see multiple founders increasing in frequency at similar rates). Instead, results suggest that selection is almost always acting on a multi-causative mutation haplotype or a rare allele private to a founder. Our observation that almost all adaptation is associated with single haplotypes is difficult to reconcile with the idea that evolution acts primarily on intermediate frequency causative alleles.

### Long-term evolution appears highly repeatable

We observed very high levels of repeatability across evolutionary replicates within drugs, with responses over drugs being different from one another. The two previous studies in budding yeast starting from an outbred base appeared highly repeatable ( Burke et al., 2014; Kosheleva & Desai, 2018) as have several studies from flies (Burke et al., 2010; Hoedjes et al., 2019; Iranmehr et al., 2021; Jalvingh et al., 2014; Jha et al., 2015; Kang et al., 2016; Kawecki et al., 2021; Kelly & Hughes, 2019; Martins et al., 2014; Michalak et al., 2017; Phillips et al., 2016, 2018; Remolina et al., 2012; Tobler et al., 2014; Turner & Miller, 2012; Zhou et al., 2011). Contradictory results have occasionally been reported in flies, in which the level of repeatability is much lower across replicate populations (Barghi et al., 2019; Griffin et al., 2017; Hardy et al., 2018). A classic result in population genetics is that once an allele reaches a frequency of greater than 2/N_e_s its dynamics are largely deterministic. As a result, causative alleles at frequencies in our base population greater than 3%, 0.3%, and 0.03% with selective coefficients as small as 0.01%, 0.1% and 1% are visible to natural selection and should behave somewhat deterministically in our populations (given N ∼750000). Since the majority of SNPs have frequencies in the yeast base population greater than 0.4% (and virtually all SNPs have frequencies greater than 0.04%) we expect evolution to be highly repeatable. In contrast, fly E&R experiments are often initiated from a large number of unknown founder haplotypes, perhaps 200-500, and effective population sizes are modest at <<1000 individuals. And only alleles present in multiple founder backgrounds have frequencies greater than 2/N_e_s and can act deterministically. A corollary is that rare variants present in fly E&R base populations are not visible to selection unless they have large selective coefficients or they drift or draft to a higher allele frequency. As a result it is difficult for adaptation to be due to rare alleles in Drosophila experiments, and experiments may appear unrepeatable if adaptation is often due to alleles with frequencies close to 2/N_e_s. It is unclear if repeatability is a prediction of a highly polygenic model; individual selection coefficients under such a model are << 0.1%, selection cannot distinguish individual causative sites, and selection is instead acting on haplotype blocks that happen to contain tens to hundreds of causative sites whose net effect is visible. Thus precise predictions of the polygenic model with respect to the repeatability of adaptation perhaps depend on the details of the distribution of selection coefficients and whether causative sites are clustered in the genome.

### Pleiotropy appears rare

In total, seven regions were detected as potentially pleiotropic, with four of these regions showing the same Most Increased Haplotype (MIH) across treatments. As only regions with LOD scores at least 1.96 standard deviations away from the mean were considered, all potentially pleiotropic sites likely have a large effect on fitness in the environments they were detected in. Theory predicts that such large-effect mutations should only affect a small number of phenotypes, as most such effects should be deleterious (Fisher, 1930; Orr, 2000), which would, at first glance, appear to run counter to what we see. In agreement with theory, previous work has shown that mutations that increase fitness in the environment they were detected in (the ‘home’ environment) can have similar effects on fitness in subtly different environments, but can have drastically different effects in different environments (Kinsler et al., 2020). This occurs because, although mutations do affect several phenotypes, only a small number of phenotypes are relevant to the condition mutations evolve in. However in rare instances, mutations could have similarly beneficial effects in very different environments as well. Another study in budding yeast has shown that although evolution tends to lead to specialization in the home environment, leading to smaller fitness gains or losses when individuals from that population are exposed to a very different environment, exceptions can occur (Jerison et al., 2020). Together, these results point to the conclusion that, while rare, alleles can have large beneficial effects in multiple different environments. This is consistent with what we observe, as only a handful of potentially large-effect sites have similarly large effects in different environments. Alternatively, overlapping aspects of the different environments used in this study may cause a phenotype important to adaptation in one environment to be important in other environments as well, leading to the same variant having similar effects across environments.

### The difficulty of validation of candidate variants

Validation of potentially causative candidate genes is beyond the scope of the current study. Gene knock-outs would not necessarily additionally implicate proposed candidate genes (since genes are often candidates due to their known loss of function phenotype), so more difficult allele replacement experiments are required. That evolution is primarily due to single rare haplotypes suggests that in many cases the candidate “allele” is not a single mutational event. Furthermore, showing that a single base change confers a selective advantage does not show that this change explains the observed haplotype change. To demonstrate this a single base change would need to be competed against alleles in which a larger region is substituted. Despite the high levels of recombination we attempt to maintain, we were unable to determine the length of haplotypes that are responding to selection. Validation via saturated replacement of all SNPs, singly and in combination, present on selected haplotype blocks is becoming possible using emerging high-throughput CRISPR/Cas9 methods (Akhmetov et al., 2018; Roy et al., 2018; Sadhu et al., 2018), although if selected haplotypes are associated with dozens of mutations (Pritchard, 2001; Thornton et al., 2013) these approaches may fail to explain haplotypic change. High throughput methods that compete entire regions in otherwise standard backgrounds hold some promise for future work.

## Conclusion

We manipulated yeast to serve as a model system for understanding evolution in outbred sexual diploid species harboring high levels of standing variation (*i.e.*, virtually all higher eukaryotes). We evolved replicate yeast populations in the presence of several chemical stressors for 216 generations with forced sex once every 18^th^ generation, at effective populations sizes approaching those of some natural populations. Like previous studies in Drosophila, we observe major genes important in selective response, with much of the genome responding to selection. Patterns of genetic change appear more repeatable than fly E&R experiments, are highly repeatable within drug treatments, but show little convergence across drugs (providing little evidence for universal pleiotropy). We estimate the rate of recombination per generation per individual per basepair (gene) to be approximately 20X (4X) higher in our yeast compared to fly E&R experiments. This greatly expanded recombinational map makes it increasingly difficult to explain genome-wide responses to selection as being due to a modest number of causative factors and traffic due to linkage disequilibrium. We make the novel observation that genome-wide haplotype change is almost always and replicably driven by only one of the 18 founder haplotypes, suggesting that selection is acting primarily on alleles private to single founders or multi-mutation haplotypes, and rarely intermediate frequency SNPs present on multiple founder haplotypes. The observation that selection almost always acts on a single haplotype likely rules out some polygenic models for the architecture of adaptation, although current models need to be more explicit and capable of making precise predictions.

## Materials and Methods

### Strains and media

The base population used in this study is a multi-parent population (MPP) derived from a full diallel cross of 18 highly characterized natural haploid founding strains (Cubillos et al., 2009) intercrossed for 12 generations to break up haplotype blocks. This population, 18F12v2, and the 18 founding strains are described in detail elsewhere (Linder et al., 2020). Non-selective propagation of yeast was carried out using rich YPD media (1% yeast extract, 2% peptone, and 2% dextrose (Fisher, Waltham, MA)) with 2% agar added for solid media. In some cases the YPD was supplemented with 100mg/mL ampicillin (YPDamp) or 200mg/mL of kanamycin (YPDk) to prevent bacterial contamination. During long-term evolution the YPDk media was spiked with 2 different doses of each chemical (a day 1 of the entire experiment dose and a day 2 onwards dose; see Table S1). Sporulation media consisted of 1% potassium acetate and a 1X dilution of a 10X amino acid stock (composed of 3.7g of CSM -lysine (Sunrise Scientific, Knoxville, TN) supplemented w/10mL of 10mg/mL lysine in 1L total volume), pH adjusted to 7 (‘PA7’). Mating media consisted of YPDamp. Spore isolation solution (‘SIS’) consisted of 25U zymolyase 100T (Amsbio, Abingdon, United Kingdom) + 10mM DTT (dithiothreitol; Thermo, Waltham, MA) + 50mM EDTA (ethylenediaminetetraacetic, Fisher) + 100mM Tris-HCl, pH 7.2 up to 500uL. Spore dispersal solution (‘SDS’) consisted of 20U zymolyase 100T + 800ug lysozyme (Sigma, St. Louis, MO) + 1% Triton X-100 + 2% dextrose + 100mM PBS, pH 7.2 up to 400uL. Double selection media (‘DSM’) consisted of 200ug/mL nourseothricin sulfate (‘clonNAT’) + 600ug/mL hygromycin b + 100ug/mL ampicillin in YPD. Concentrations of cloNAT and hygromycin b are twice as high as normal as DSM was added in equal volume to the media used for mating (i.e. diluted 2x).

### Long-term evolution of diploid budding yeast

The long-term evolution experiment was initiated from 6mL of 18F12v2 (hereafter the base population) thawed at RT, spun down, and resuspended in 20mL of YPDamp, split into two 50mL erlenmeyer flasks (covered) and incubated at 30°C at 275 RPM for 3h to ensure cells had entered exponential growth. Cells were harvested in YPD, counted using a haemocytometer, and 140uL of culture containing ∼1.4 million cells was transferred into shallow 24-well plates (Phenix, Candler, NC; TCG-662160) containing 1.4mL YPDk supplemented with 15 different chemicals (as well as YPDk-only controls). These 24-well plates allow for a 1.5mL working volume with shaking with minimal cross-contamination. In total, 16 replicate populations were spiked into each of the 15 chemicals, along with a total of 10 YPDk-only replicate control populations. Wells containing only YPDk (no cells) were scattered across plates to monitor for signs of cross-contamination (6 negative control wells total). The total experiment thus consisted of eleven 24 well plates.

For transferring cultures to new media, the collection of 16 different YPDk + chemical plates were made in bulk and stored at -80°C for up to 6 months to minimize batch effects. Plates were thawed at 30°C for ∼2h just prior to being used. Tables 1 and S1 list the chemicals used and their known effects in budding yeast, as well as the doses used in this experiment. Chemicals used included: chlorpromazine, cisplatin, fluconazole (all from Cayman, Ann Arbor, MI), diamide, sodium sulfite, dmso, urea (all from Fisher), ethanol, sodium chloride, nicotine (all from VWR, Radnor, PA), cadmium chloride, nicotinamide (both from Sigma), tunicamycin (Enzo Life Sciences, Farmingdale, NY), glacial acetic acid (J.T. Baker, Phillipsburg, NJ), and caffeine (ICN Biomedicals, Costa Mesa, CA).

The schematic of the long-term evolution regimen is shown in Figure 1. On the first Tuesday of the entire experiment only, a lower dose of most chemicals was used to acclimate cells to the chemical challenge (to allow cells to physiologically adapt). For the remainder of the experiment, an increased dose was used to maintain a high degree of selective pressure. On Tuesdays, Wednesdays, and Thursdays, 10x serial dilutions of cells were transferred to newly thawed chemical plates using a custom-built liquid-handling robot using 1000uL Tecan Freedom EVO Tips (Phenix, TRXF-HTR1000CS, filtered). The robot is contained in an acrylic cover with a large HEPA-filtered blower placed on top to maintain laminar flow and ensure that cultures could be transferred aseptically. Plates were covered in adhesive membranes (VWR, 60941-086) and transferred to a plate-holding device that was placed into a shaker-incubator. Cultures were incubated at 30°C for 24h at 175 RPM. During plate transfers, it is of note that the adhesive membranes were pierced directly by the robot without need for removing the membranes beforehand to minimize the chances of cross-contamination. In addition, every Thursday, the OD630 of all cultures were read in a shallow, clear 96-well plate (Phenix, MPA3370), and glycerol stocks (an equal volume of 40% glycerol added to the culture) were created for three copies of all cultures and stored at -80°C to maintain a fossil record of evolution.

On Fridays, cells were transferred to deep 24-well plates (Phenix, MP-2061), spun down, washed once with 1mL of sterile milli Q water (smq H2O), then resuspended in 1mL of sporulation media. Cultures were incubated for 3d (over the weekend) at 30°C at 200 RPM. All spin downs of plates with adhesive membranes were carried out at 1500 RPM, as faster speeds caused membranes to tear. On Mondays, a random subset of cultures were examined to ensure at least 50% of the culture had sporulated, after which the deep 24-well plates were spun down and pellets resuspended in 500uL of SIS. Cultures were transferred to a deep 96-well plate (Phenix, M-0564), sealed with a plate seal, vortexed on a plate vortexer for 5m at 1200 RPM, then incubated at 30°C for 1h at 200 RPM to spheroplast cells. Cultures were then spun down and resuspended in 1% Tween 20 to lyse unsporulated cells. Cultures were again spun down, resuspended in 400uL of SDS, and transferred to another deep 96-well plate loaded with 300uL of 400um silica beads. Plates were sealed (Phenix, SMX-DW96), tetrads were disrupted and sister spores dispersed by shaking in the GenoGrinder 2000 at a setting of 1500 strokes per minute for 20m. To transfer disrupted spores to a new plate we used a chimney PCR plate (LightLabs, Aurora, CO; A-3004-C) with holes burned into the base using a soldering iron- the top of the chimney plate was forced into the deep 96-well plate and the base inserted into a second deep 96-well plate (Phenix, MDZ96-22SV) and spun until the centrifuge reached 1500 RPM. This process was then repeated again to minimize the carry-over of beads. Cells were pelleted, washed in YPDamp, resuspended in 250uL of YPDamp, and allowed to mate for 3.5h at 30°C at 25 RPM. 250uL of DSM was then added to all cultures to select for successfully mated diploids and cells were incubated overnight at 175 RPM at 30°C. On Tuesday, 140uL of cells were transferred into chemical plates to begin the cycle again. In total, evolution was carried out for 12 weeks, with an estimated 18 mitotic and 1 meiotic generation occurring each week for a total of 216 mitotic and 12 meiotic generations.

### Revival of frozen stock to resume evolution

Periodically, evolution would need to be ‘rebooted’ from a previously archived week. This occurred due to technical issues with our instruments or other catastrophic cross-contamination events. To reboot evolution, glycerol stock plates were taken from the - 80°C freezer on a Monday, spun down briefly (up to 1500 RPM), placed on the robot deck, and allowed to thaw completely at RT. Cultures were then transferred to deep 96- well plates, spun down, washed with 330uL of smq H2O, spun down again, then resuspended in 500uL of YPDamp and incubated O/N at 175 RPM at 30°C to allow the populations to recover. Cultures were then serial transferred into their respective chemical stressor in shallow 24-well plates for 3 days before sporulating on Friday. As the glycerol stocks were made on a Thursday, a reboot skips a meiosis cycle. *This strategy was necessitated as we observed that the rate of sporulation increased the longer we allowed cultures to divide mitotically with serial dilutions after being revived from a glycerol stock*. We have not seen this claim elsewhere in the literature, but it is a very replicable result.

### Whole genome sequencing of the evolved populations

Archived week 12 evolved populations were thawed and incubated in deep 96-well plates in 960uL of fresh YPDamp O/N at 30°C at 175 RPM. Cells were then harvested, washed once with buffer TE, and genomic DNA extracted using a modified 96-well DNA extraction protocol. Spheroplasting was carried out at 37°C for 30m with zymolyase-containing buffer, followed by incubation in lysis buffer and proteinase K at 50°C for 60m (68°C for 10m at the end to kill the proteinase activity). Cell lysates were then incubated in 5M potassium acetate at 4°C for 30m. An additional 100uL of sigma water was added and samples were spun down and the supernatant transferred to a clean 96- well PCR plate. An additional centrifugation step was carried out, followed by transferring the DNA-containing supernatant to an equal volume of RT 100% isopropanol. Samples were then spun down, washed with RT 80% ethanol, spun down again, air dried, and finally purified DNA was resuspended in 50uL of warm buffer EB. DNA was quantified using the Qubit 1.0, followed by preparing the samples for Illumina sequencing using the Nextera Flex kit with modifications. Reactions were carried out at 1/5^th^ of the normal volume as detailed in our previous study (Linder et al., 2020). Libraries were sequenced on the HiSeq4000 with PE100 reads. Coverage per sample ranged from 5x to 145x, with a mean coverage of 54x (Figure 1- figure supplement 2).

### Estimation of daily bottlenecks during a week of the long-term evolution regimen

To estimate bottlenecks and the number of divisions experienced by our populations during a typical week of evolution, a single sexual (see below) YPD population was sampled daily, starting Monday and ending Friday during week 11 of evolution (YPD replicate R10 - see Figure 1- figure supplement 1A/B). Sampling consisted of making serial dilutions in 0.1M PBS, pH 7.2, vortexing the dilutions vigorously, and plating out the dilution series onto YPDA plates shaken with beads to inhibit the clumping of cells. Plates were incubated for 2-3d at 30°C and imaged using a geldoc imager with the lids off. Colonies were automatically counted using ImageJ as detailed in (Stolze et al., 2019). The adjustable watershed add-on was used with tolerance set to 0.1. Particle size was set to 0.015” and circularity to 0.3. On Monday, samples were taken at three time-points (Figure 1- figure supplement 1C): a) post 3-days of sporulation while cells were still in sporulation media before any processing had occurred (with colonies measuring the number of viable tetrads), b) after spores had been dispersed and isolated, but just before mating (with colonies measuring the number of viable spores), and c) after 3.5-hours of mating before the addition of DSM or cell divisions would have occurred (measuring the number of viable diploid cells). Not surprisingly the maximal amount of bottle-necking experienced by the populations over the course of the week is observed in counts of diploid cells immediately after mating, as this time-point integrates over several previous steps in which spores are killed (due to breaking the ascus, vigorous dispersion, and/or vegetative cell killing steps) and/or are unable to mate or result in inviable diploids. This weekly nadir in census population size is important, as it is closely related to the effective population size we were able to maintain during our experiment. We estimate that through much of the weekly cycle we maintain tens to hundreds of millions of cells, with a single weekly bottleneck in the high hundreds of thousands of diploid individuals.

### Tracking cultures through time via weekly OD630 measurements

To keep track of which populations went extinct over the course of the experiment, OD630 measurements were taken every Thursday of all cultures in a clear, shallow 96- well plate. These weekly measurements consisted of using the liquid-handling robot to transfer 140uL from each shallow 24-well plate with cultures incubated between 23-25h (at the same time as serial transfers to fresh chemical plates were carried out). As a side-effect of measuring the total number of cells through a typical week of evolution (described above), we are able to estimate that cultures likely had not reached saturation at this point as saturated cultures of *S. cerevisiae* normally reach concentrations of over 5×10^8^ cells/mL (Chan et al., 2013). Thus, these OD630 measurements were taken while cultures were still growing and could be averaged across replicates and across time-points (to reduce the noisiness of the data) to correlate OD630 with the average number of well-behaved, outbred sexual populations that evolved from a chemical treatment.

### Estimation of fitness gains after 12 weeks of evolution

To estimate the fitness gains of week 12 populations, a single evolved week 12 replicate population from each chemical was thawed, (along with the ancestral, week 0 population) transferred to a 96 deep-well plate and recovered O/N in 700uL of YPD supplemented with 100ug/mL ampicillin. To reacclimate populations to their respective chemicals of interest, populations were cultured in conditions nearly identical to the evolutionary regimen (the ancestral population was passaged only in YPD). Briefly, serial transfers of 140uL from each culture were made into a 24-well plate with the day 1 dose of their respective chemicals, followed by 24h of incubation at 30°C at 175 RPM, serial transfer of 140uL to the day 2+ dose of their respective chemicals, incubation at 30°C at 175 RPM for another day, followed by serial transfer of 50-200uL of culture to one (for our initial attempt) or eight (for our second attempt) fresh day 2+ chemical plates, which were incubated over a 24h period at 30°C at 175 RPM. Initially, we tried to measure the OD630 of a single plate over the 24h period with continuous shaking at 30°C in a plate reader, but discovered that several populations tended to form large clumps on the bottom of the wells, causing sudden spikes and dips in OD630 readings and making it impossible to derive accurate growth curves from this data. Therefore, we repeated the experiment as described above, except that during the final day of incubation, plates were removed at several time-points over the 24h period, including: 2.5h, 8h, 9.5h, 12h, 14h, 16h, and 24h, to estimate the growth rate of each of the week 12 populations and the ancestral population in YPD. The entirety of each well was transferred to a 96 deep-well plate, spun down, supernatant was removed, and pellets were stored at 4°C overnight. The next day, all pellets were resuspended in 0.1M PBS, pH 7.2 and kept at 4°C. Samples were selected for further analysis from cultures that had not yet reached saturation but were likely no longer in lag phase, including the 2.5h time-point to be used as the baseline and one or two later time-points. Samples were sonicated briefly on ice to break up clumps in 0.1M PBS, pH 7.2 containing 0.015% NP40 (settings were 30% of maximum amplitude- about 4-6 volts, for 20s, continuous sonication using a Sonics Vibra cell sonicator). Serial dilutions of cells were plated onto YPDA, incubated for 3d at 30°C, then colonies were counted either manually when possible or using ImageJ with the same parameters as described above. To estimate fitness in the base population, we assumed cultures initially barely survive the daily 10X dilution (with the exception of the YPD-only condition) at the start of the experiment, as our initial doses of each chemical were chosen to be close to the maximum possible tolerated whilst avoiding extinction. We also observe early generation cultures decreasing in viable cell counts over time, but bouncing back by the end of the first week of evolution. Under this assumption, the population growth rate per hour for week zero, derived from the equation 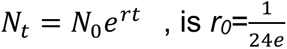. The growth rate at week 12 can similarly be estimated as 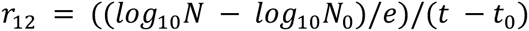, which is the same as the slope from the regression of log10 counts on time in hours divided by *e*. From this, the change in population growth rate (as a proxy for fitness gain) is *r*_12_/*r*_0_= 24 ∗slope estimate obtained from the 12-week evolved population. For estimating change in population growth rate for the YPD-only condition, the growth rate at week 0 (*r*_0_) was estimated in the same way, using the same time-points, as for week 12.

### Haplotype calling in Illumina sequenced evolved populations

A custom in-house haplotype caller was used to impute sliding window founder haplotype frequencies genome-wide, as described in (Linder et al., 2020). Briefly, we slide through the genome in 1kb steps, considering a 60kb window for each step. Due to the repetitive nature of the ends of chromosomes (Blackburn & Gall, 1978), accurate haplotype frequency estimates are difficult to obtain, and so these regions are excluded. In total, we estimate haplotype frequencies at 11,574 loci spaced every 1kb throughout the genome. The 60kb window was decided by trial and error in our previous work and results in highly accurate haplotype frequency estimates with absolute per haplotype frequency errors of ∼0.01 at the coverages employed in this study (Linder et al., 2020). The decision to use a 1kb step size balanced resolution with computational speed- at a 1kb step size the average absolute change in haplotype frequency between adjacent positions in founder AB3 is ∼5.6×10^-7^. Figure 2- figure supplement 1 shows the distribution of founder AB3 haplotype changes between all adjacent intervals estimated for the 55 populations classified as outbred sexuals and that clustered together from the seven chemicals used in downstream analyses (see below).

Only samples with genome-wide coverage greater than or equal to 5x were used for downstream analyses. Due to the unique mosaic-like genome structure of many natural yeast isolates (Liti et al., 2009; Peter et al., 2018; Schacherer et al., 2009) there are many tens to hundreds of kilobase sized regions of the genome over which two or more of the 18 founder haplotypes cannot be distinguished from one another. In the case where two (or more) founder haplotypes cannot be distinguished from one another we are unable to estimate the frequency of those two founders for that region, but the sum of their frequencies is estimated as well as any other haplotype. In our previous work we thus estimated the frequency of *f* founders that cannot be distinguished for some window as the sum of their frequencies divided by *f*. When looking at regions important in adaptation, in some cases, this averaging approach is misleading, so here we took the different approach of defining new “synthetic founders”. For example, for a region at which founders A11 and A12 cannot be distinguished, we create a founder called ‘A11A12’ for that window and track its frequency. This results in a more accurate picture of how regions are evolving, at the price of tracking a variable number of founder haplotypes for each region of the genome. We found that plotting the haplotype frequencies for the 23 most common haplotype combinations over all replicate populations and all chemical treatments (and an appropriate color palette) allowed us to visualize evolutionary change more easily. Over all windows, replicates, and treatments the 23 most common synthetic haplotype combinations account for an average of 94.7% of the observed haplotype frequencies.

### Peak calling

We created a Chi-square test statistic to identify regions of the genome most changed due to selection within any given chemical treatment. At any given position we identified the *K* haplotypes (including possibly synthetic haplotypes) with a starting frequency of at least 1% in the base population. At that position for the *R* replicate evolved populations within the *k*^th^ haplotype we calculate a normalized deviate from the expected change in allele frequency as

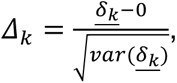

Where *<u>δ_k_</u>* is the average change in arcsin square root transformed haplotype frequency between the base and the R evolved replicate populations. Since *Δ_k_* is distributed as a unit normal, the sum of 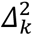 over the K haplotypes is distributed as a Chi-squared with *k* degrees of freedom. Although *var(<u>δ_k_</u>)* is unknown, we assume it is constant over haplotypes and estimate it as:

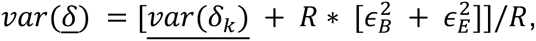

where ϵ*_B/E_* is the average error in the haplotype frequency estimates in the <u>B</u>ase or <u>E</u>volved populations, which we estimate as 0.004 or 0.01 respectively (Linder et al., 2020). That is, the average variance between experimental replicates plus the variance due to haplotype estimation error in the base and evolved populations. The different haplotype frequency errors are due to their being estimated from >2000x coverage sequencing in the base population as opposed to an average 53x in the evolved populations (with a range of 9x to 145x, Figure 1- figure supplement 2). We show in (Linder et al., 2020) that the error in haplotype frequency estimation is not strongly affected by sequence coverage, consistent with other such reports in the literature (Tilk et al., 2019). Since much of the genome for all chemicals is above the nominal significance threshold suggested by the Chi-square test, we attempted to call local peaks using an algorithm that finds local minima and maxima in a vector with an adjustable threshold (Friedland, 2017).

### Per-site heterozygosity deviation

The per-site haplotype heterozygosity was calculated at each of the 11,574 loci for the base and each evolved population as one minus the sum of the squared haplotype frequencies. We then calculated the average reduction in per-site heterozygosity by summing the differences in heterozygosity per-site between the base and each evolved population and dividing by the number of loci (11,574).

### Per-haplotype deviation

The genome-wide per-haplotype frequency deviation for replicate *i* was calculated for the frequency of the *h^th^* haplotype in that replicate (*f_h,i_*) at each position as the absolute difference between *f_h,i_* and *f_h,j ≠ i_*, which was simply averaged over the *h* haplotypes to obtain an overall deviation.

### Classification of evolved replicate populations

We observed different types of evolved replicates after 12 weeks of evolution (Figure 2). Closer analysis led us to believe that some populations had been invaded by asexual ‘cheaters’ that rose to a high population frequency. We believe this was driven by individuals exploiting a strategy in which they did not sporulate and/or randomly mate, while being able to survive our stringent spore isolation protocol. The observed ∼600- fold reduction in the number of mated spores by Monday evening relative to the number of diploids going into sporulation on Friday makes it clear there was a huge fitness cost to “playing by the rules”. We observed two types of cheater populations. One cheater type was characterized by very low per-site heterozygosity and appeared haploid at all but the mating type locus (where heterozygosity was maintained), with relative sequence coverage supporting haploids having either a second copy of the entire chromosome III or just the mating type locus itself. The other cheater type appears to have fixed a single, heterozygous highly recombinant diploid clone (see Note S1). In both cases, it is possible the clone that came to dominate the population was able to both exploit our scheme for enforcing outcrossing, and perhaps had a mutation of large effect that allowed it to survive the chemical challenge. The remaining populations appear to have evolved in a manner typical of sexual outbreds, with considerable genome-wide heterozygosity and raw haplotype frequencies varying in frequency across the genome. All evolved populations were thus classified into one of three classes: aneuploid haploid, clonal diploid, or outbred sexual. Per-site heterozygosity and genome-wide haplotype frequency profiles were used as metrics for classification. Specifically, populations with per-site heterozygosity profiles close to 0 with a single haplotype fixed genome-wide (except at the mating-type locus) were classified as aneuploid haploids. Populations with a bimodal per-site heterozygosity profile (with peaks centered close to 0 and 0.5) and with per-site haplotype frequencies either fixed for a single haplotype or split evenly between two haplotypes were classified as clonal diploid. The remaining populations were characterized by per-site heterozygosity profiles with a unimodal distribution and negative skew, while haplotype frequencies were highly heterogeneous throughout the genome and were classified as outbred sexual.

### Determining recombination rate in budding yeast and D. melanogaster

Two closely related metrics for estimating the recombination rate in *S. cerevisiae* vs *D. melanogaster* are used. The first estimates the average recombination rate per generation per Mb. This is calculated as 90 ∗ (1/18)/12 = 0.42, due to an estimated 90 crossovers occurring in budding yeast during meiosis with a single meiosis every 18 mitotic generations and a genome size of approximately 12Mb. For *D. melanogaster*, the sex-averaged recombination rate is calculated as 5 ∗ (1/2)/120 = 0.02, due to an estimated 5 crossovers occurring per generation in females only and a genome size of approximately 120Mb. The per-gene recombination rate is estimated similarly, with size of the genome replaced by the number of genes- approximately 14,000 in *D. melanogaster* for a recombination rate of ∼0.0002 per generation per gene as compared to approximately 6,275 genes in budding yeast for a recombination rate of ∼0.0008 per generation per gene.

### Determining sequence-level conservation at potentially functional variants

Sequence-level conservation information was taken from the UCSC genome browser based on a phylogenetic hidden Markov model (phastCons) comparing seven species of the genus *Saccharomyces*. Sequences were called as highly conserved if the level of conservation was at least 60%.

### Detecting chromosomal or segmental duplications

To scan for the presence of large-scale duplications within the 55 evolved populations that passed all filters, coverage was determined in 2kb non-overlapping intervals throughout the genome in the base and evolved populations. The relative coverage of each site was computed and the ratio of relative coverage of the evolved populations over the base population was calculated to determine the normalized fold-coverage at each site. Sites with normalized fold-coverage greater than or equal to 1.25 (which signifies that the relative coverage in the evolved population is at least 25% greater than in the base population at a particular site) were used as input for a Hidden Markov Model to predict the bounds of large, duplicated regions, which was carried out in R using a custom script.

### Detecting de novo single nucleotide variants

To scan for the presence of *de novo* mutations in all evolved populations, SNPs found in our evolved populations were considered as potential candidate *de novo* SNVs if they passed a series of filters, including: excluding SNPs present in any of the original founders or base population, including clones directly derived from the base population (Linder et al., 2020), excluding sites with less than 10x sequencing coverage, and only including SNVs at which at least 20% of the reads were called as the mutant allele. Further, to filter out spurious mutations that may have resulted as artifacts of the culturing conditions in general, mutations that showed up in more than five different chemical treatments were excluded. SNPeff (Cingolani, Platts, et al., 2012) and SNPsift (Cingolani, Patel, et al., 2012) were used to predict the impact of all *de novo* mutations detected that occurred in coding regions of annotated genes. Intergenic SNVs were then further annotated based on genomic overlap with features other than genes using the saccharomyces_cerevisiae_R64-2-1_20150113.gff file downloaded from the SGD website. Intergenic SNVs that did not overlap any known annotated genomic features were further analyzed by using them as input for the R package motifbreakR (Coetzee et al., 2015), which predicts whether or not SNPs disrupt predicted transcription factor binding sites. The list of *de novo* SNVs were finally filtered to only include the 55 evolved populations analyzed throughout this study.

### Data and software availability

Raw sequencing reads have been uploaded to the SRA under BioProject PRJNA748708. The accessions are from SRR15208200 to SRR15208448. We host some useful processed files (SNP and haplotype calls, and populations included in the study) at: http://wfitch.bio.uci.edu/~tdlong/sandvox/publications.html in the Linder et al 2021 link on the right side of the webpage. The haplotype caller is described in a previous publication (Linder et al., 2020), but the github is here: https://github.com/tdlong/yeast_resource.

## Acknowledgements

Work was funded by NIH grant FG18445 to ADL. This work was made possible, in part, through access to the Genomics High Throughput Facility Shared Resource of the Cancer Center Support Grant (P30CA-062203) at the University of California, Irvine and NIH shared instrumentation grants 1S10RR025496-01, 1S10OD010794-01, and 1S10OD021718-01.

## Supplementary Information

**Figure 1- figure supplement 1.**
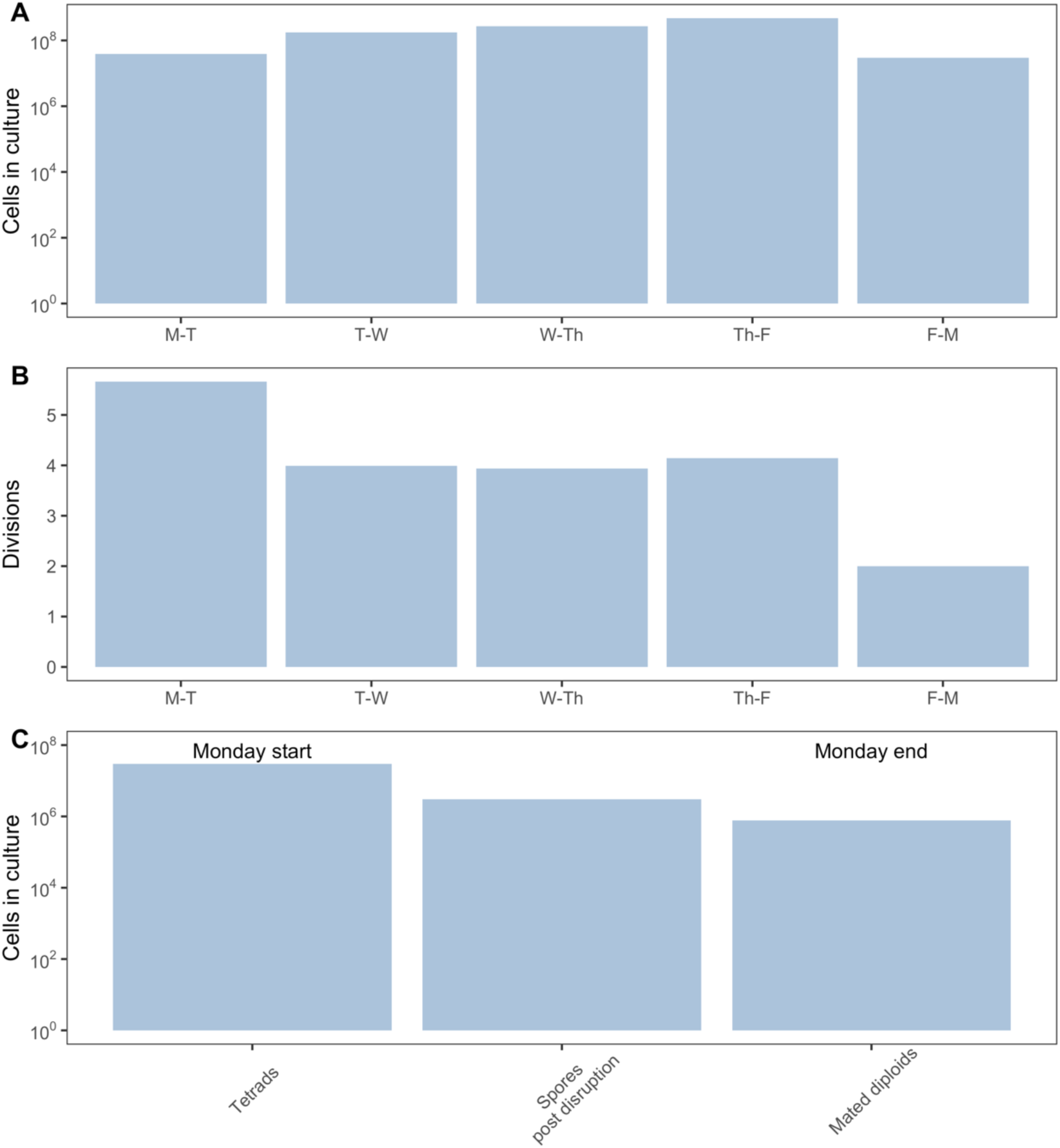
Panel (A) shows the estimated number of cells present in a YPD culture at the time of transfer going into the next day for a week of evolution. The number of mitotic and meiotic generations estimated to occur each day are shown in panel (B). One mitotic and one meiotic generation occur from Friday to Monday. Panel (C) shows the estimated total number of tetrads/cells present in a population at different stages of the Monday regimen, starting with the initial sporulated culture, then the viable spores remaining after killing unsporulated cells, followed by the viable diploids which result after random mating of isolated spores (methods). The vigorous disruption of unsporulated cells leads to a dramatic loss of viable cells (∼10-fold), with an additional ∼4-fold loss after random mating; nonetheless, over 750,000 viable individuals are left by the end of the day, which is our estimate of the population bottleneck during a typical week of evolution.

**Figure 1- figure supplement 2.**
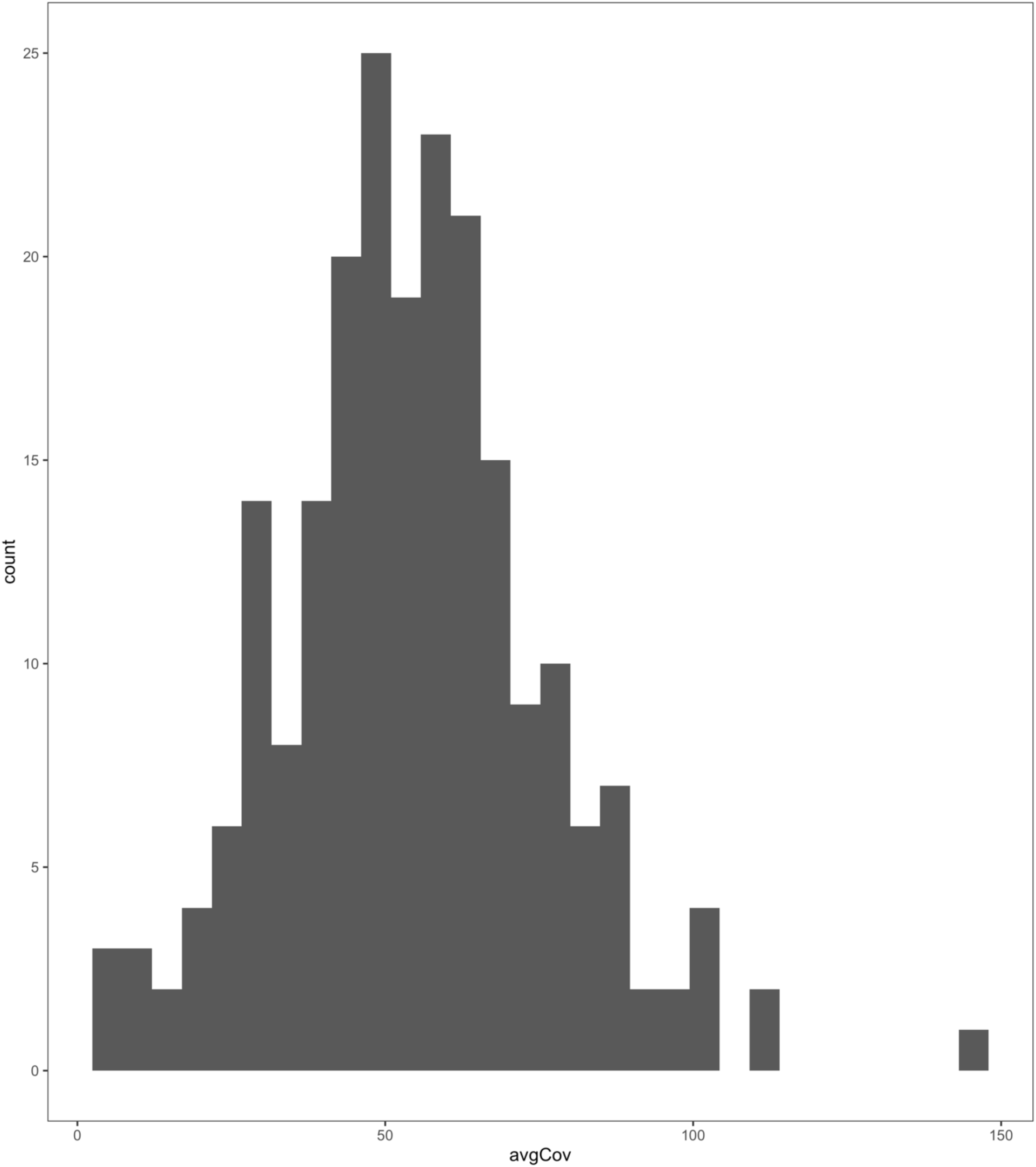
Average genome-wide short read coverage over all populations (excluding the base which was sequenced at 2226X coverage).

**Figure 2- figure supplement 1.**
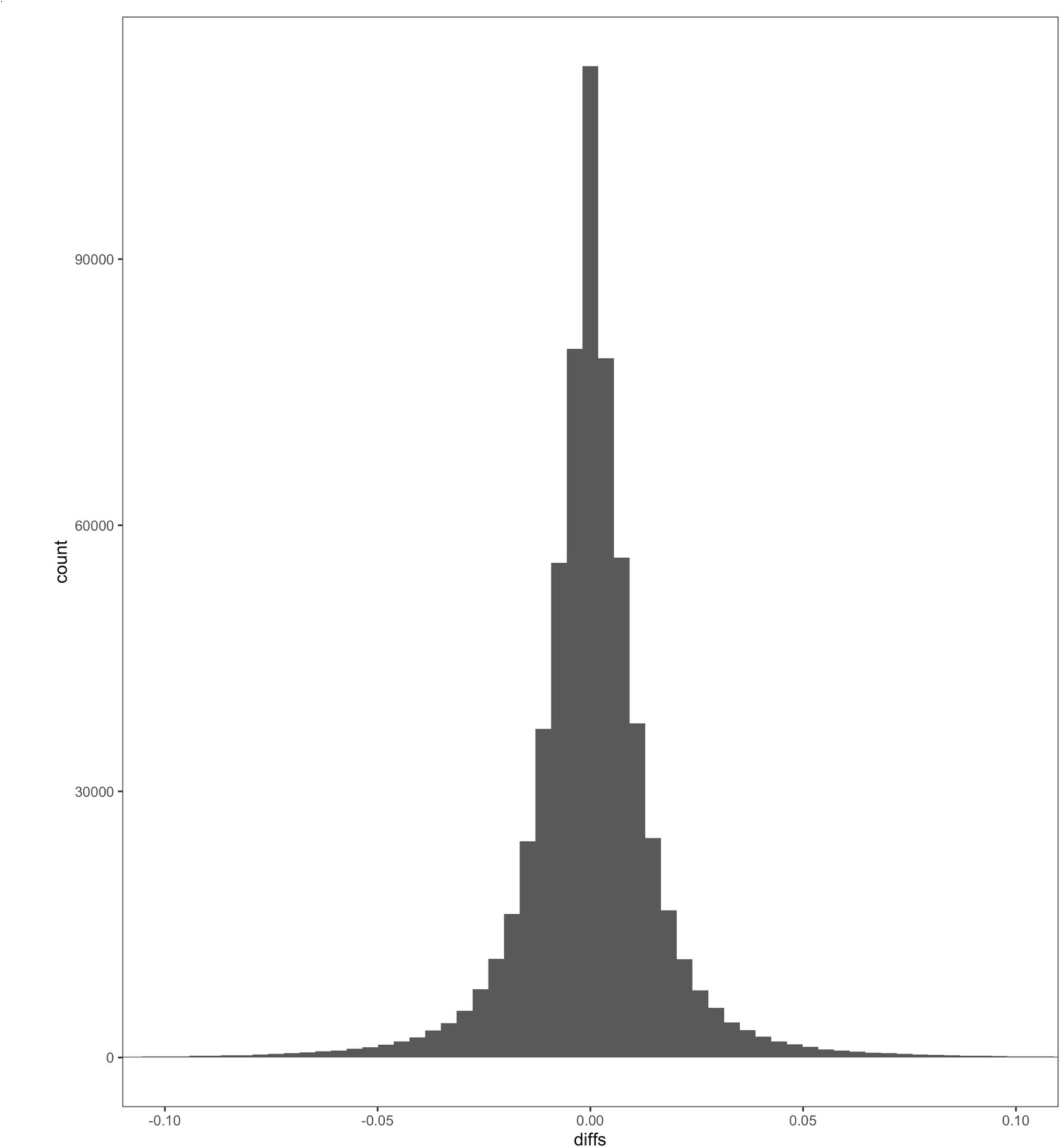
Haplotype frequency change between all adjacent intervals estimated for founder AB3. Only the 55 populations classified as outbred sexual and that clustered together from the seven chemicals used in downstream analyses were used for this analysis.

**Figure 2- figure supplement 2.**
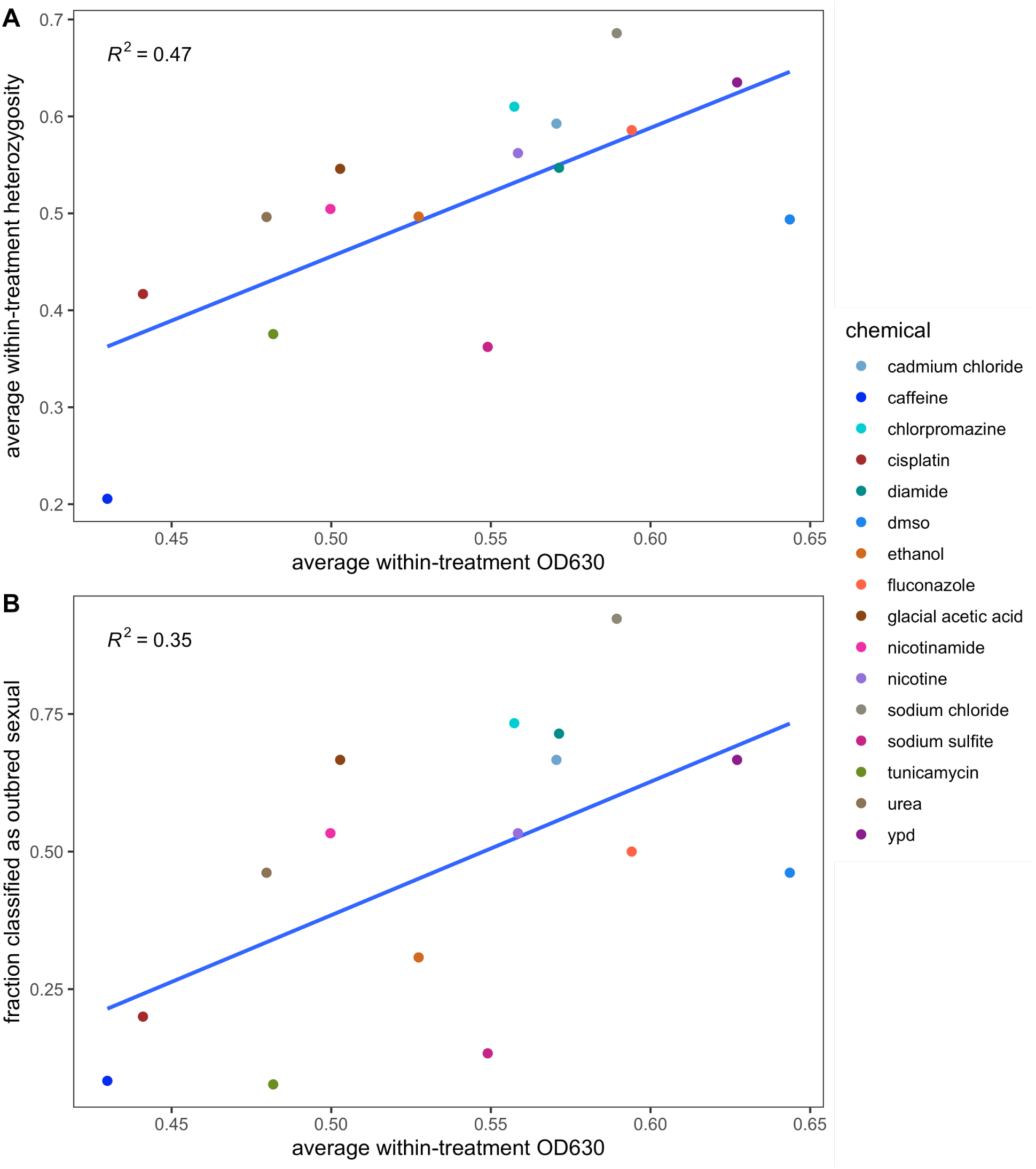
The 12-week straight average OD630 plotted against the mean per-site heterozygosity at week 12 (panel A) and the fraction of populations classified as outbred sexual (panel B) for each chemical is shown. A simple linear model regressing heterozygosity or classification on OD630 is shown. OD630 measurements were taken once a week on Thursdays for all cultures after between 23- 25h of growth, at which point cultures had not yet become fully saturated.

**Figure 2- figure supplement 3.**
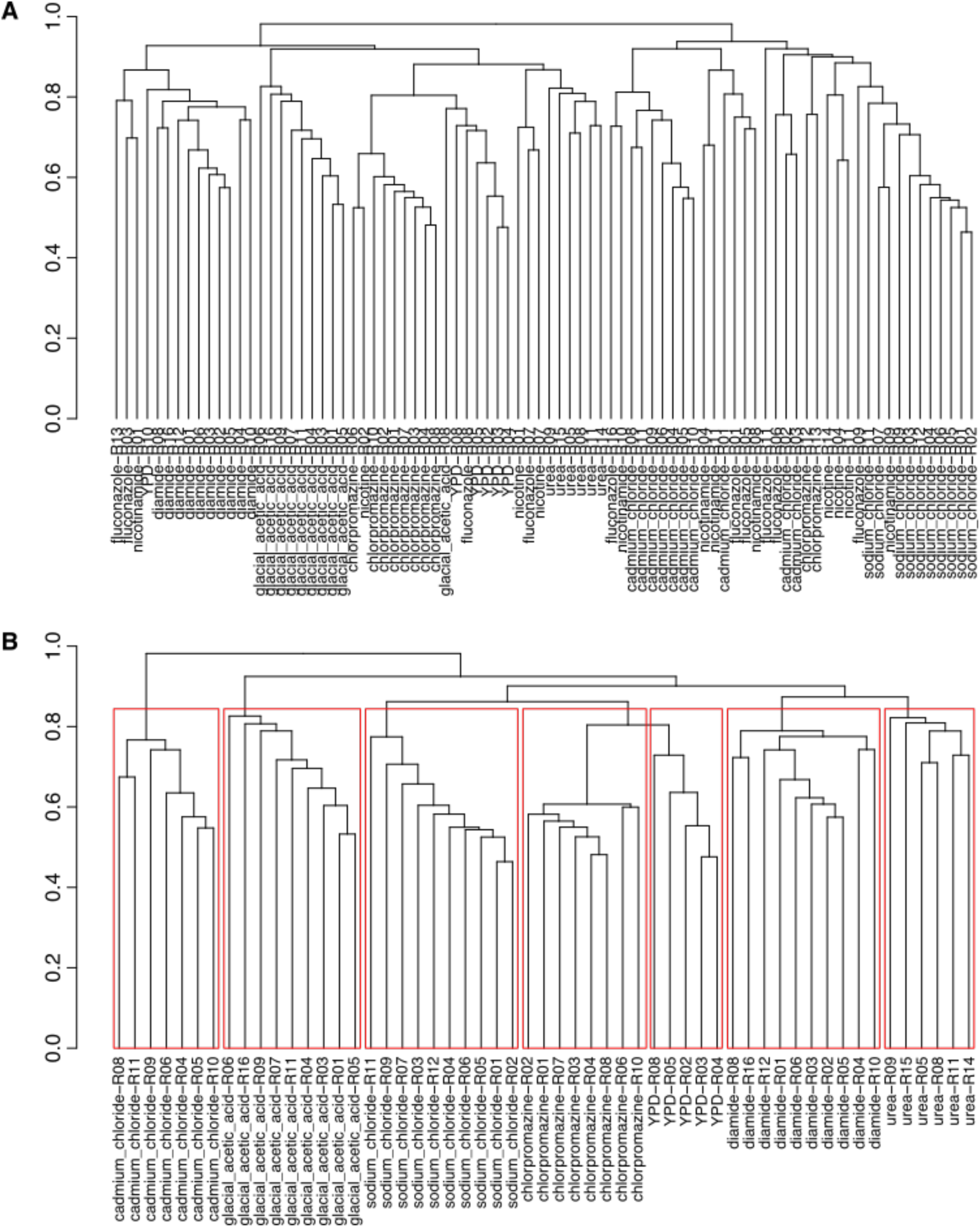
Dendrograms derived from calculating (1- Spearmans rho^2) per-site between each pair of replicates across all chemical treatments. Panel (A) shows replicates that passed the initial heterozygosity filter in chemicals with at least 6 outbred sexual replicates left, while panel (B) shows replicates that could be clustered into distinct groups by chemical stressor.

**Figure 2- figure supplement 4.**
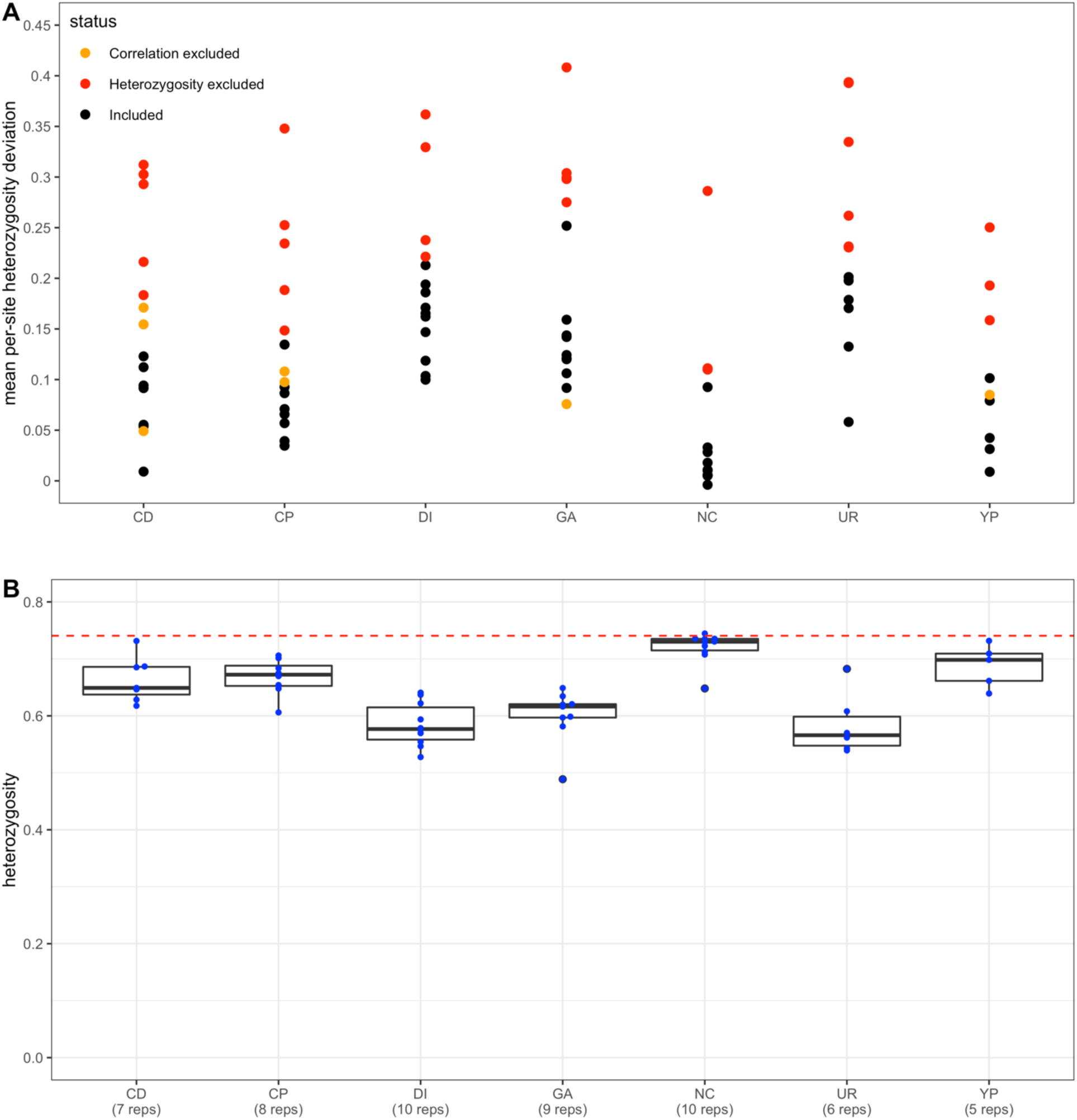
Panel (A) depicts the mean per-site heterozygosity deviation of each replicate population across the seven chemicals analyzed in this study. Replicates excluded due to either abnormal per-site heterozygosity profiles or a lack of correlation with the majority of replicates within a chemical treatment are colored red and orange, respectively. Panel (B) shows the mean per-site heterozygosity of each replicate population for each chemical treatment, with the dashed red line depicting the mean per-site heterozygosity of the base population.

**Figure 3- figure supplement 1.**
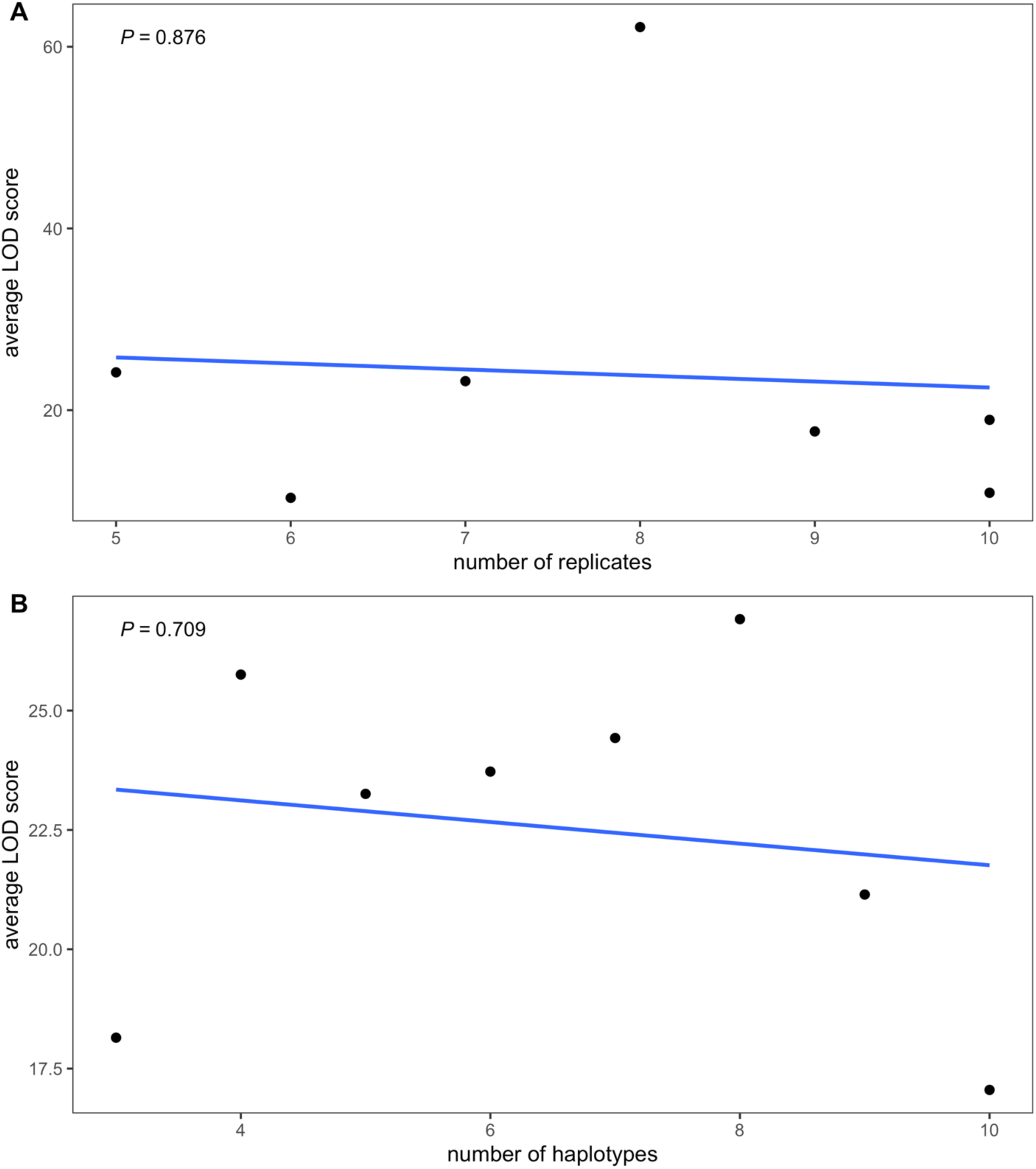
Panel (A) shows the relationship between the number of replicate populations and the average genome-wide LOD score, calculated by regressing the average LOD score on the number of replicates for each chemical. Panel (B) shows the result of a similar analysis regressing average LOD score on the number of haplotypes included for analysis at each locus over all 7 chemicals.

**Figure 4- figure supplement 1.**
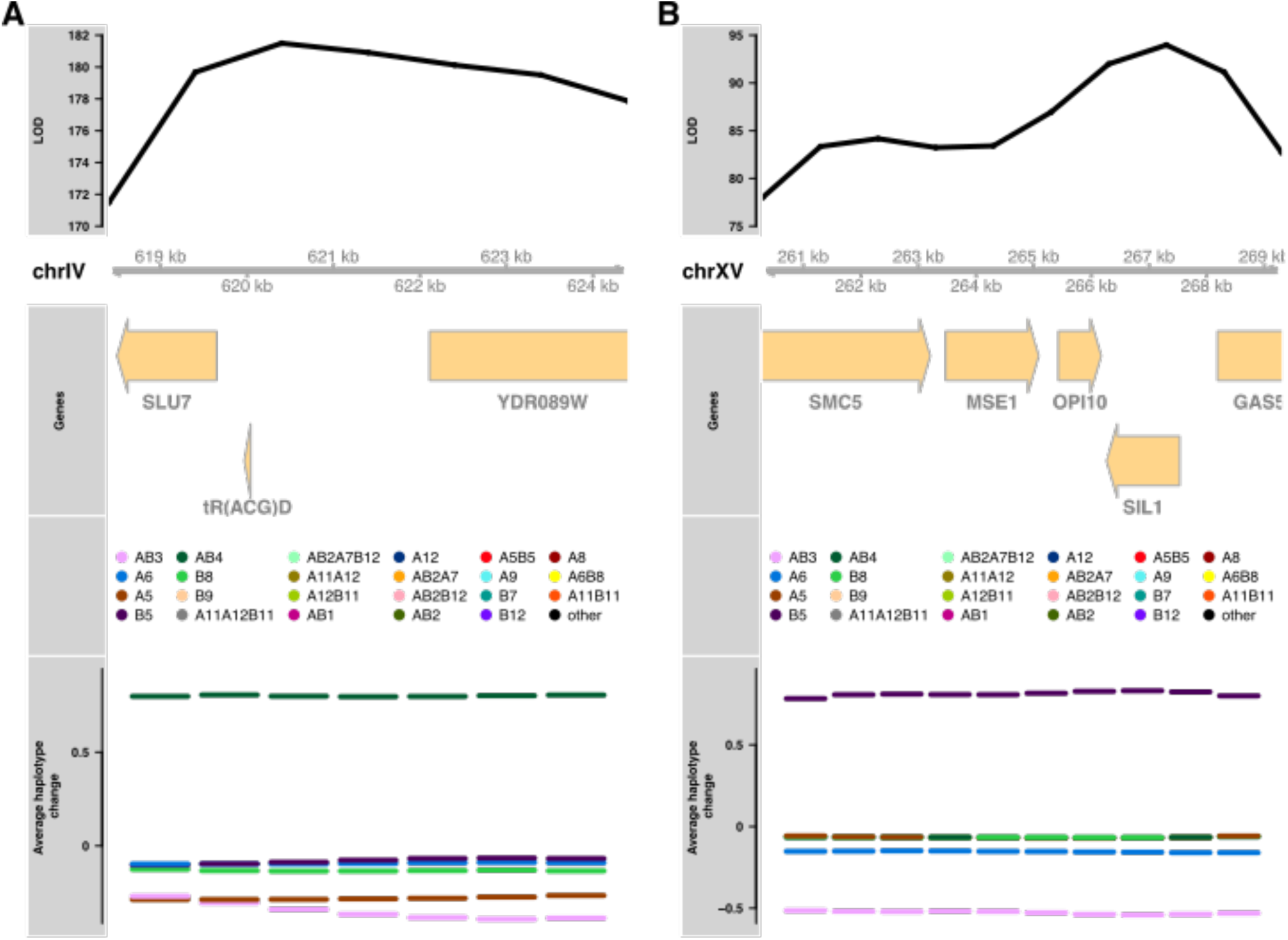
Panel (A) shows genes in a 6kb region underlying a top 3 peak in chlorpromazine and panel (B) a 9 kb region for a top 3 peak in diamide.

**Figure 4- figure supplement 2.**
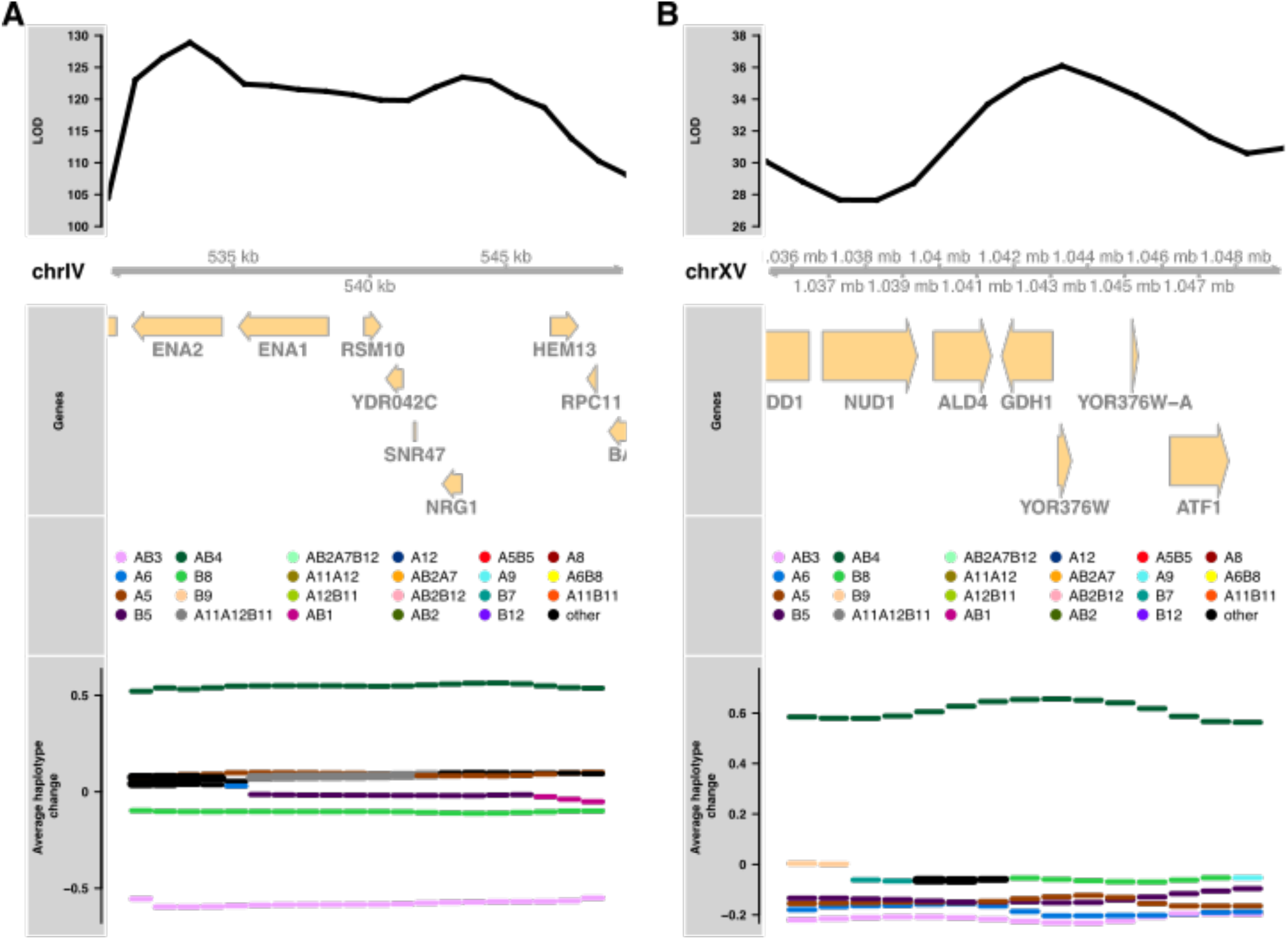
Panel (A) shows genes in a 19kb region underlying a top 3 peak in sodium chloride and panel (B) a 14kb region for a top 3 peak in urea.

**Figure 5- figure supplement 1.**
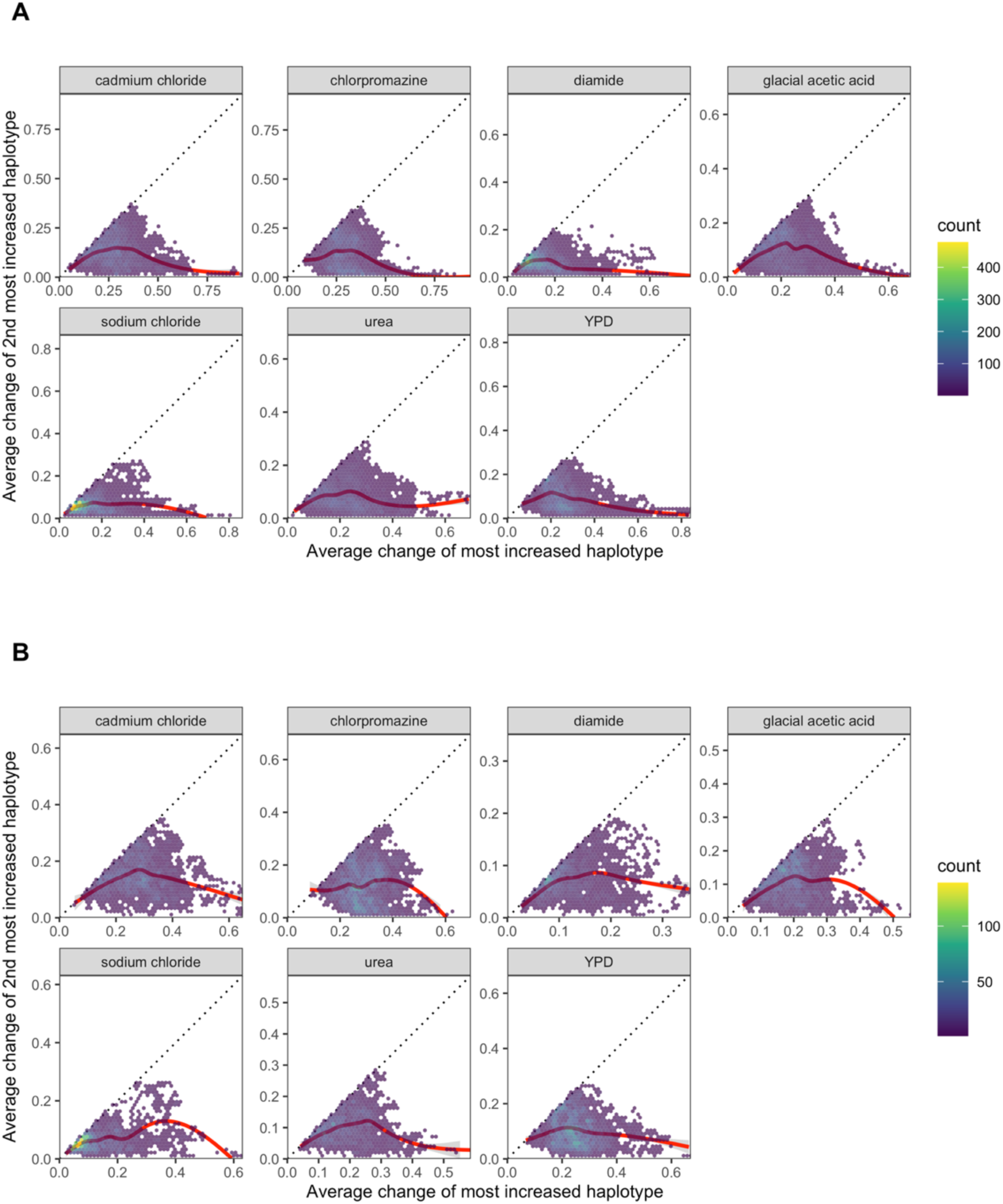
Panel (A) shows the average change of the Most Increased Haplotype (MIH) vs that of the second most increased haplotype within each chemical treatment across all loci. Panel (B) is similar except that only loci at which the second most increased haplotype was at a higher starting frequency in the base population than the MIH are plotted. Loess smoothing is represented as red lines.

**Figure 6- figure supplement 1.**
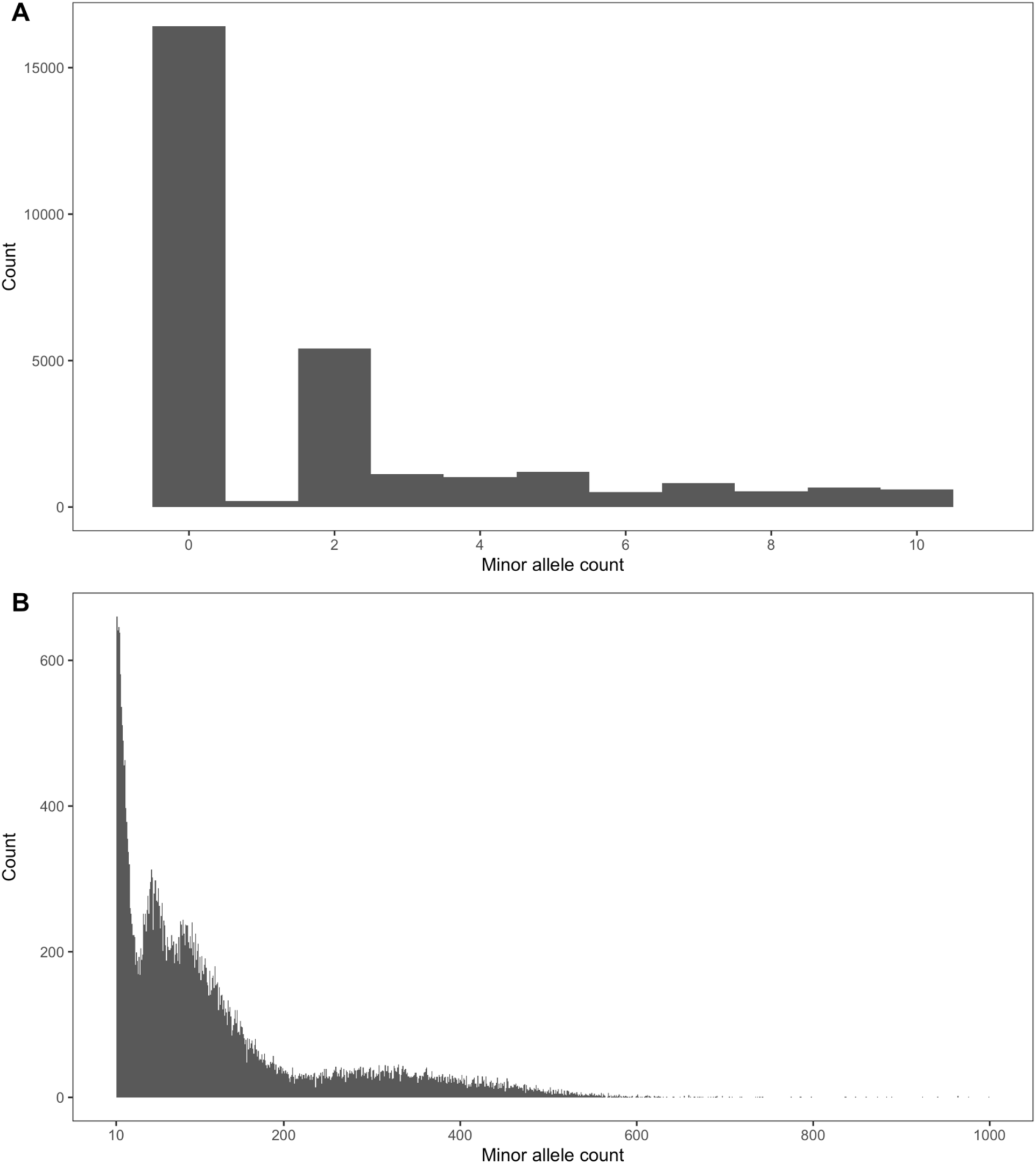
Panel (A) shows the number of times the minor allele of all SNPs private to a single founder received between 0-10 sequencing reads in the base population (the Minor allele count), which was sequenced to a depth of 2226X. Panel (B) is similar but the range is from 10-1000 sequencing reads in the base population.

**Figure 6- figure supplement 2.**
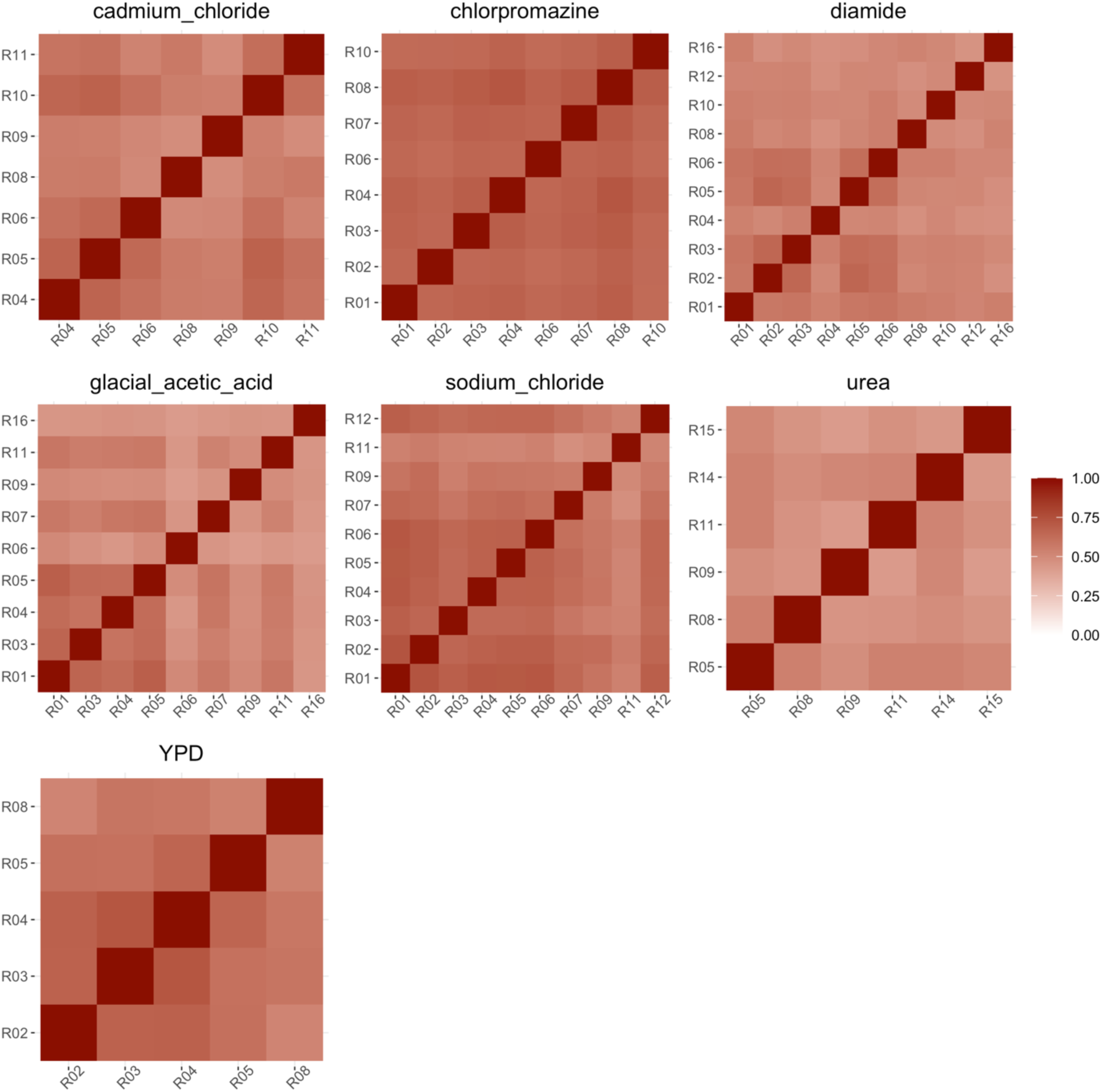
Heat maps of the Spearman correlations within chemical treatments.

**Figure 6- figure supplement 3.**
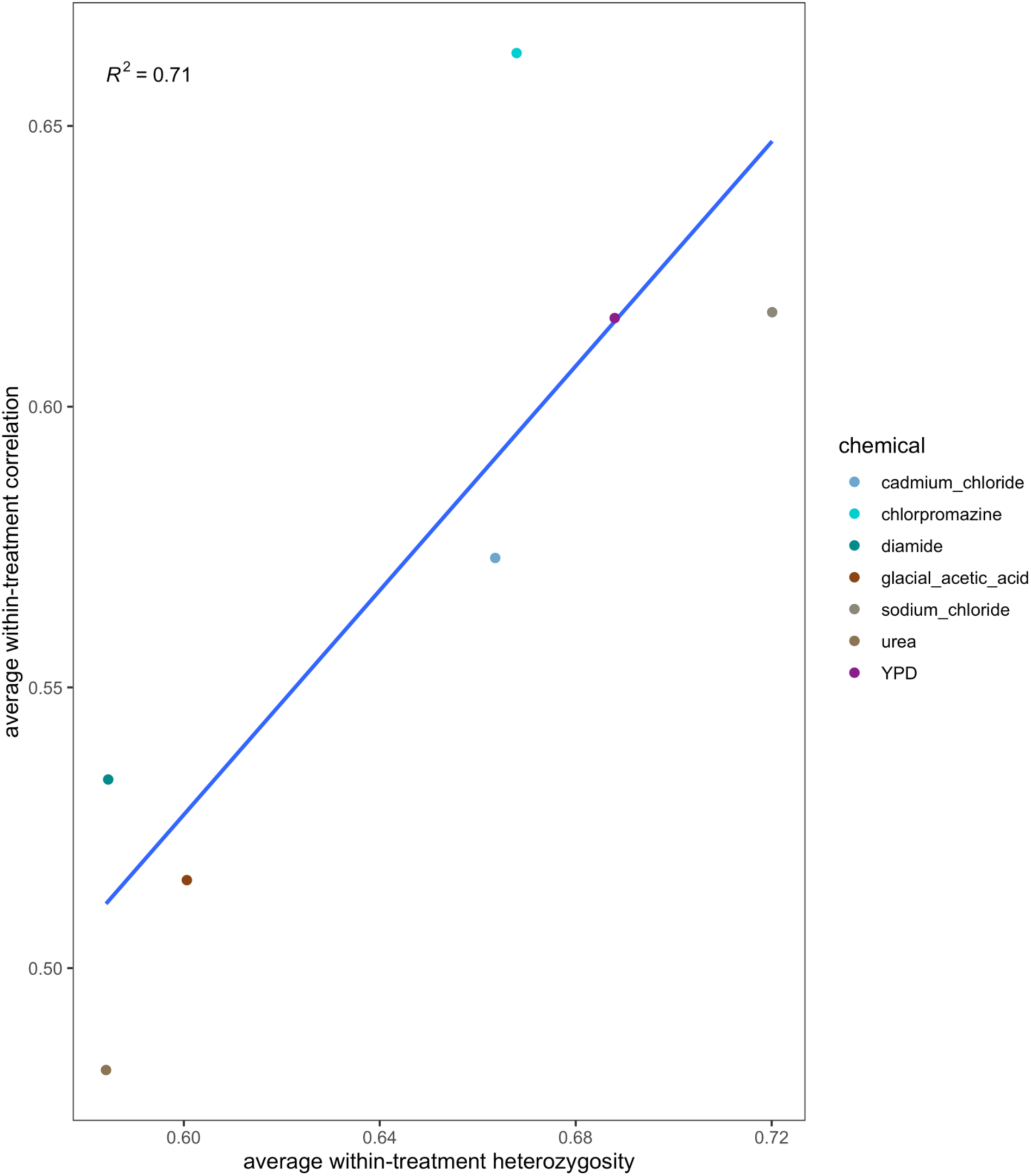
The correlation between the average heterozygosity and the mean within treatment Spearman correlation.

**Figure 6- figure supplement 4.**
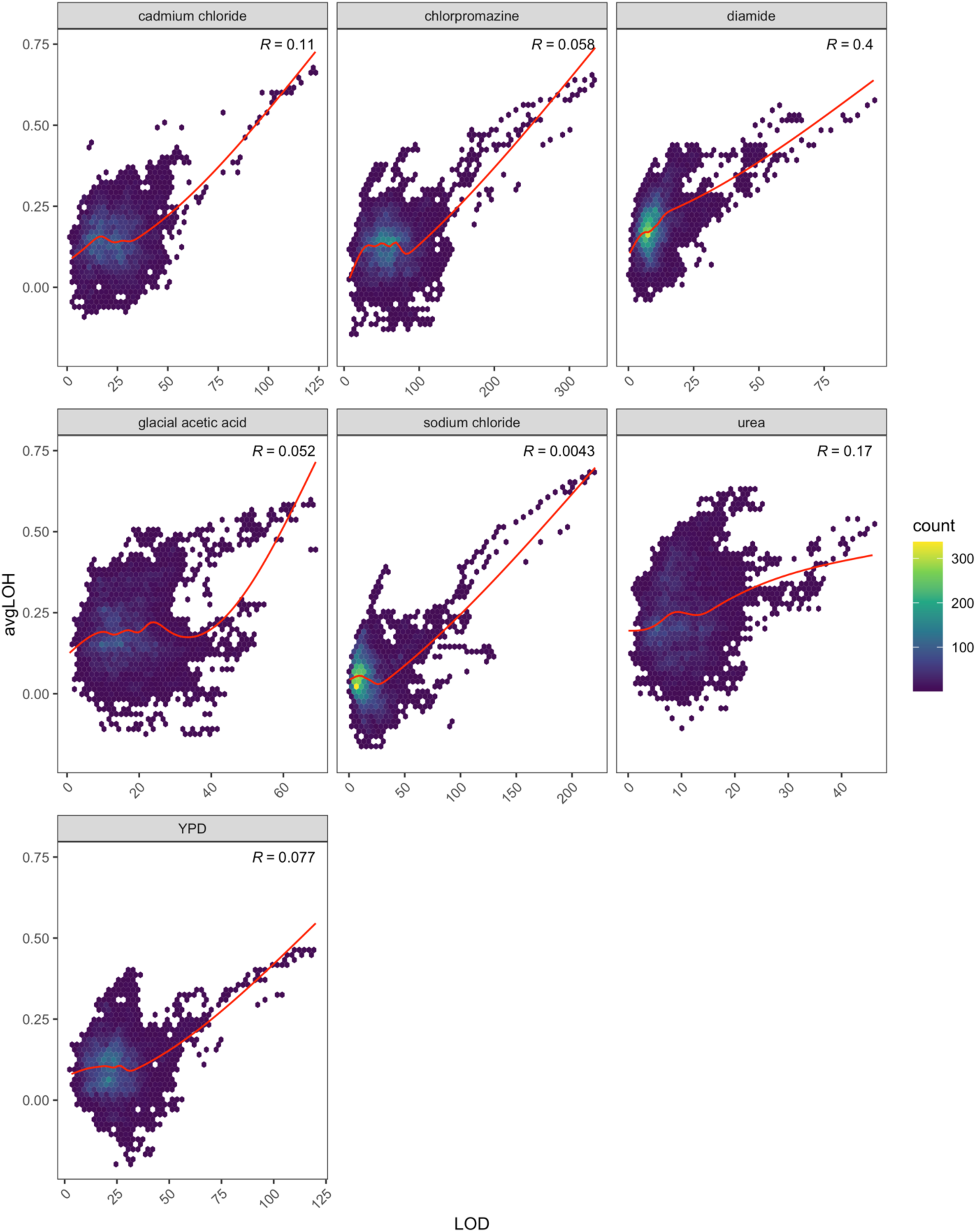
The correlation between LOD score and the average loss-of-heterozygosity per site genome-wide.

**Figure 6- figure supplement 5.**
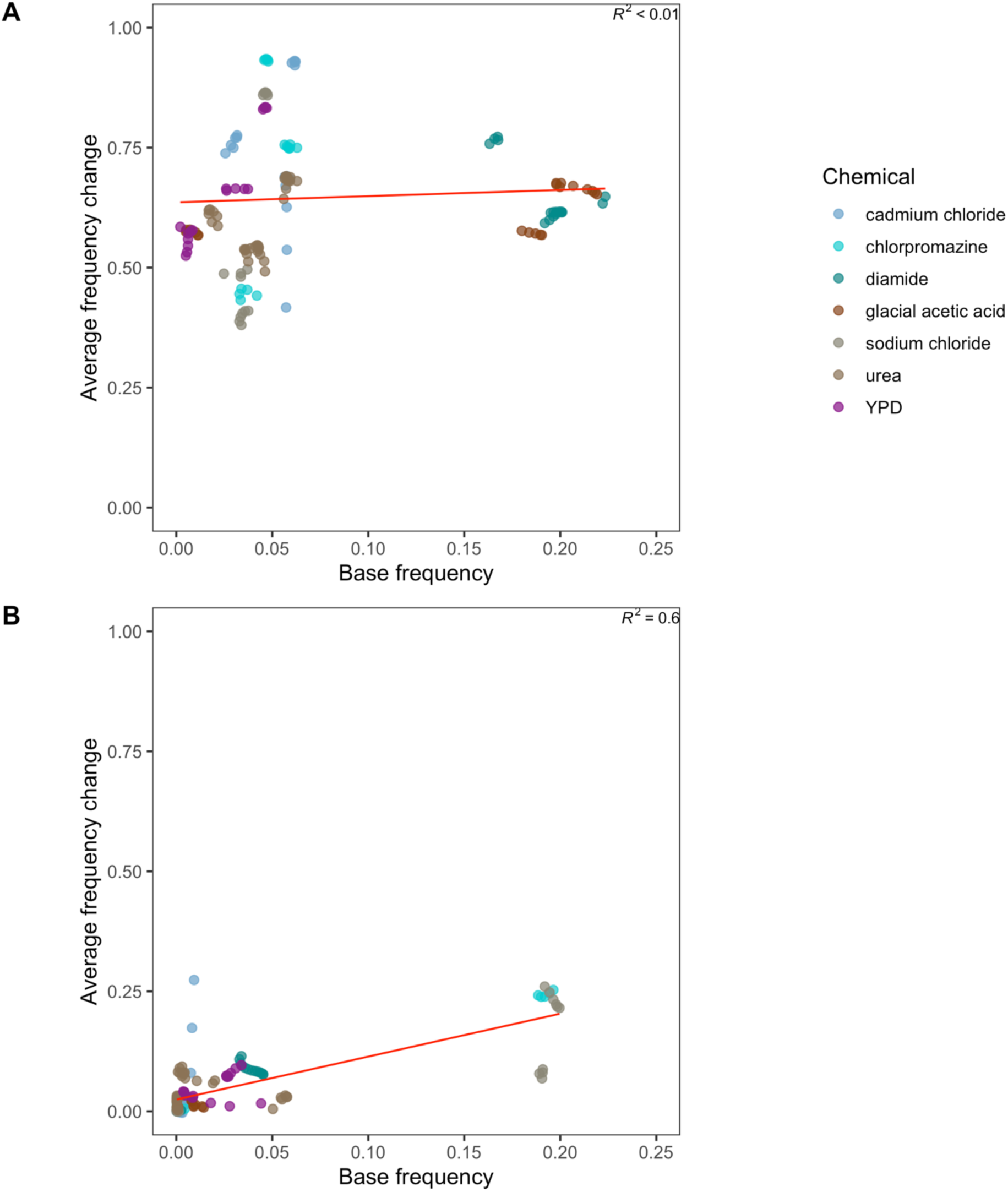
Panel (A) shows the average frequency change of the most increased haplotype while panel (B) shows the next most increased haplotype at each of the 21 major peaks detected regressed onto the initial frequency of the respective haplotypes in the base population.

**Figure 6- figure supplement 6.**
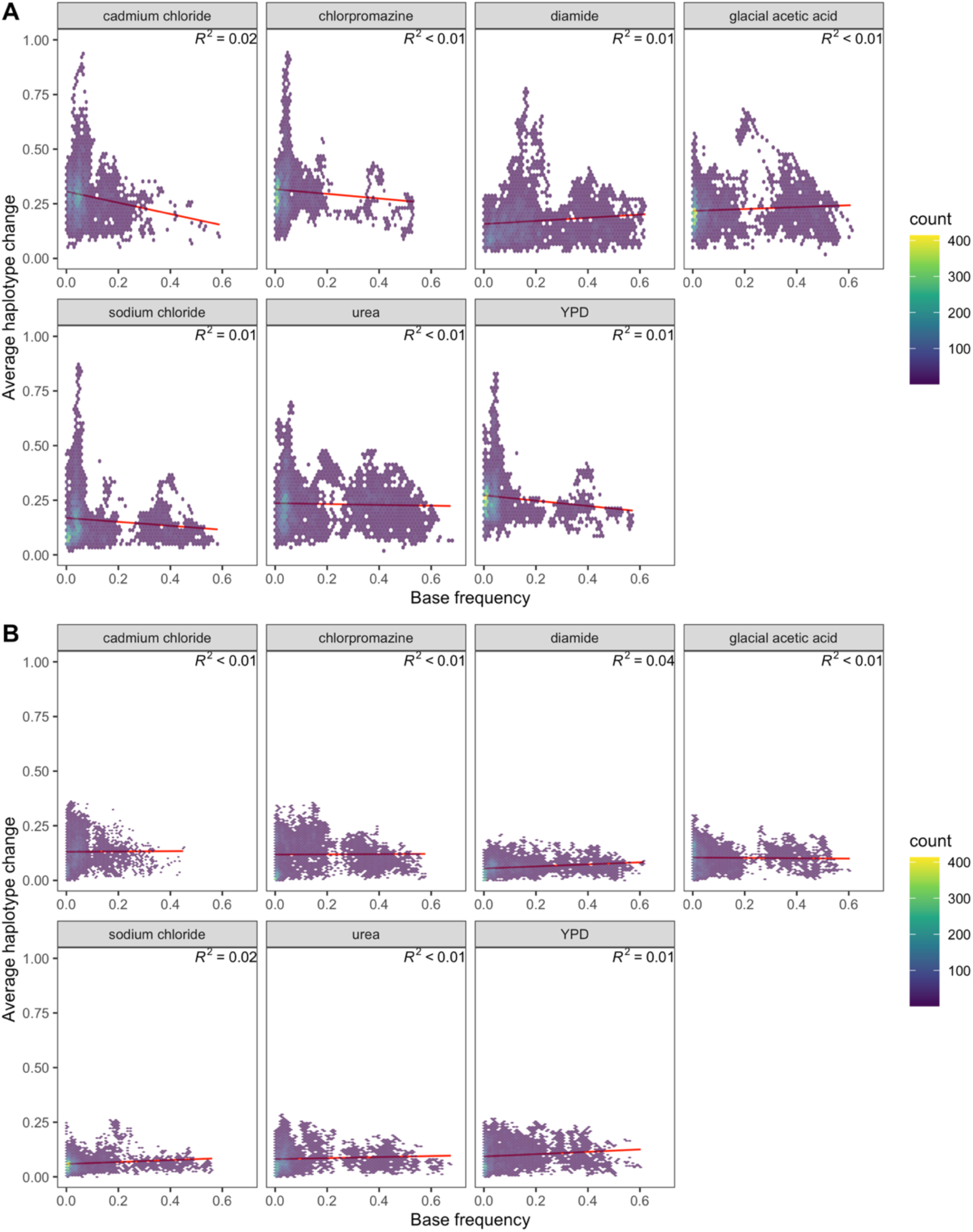
Panel (A) shows the average frequency change of the most increased haplotype while panel (B) shows the next most increased haplotype per-site genome-wide regressed onto the initial frequency of the respective haplotypes in the base population.

**Figure 9- figure supplement 1.**
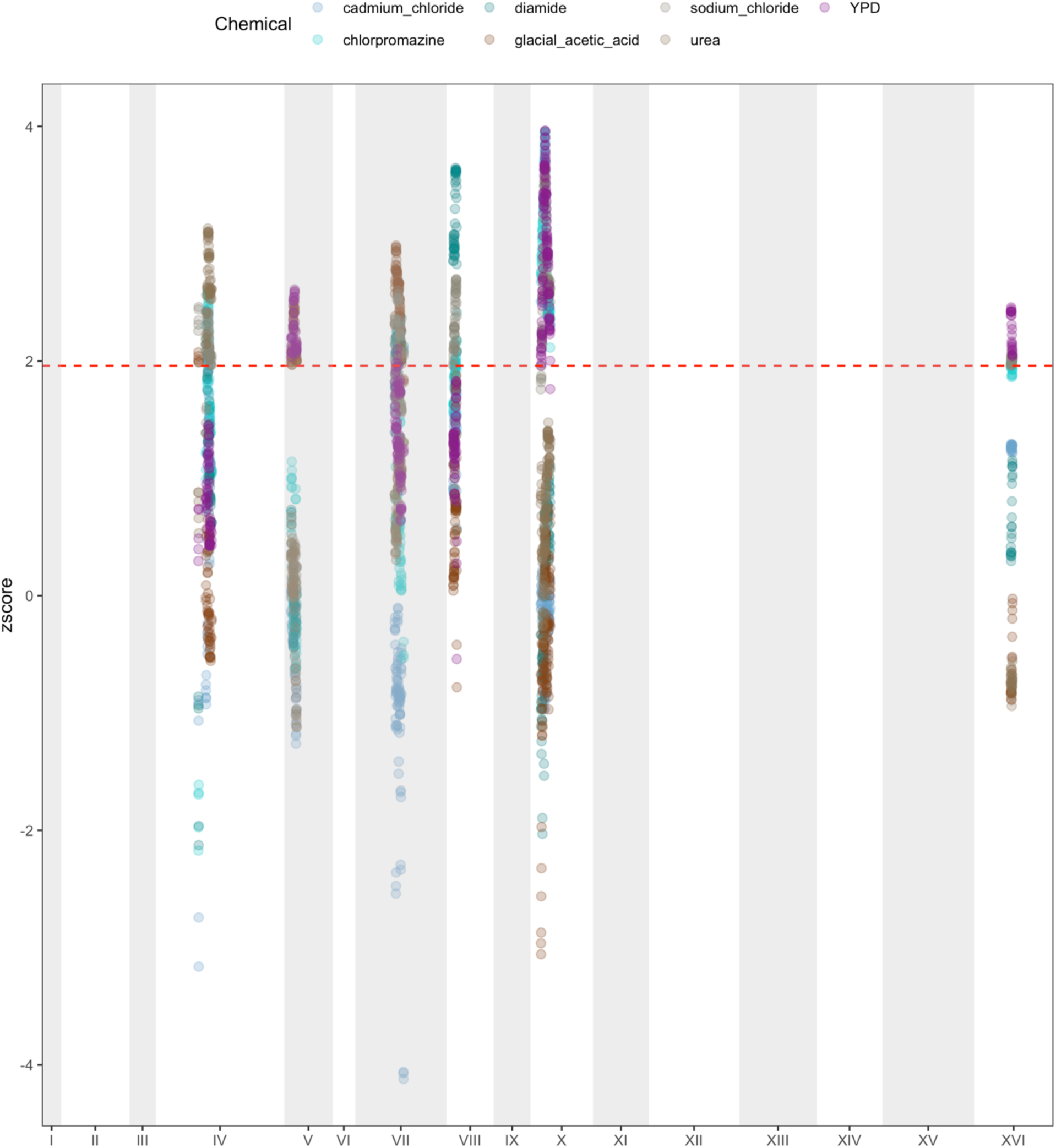
Only regions in which two or more chemicals had a z- score >= 1.96 were plotted to find regions that may represent instances of pleiotropy.

**Figure 9- figure supplement 2.**
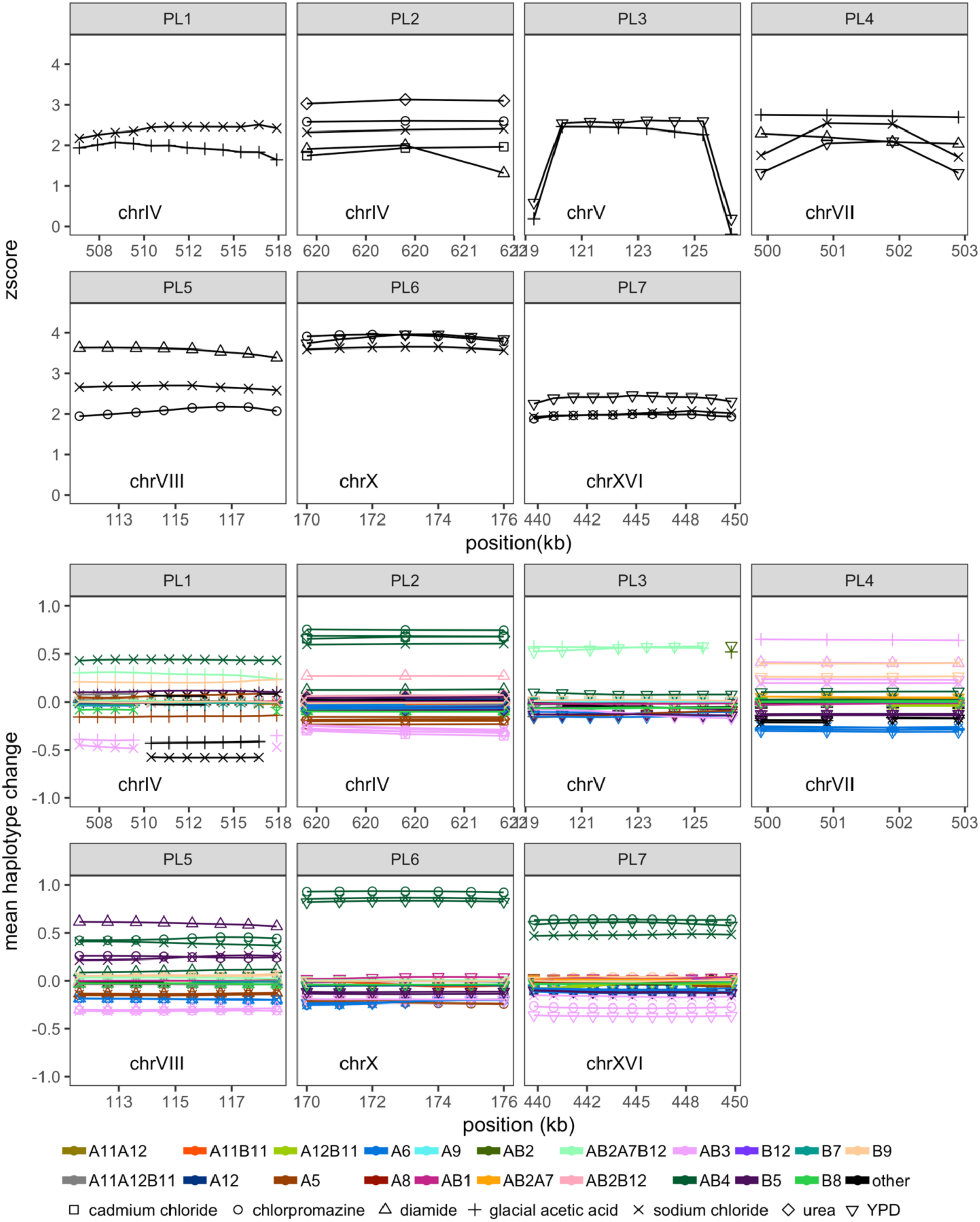
Potentially pleiotropic regions (labelled by peak ID) are shown. The z-score of transformed LOD scores in shown in panel (A), while panel (B) shows the mean haplotype frequency change across each region.

**Table S1.** Chemicals and doses used in this study

| Chemical | Day 1 dose | Days 2+ dose | General Impact | Effects in budding yeast |
| --- | --- | --- | --- | --- |
| Cadmium chloride | 400uM | 600uM | Widespread environmental contaminant and type I carcinogen | Induces oxidative stress and heavy metal toxicity |
| Caffeine | 16mM | 18mM | Widely used legal stimulant that is highly addictive | Inhibits homologous recombination and extends lifespan in yeast by inhibiting RAD51 and mTORC1, respectively |
| Chlorpromazine | 8uM | 10uM | Commonly prescribed anti-psychotic with idiosyncratic toxicity | Induces oxidative stress and the unfolded protein response, inhibits protein synthesis, alters membrane integrity, and inhibits intracellular protein trafficking |
| Cisplatin | 400uM | 600uM | Frequently used chemotherapeutic agent with high rates of intrinsic and acquired resistance | Induces DNA damage by crosslinking with purine bases |
| Diamide | 2.5mM | 2.7mM | Widely used insecticide | Induces oxidative stress |
| DMSO | 8.50% | 7% | Widely used organic solvent | Induces oxidative stress |
| Ethanol | 12.50% | 12.50% | Alcohol commonly encountered in nature | Induces oxidative stress |
| Fluconazole | 100uM | 5uM | Anti-fungal | Interferes with formation of the cell membrane |
| Glacial acetic acid | 100mM | 120mM | Accumulates in yeast culture during stationary phase | Induces oxidative stress and causes intracellular acidification |
| Sodium chloride | 900mM | 950mM | Commonly used preservative and food additive | Induces osmotic stress |
| Nicotinamide | 220mM | 225mM | NAD<sup>+</sup> precursor shown to protect against obesity and extend lifespan in mice | Protects against DNA damage |
| Nicotine | 7mM | 9mM | Widely used legal stimulant that is highly addictive | Induces oxidative stress |
| Sodium sulphite | 85mM | 90mM | Used as a preservative in winemaking | Causes ATP depletion |
| Tunicamycin | 0.8uM | 1uM | Inhibits tumor cell growth | Inhibits protein glycosylation and formation of N-acetylglucosamine lipid intermediates |
| Urea | 900mM | 650mM | Inovolved in nitrogen excretion and used in fertilizers | Denatures proteins |
| YPD | NA | NA | Rich media used for yeast cell culturing | Supports rapid growth of budding yeast |

**Table S2.**
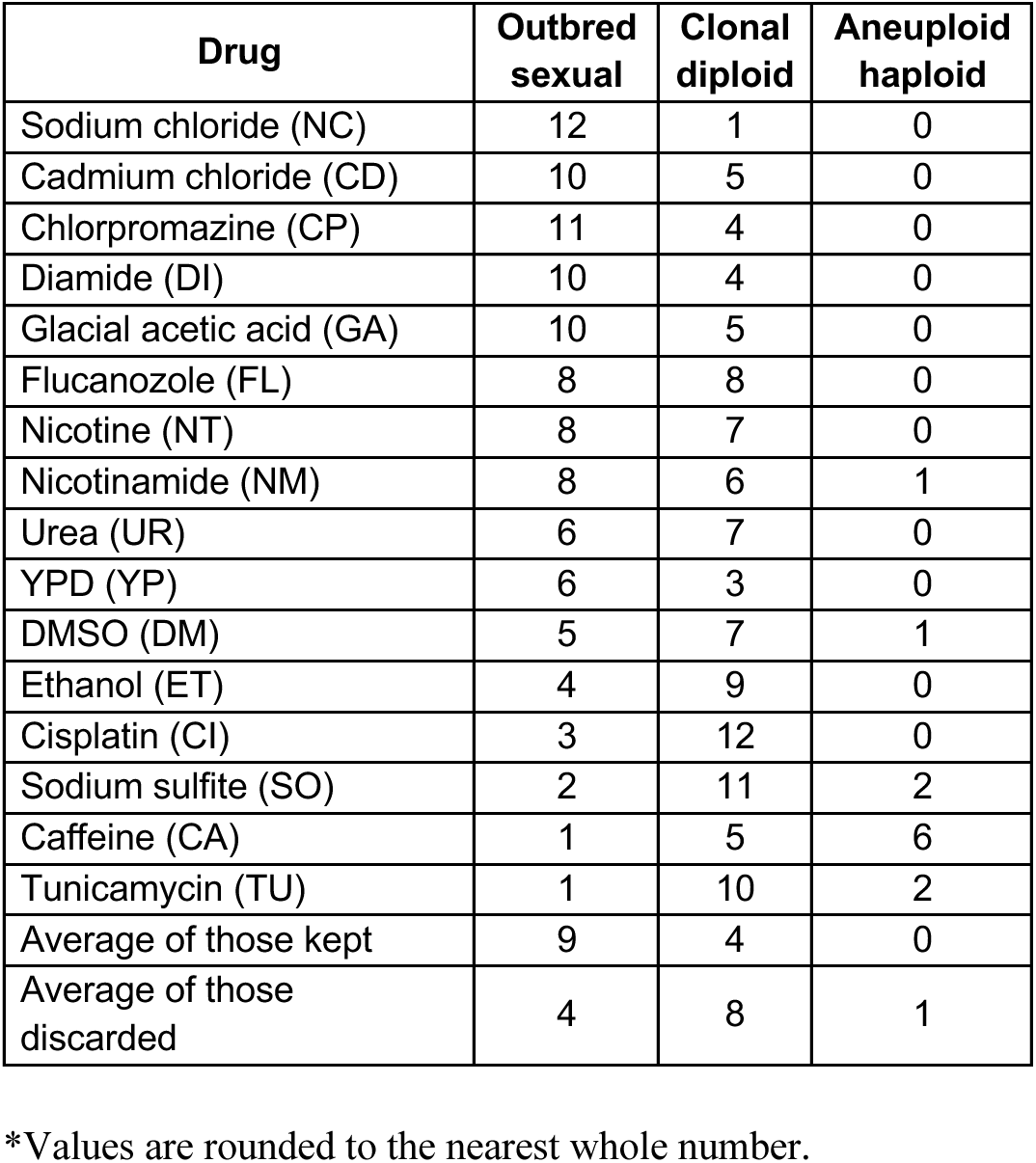
Count of populations per chemical detected in each class

**Table S3.** Large-scale duplications detected in this study.

| Chemical | chr | Interval | Total<br>reps | Replicate(fold<br>change) | Duplication<br>type |
| --- | --- | --- | --- | --- | --- |
| CD | II | NA | 7 | R04(1.4),<br>R05(1.4),<br>R06(1.4),<br>R08(1.4),<br>R09(1.4),<br>R10(1.4),<br>R11(1.2) | whole |
| CP | III | 292,285-310,285 | 8 | R02(1) | partial |
| CP | XVI | 912,842-940,842 | 8 | R02(1.3) | partial |
| CP | XVI | 920,842-940,842 | 8 | R03(1.3),<br>R06(1.3),<br>R08(1.2) | partial |
| DI | III | 158,285-226,285 | 10 | R10(1.4) | partial |
| DI | III | 160,285-226,285 | 10 | R16(1.3) | partial |
| DI | VI | 206,139-264,139 | 10 | R01(2),<br>R02(1.9),<br>R03(1.8),<br>R04(2), R05(2),<br>R06(2),<br>R08(1.9),<br>R10(2), R12(2),<br>R16(2) | partial |
| DI | XVI | 920,842-940,842 | 10 | R01(1.3),<br>R04(1.2) | partial |
| GA | II | 801,444-805,444 | 9 | R09(1.2) | partial |
| GA | VII | 1,049,893-1,081,893 | 9 | R03(1) | partial |
| GA | XVI | 12,842-202,842 | 9 | R06(1.3) | partial |
| GA | XVI | 12,842-228,842 | 9 | R04(1.2) | partial |
| GA | XVI | 12,842-368,842 | 9 | R11(1.3),<br>R16(1.4) | partial |
| GA | XVI | 12,842-370,842 | 9 | R07(1.4) | partial |
| GA | XVI | 22,842-368,842 | 9 | R09(1.4) | partial |
| GA | XVI | 920,842-940,842 | 9 | R09(1.3) | partial |
| GA | XVI | 926,842-940,842 | 9 | R05(1.3) | partial |
| NC | III | 200,285-310,285 | 10 | R03(1.3),<br>R07(1.4) | partial |
| NC | III | NA | 10 | R11(1.3) | whole |
| NC | V | NA | 10 | R11(1.2) | whole |
| NC | VI | 256,139-264,139 | 10 | R07(1.1) | partial |
| NC | XVI | 920,842-940,842 | 10 | R02(1.2) | partial |
| UR | V | 325,329-565,329 | 6 | R15(1.2) | partial |
| UR | V | NA | 6 | R08(1.4),<br>R09(1.4),<br>R14(1.3) | whole |
| UR | VI | 204,139-264,139 | 6 | R09(1.1) | partial |
| UR | XIII | 916,402-922,402 | 6 | R09(1.1) | partial |
| UR | XVI | 936,842-940,842 | 6 | R15(1.4) | partial |
| YP | XVI | 920,842-940,842 | 5 | R08(1.2) | partial |

**Table S4.** Single nucleotide variants detected in this study.

| Chemical | chr | Pos | Total<br>reps | Replicate<br>(frequency) | mutation | Mutation type | Gene hit |
| --- | --- | --- | --- | --- | --- | --- | --- |
| CD | I | 106,071 | 7 | R04(0.2) | C>A | TFBS_Disruption | LTE1 |
| CD | IV | 1,924 | 7 | R06(0.2) | T>C | Synonymous | COS7 |
| CD | IV | 169,930 | 7 | R10(0.2) | T>A | TFBS_Disruption | MHF2 |
| CD | IV | 865,025 | 7 | R04(0.3) | A>T | Missense | UME6 |
| CD | V | 520,777 | 7 | R06(0.3) | G>A | Intergenic | ARS521 |
| CD | VI | 6,810 | 7 | R05(0.2) | C>T | Synonymous | COS4 |
| CD | VII | 611,564 | 7 | R06(0.2) | G>T | Intergenic | ERG25 |
| CD | XI | 61,163 | 7 | R09(0.2) | G>T | Missense | TOR2 |
| CD | XII | 873,696 | 7 | R11(0.2) | C>T | TFBS_Disruption | PSY3 |
| CD | XIII | 472,636 | 7 | R06(0.3) | T>A | TFBS_Disruption | YMR102C |
| CD | XIII | 559,094 | 7 | R06(0.2) | G>A | Intergenic | TIF34 |
| CD | XIII | 865,417 | 7 | R10(0.2) | C>T | TFBS_Disruption | DYN3 |
| CD | XIV | 196,170 | 7 | R05(0.6) | A>C | Intergenic | ARS1412 |
| CD | XVI | 287,581 | 7 | R05(0.2) | A>T | Missense | MKK2 |
| CP | II | 166,036 | 8 | R10(0.2) | A>T | TFBS_Disruption | YBL029W |
| CP | II | 251,994 | 8 | R02(0.2) | G>T | TFBS_Disruption | DSF2 |
| CP | II | 251,995 | 8 | R02(0.2) | A>T | TFBS_Disruption | DSF2 |
| CP | II | 251,999 | 8 | R02(0.2) | G>T | Intergenic | DSF2 |
| CP | II | 505,129 | 8 | R02(0.7) | G>A | Synonymous | CKS1 |
| CP | III | 1,902 | 8 | R06(0.2) | G>A | Missense | YCL076W |
| CP | IV | 693,336 | 8 | R02(0.3) | G>A | TFBS_Disruption | TRM1 |
| CP | V | 492,019 | 8 | R07(0.2) | G>T | TFBS_Disruption | BUR6 |
| CP | IX | 415,829 | 8 | R02(0.4) | A>T | TFBS_Disruption | DAL3 |
| CP | X | 173,267 | 8 | R06(0.3) | C>A | Intergenic | URA2 |
| CP | X | 176,417 | 8 | R02(0.2) | A>T | Missense | TRK1 |
| CP | X | 176,418 | 8 | R02(0.2) | A>T | Missense | TRK1 |
| CP | X | 193,334 | 8 | R02(0.3) | T>A | Synonymous | PHO86 |
| CP | X | 193,335 | 8 | R02(0.2) | T>A | Missense | PHO86 |
| CP | X | 193,336 | 8 | R02(0.3) | T>A | Premature_stop | PHO86 |
| CP | X | 675,076 | 8 | R10(0.9) | T>C | Synonymous | SGM1 |
| CP | XI | 286,208 | 8 | R04(0.3) | A>T | Stop_lost | VMA5 |
| CP | XI | 398,709 | 8 | R06(0.3),<br>R08(0.2) | C>A | Missense | MAK11 |
| CP | XI | 450,307 | 8 | R01(0.3) | G>A | TFBS_Disruption | YKR005C |
| CP | XI | 489,811 | 8 | R03(0.2) | A>T | TFBS_Disruption | GCN3 |
| CP | XII | 122,034 | 8 | R10(0.6) | A>C | Intergenic | PUF3 |
| CP | XII | 533,006 | 8 | R02(0.2) | C>T | Missense | ATG26 |
| CP | XII | 547,549 | 8 | R10(0.2) | G>A | Missense | NOP56 |
| CP | XII | 587,566 | 8 | R02(0.2) | G>A | Missense | YLR224W |
| CP | XII | 1,042,117 | 8 | R10(0.3) | A>T | Premature_stop | RIF2 |
| CP | XIII | 225,438 | 8 | R02(0.5) | T>C | TFBS_Disruption | YML6 |
| CP | XIII | 677,460 | 8 | R02(0.2) | C>T | Synonymous | HFA1 |
| CP | XIII | 677,461 | 8 | R02(0.3) | A>T | Missense | HFA1 |
| CP | XV | 37,749 | 8 | R02(0.3) | A>G | Intergenic | YOL153C |
| CP | XV | 227,560 | 8 | R06(0.3) | T>A | TFBS_Disruption | tG(GCC)O1 |
| CP | XV | 1,079,939 | 8 | R08(0.3) | T>A | TFBS_Disruption | HSP33 |
| CP | XV | 1,079,940 | 8 | R08(0.3) | T>A | TFBS_Disruption | HSP33 |
| CP | XVI | 814,209 | 8 | R10(0.2) | G>T | Intergenic | LOA1 |
| DI | I | 662 | 10 | R02(0.2) | C>T | Missense | YAL068W-A |
| DI | I | 45,628 | 10 | R01(0.2) | C>T | TFBS_Disruption | ACS1 |
| DI | I | 220,215 | 10 | R08(0.4) | C>T | Synonymous | YAR064W |
| DI | I | 226,241 | 10 | R06(0.2) | C>T | Missense | PHO11 |
| DI | II | 45,232 | 10 | R05(0.2) | A>T | TFBS_Disruption | ROX3 |
| DI | II | 69,628 | 10 | R02(0.3) | C>A | Intergenic | CDC27 |
| DI | II | 645,983 | 10 | R01(0.2) | G>A | Intergenic | ERV15 |
| DI | IV | 60,293 | 10 | R12(0.3) | C>T | Missense | HBT1 |
| DI | IV | 60,294 | 10 | R12(0.3) | G>T | Missense | HBT1 |
| DI | IV | 383,478 | 10 | R06(0.3) | T>C | Missense | PRM7 |
| DI | IV | 1,442,247 | 10 | R12(0.2) | A>T | Missense | PUF6 |
| DI | IV | 1,517,694 | 10 | R02(0.2) | A>T | TFBS_Disruption | IRC4 |
| DI | V | 148,091 | 10 | R04(0.2) | G>A | TFBS_Disruption | GIM4 |
| DI | V | 386,140 | 10 | R01(0.3) | C>A | TFBS_Disruption | SWI4 |
| DI | V | 492,007 | 10 | R01(0.2) | A>T | Intergenic | BUR6 |
| DI | V | 516,605 | 10 | R03(0.2) | C>T | Missense | DNF1 |
| DI | VII | 687,607 | 10 | R12(0.3) | T>C | TFBS_Disruption | ESP1 |
| DI | VII | 1,011,014 | 10 | R01(0.2) | G>A | TFBS_Disruption | RAD2 |
| DI | VIII | 66,694 | 10 | R01(0.4) | A>G | Synonymous | OPI1 |
| DI | VIII | 266,228 | 10 | R12(0.2) | C>T | Missense | YHR080C |
| DI | VIII | 266,233 | 10 | R12(0.2) | C>T | Synonymous | YHR080C |
| DI | IX | 139,397 | 10 | R12(0.4) | G>C | Intergenic | RHO3 |
| DI | IX | 155,254 | 10 | R12(0.2) | A>T | Intergenic | COX5B |
| DI | IX | 204,002 | 10 | R12(0.2) | G>A | Missense | CAB2 |
| DI | IX | 210,821 | 10 | R08(0.2) | C>T | TFBS_Disruption | tl(AAU)I2 |
| DI | X | 436,041 | 10 | R12(0.2) | T>C | TFBS_Disruption | AVT1 |
| DI | X | 533,912 | 10 | R12(0.2) | T>A | TFBS_Disruption | BFA1 |
| DI | X | 636,907 | 10 | R05(0.2) | C>T | TFBS_Disruption | YJR111C |
| DI | X | 738,407 | 10 | R10(0.3) | T>A | Missense | MPH3 |
| DI | XI | 581,078 | 10 | R04(0.2) | G>A | TFBS_Disruption | YKR075C |
| DI | XII | 545,956 | 10 | R01(0.2) | A>T | TFBS_Disruption | NOP56 |
| DI | XII | 751,305 | 10 | R05(0.2) | A>T | Missense | IMH1 |
| DI | XIII | 58,787 | 10 | R01(0.3) | A>T | TFBS_Disruption | SEC65 |
| DI | XIII | 559,094 | 10 | R16(0.2) | G>A | Intergenic | TIF34 |
| DI | XIII | 822,630 | 10 | R08(0.2) | A>T | TFBS_Disruption | PRM15 |
| DI | XIV | 79,576 | 10 | R12(0.2) | G>A | Missense | RIM21 |
| DI | XIV | 551,173 | 10 | R01(0.2) | A>T | Missense | COG6 |
| DI | XIV | 551,174 | 10 | R01(0.2) | G>T | Missense | COG6 |
| DI | XV | 908,360 | 10 | R01(0.2) | G>T | Intergenic | ARS1528 |
| DI | XV | 930,496 | 10 | R05(0.2) | G>A | TFBS_Disruption | MYO2 |
| DI | XV | 958,391 | 10 | R12(0.2) | C>T | Missense | UBC11 |
| DI | XVI | 868,344 | 10 | R12(0.3) | T>C | TFBS_Disruption | ORC4 |
| DI | M | 43,971 | 10 | R05(0.3) | T>A | TFBS_Disruption | COB-OLI1 |
| GA | I | 662 | 9 | R07(0.3) | C>T | Missense | YAL068W-A |
| GA | I | 42,085 | 9 | R09(0.4) | G>A | Intergenic | ARS105 |
| GA | II | 7,420 | 9 | R09(0.3) | C>T | TFBS_Disruption | YBL108W |
| GA | II | 28,453 | 9 | R11(0.2) | G>T | Synonymous | YBL100W-C |
| GA | II | 537,692 | 9 | R11(0.2) | C>A | Intergenic | YSW1 |
| GA | V | 141,253 | 9 | R16(0.2) | T>A | Intergenic | YEL008C-A |
| GA | VI | 107,628 | 9 | R06(0.2) | A>T | TFBS_Disruption | YFL015C |
| GA | VII | 121,047 | 9 | R09(0.2) | T>A | Intergenic | MCM6 |
| GA | VIII | 324,276 | 9 | R16(0.2) | G>A | Missense | GRE3 |
| GA | VIII | 370,610 | 9 | R09(0.3) | A>T | TFBS_Disruption | ECM14 |
| GA | IX | 231,393 | 9 | R16(0.2) | T>A | Intergenic | MAM33 |
| GA | IX | 310,742 | 9 | R16(0.3) | G>T | TFBS_Disruption | YKE4 |
| GA | IX | 391,333 | 9 | R07(0.6) | A>G | Synonymous | FLO11 |
| GA | IX | 407,727 | 9 | R05(0.8) | A>G | TFBS_Disruption | DAL1 |
| GA | X | 461,703 | 9 | R16(0.3) | T>A | TFBS_Disruption | TMA22 |
| GA | X | 537,895 | 9 | R16(0.2) | C>T | Intergenic | YJRCdelta16 |
| GA | X | 580,729 | 9 | R16(0.2) | G>T | Premature_stop | AIM24 |
| GA | XI | 219,865 | 9 | R16(0.7) | A>C | Intergenic | VPH2 |
| GA | XI | 635,512 | 9 | R11(0.2) | G>T | Intergenic | UBP11 |
| GA | XII | 11,774 | 9 | R04(0.2) | C>T | Missense | YLL065W |
| GA | XIII | 163,384 | 9 | R16(0.3) | C>A | Intergenic | IMD4 |
| GA | XIV | 753,650 | 9 | R16(0.2) | G>T | Missense | YNR065C |
| GA | XV | 21,713 | 9 | R09(0.2) | T>A | Intergenic | ENB1 |
| GA | XV | 444,518 | 9 | R04(0.2) | A>T | TFBS_Disruption | YOR062C |
| GA | XV | 470,717 | 9 | R05(0.2) | G>T | Missense | SKI7 |
| GA | XV | 711,067 | 9 | R16(0.2) | G>T | Missense | PEX27 |
| GA | XV | 711,068 | 9 | R16(0.2) | A>T | Missense | PEX27 |
| GA | XV | 908,360 | 9 | R09(0.3) | G>T | Intergenic | ARS1528 |
| GA | XV | 908,361 | 9 | R09(0.2) | G>T | Intergenic | ARS1528 |
| GA | XV | 1,079,760 | 9 | R05(0.3) | G>A | Intergenic | HSP33 |
| GA | XVI | 198,563 | 9 | R16(0.2) | C>A | Intergenic | MRN1 |
| NC | I | 105,416 | 10 | R09(0.2) | G>T | Missense | LTE1 |
| NC | II | 407,753 | 10 | R05(0.2) | C>T | TFBS_Disruption | UBC4 |
| NC | II | 477,371 | 10 | R12(0.3) | C>A | TFBS_Disruption | TKL2 |
| NC | III | 150,045 | 10 | R04(0.7) | C>T | Intergenic | tM(CAU)C |
| NC | IV | 99,505 | 10 | R04(0.3) | T>G | Intergenic | TRM8 |
| NC | IV | 169,868 | 10 | R06(0.2) | T>A | Intergenic | MHF2 |
| NC | IV | 551,805 | 10 | R06(0.3) | G>A | TFBS_Disruption | HEM12 |
| NC | IV | 688,415 | 10 | R04(0.4) | A>G | Synonymous | VBA4 |
| NC | IV | 1,254,549 | 10 | R01(0.2) | G>T | Intergenic | SAC7 |
| NC | IV | 1,523,881 | 10 | R06(0.2) | A>C | Intergenic | PAU10-YDR543C |
| NC | V | 86,707 | 10 | R09(0.2) | A>T | Intergenic | YEL034C-A |
| NC | V | 148,091 | 10 | R03(0.3) | G>A | TFBS_Disruption | GIM4 |
| NC | V | 274,150 | 10 | R05(0.3) | T>A | Intergenic | FCY21 |
| NC | VI | 4,981 | 10 | R12(0.3) | G>A | Intergenic | YFL063W |
| NC | VII | 627,235 | 10 | R06(0.2) | C>T | Synonymous | YGR069W |
| NC | VII | 784,148 | 10 | R02(0.2) | G>A | Intergenic | ECL1 |
| NC | VII | 965,741 | 10 | R04(0.2) | A>T | TFBS_Disruption | YGR237C |
| NC | VII | 1,072,826 | 10 | R09(0.2) | C>A | TFBS_Disruption | MAL13-MAL11 |
| NC | VIII | 456,255 | 10 | R03(0.2) | C>A | Intergenic | YHR177W |
| NC | VIII | 528,898 | 10 | R01(0.5) | G>A | Intergenic | FLO5-YHR212C |
| NC | IX | 241,023 | 10 | R07(0.2) | G>A | Intergenic | RNR3 |
| NC | X | 90,461 | 10 | R12(0.5) | A>C | TFBS_Disruption | ATG27 |
| NC | X | 196,100 | 10 | R06(0.3),<br>R12(0.3) | A>T | TFBS_Disruption | ASF1 |
| NC | X | 389,869 | 10 | R04(0.2) | A>T | Missense | VPS53 |
| NC | X | 525,764 | 10 | R03(0.3) | T>A | TFBS_Disruption | ANB1 |
| NC | X | 586,416 | 10 | R05(0.2) | C>A | Missense | YJR087W |
| NC | XI | 1,436 | 10 | R04(0.3) | A>G | Intergenic | YKL223W |
| NC | XI | 255,762 | 10 | R04(0.5) | T>C | Synonymous | UTP11 |
| NC | XII | 51,985 | 10 | R11(0.3) | G>A | Intergenic | FPS1 |
| NC | XII | 86,696 | 10 | R12(0.2) | G>A | TFBS_Disruption | ISA1 |
| NC | XII | 97,693 | 10 | R04(0.7) | G>A | TFBS_Disruption | SSA2 |
| NC | XII | 150,971 | 10 | R09(0.3),<br>R11(0.5) | A>T | Intergenic | ARS1208 |
| NC | XII | 150,974 | 10 | R09(0.2),<br>R11(0.4) | A>T | Intergenic | ARS1208 |
| NC | XII | 683,211 | 10 | R07(0.3) | G>A | Missense | YLR271W |
| NC | XII | 845,918 | 10 | R04(0.2) | G>T | Intergenic | VPS38 |
| NC | XII | 875,494 | 10 | R01(0.2) | C>T | TFBS_Disruption | FBP1 |
| NC | XIII | 234,792 | 10 | R12(0.3) | C>T | Synonymous | YML018C |
| NC | XIII | 276,372 | 10 | R12(0.2) | G>A | Missense | TAF4 |
| NC | XIII | 371,021 | 10 | R04(0.2) | A>T | Intergenic | ARS1312 |
| NC | XIII | 775,210 | 10 | R11(0.2) | C>T | TFBS_Disruption | YMR252C |
| NC | XIII | 822,630 | 10 | R12(0.3) | A>T | TFBS_Disruption | PRM15 |
| NC | XIII | 892,720 | 10 | R02(0.2) | G>A | TFBS_Disruption | PSE1 |
| NC | XIV | 172,410 | 10 | R04(0.3) | G>A | Intergenic | MRPL17 |
| NC | XIV | 172,411 | 10 | R04(0.2) | G>A | TFBS_Disruption | MRPL17 |
| NC | XIV | 174,753 | 10 | R04(0.2) | G>T | TFBS_Disruption | NRD1 |
| NC | XIV | 191,318 | 10 | R02(0.2) | G>T | Intergenic | ATG2 |
| NC | XIV | 573,696 | 10 | R03(0.2) | A>T | Missense | YNL033W |
| NC | XIV | 690,093 | 10 | R04(0.2) | G>T | TFBS_Disruption | SOL1 |
| NC | XIV | 758,691 | 10 | R09(0.6) | G>C | Missense | DSE4 |
| NC | XV | 243,362 | 10 | R09(0.3) | C>T | TFBS_Disruption | LDS2 |
| NC | XV | 348,674 | 10 | R09(0.2) | C>A | TFBS_Disruption | TIR2-AUS1 |
| NC | XV | 759,543 | 10 | R05(0.2) | A>T | Intergenic | snR35 |
| NC | XVI | 352,487 | 10 | R04(0.2) | A>T | TFBS_Disruption | SSE1 |
| NC | XVI | 381,201 | 10 | R06(0.2) | A>T | TFBS_Disruption | RLM1 |
| NC | XVI | 835,344 | 10 | R09(0.2) | C>T | Intergenic | PIN3 |
| NC | M | 37,760 | 10 | R11(0.6) | A>G | Missense | BI2 |
| UR | II | 347,317 | 6 | R09(0.3) | C>T | TFBS_Disruption | PRP6 |
| UR | IV | 210,699 | 6 | R05(0.2) | G>T | TFBS_Disruption | RPO21 |
| UR | IV | 561,086 | 6 | R09(0.3) | G>T | Intergenic | DBF4 |
| UR | IV | 1,061,955 | 6 | R09(0.2) | C>T | Synonymous | PRO1 |
| UR | VII | 717,188 | 6 | R09(0.2) | C>T | TFBS_Disruption | SHY1 |
| UR | VII | 1,063,279 | 6 | R05(0.2) | C>A | Intergenic | ZUO1 |
| UR | X | 151,178 | 6 | R09(0.2) | C>A | Premature_stop | RPB4 |
| UR | X | 579,823 | 6 | R09(0.2) | G>A | Missense | BNA2 |
| UR | X | 716,443 | 6 | R09(0.3) | C>T | Intergenic | DAN4 |
| UR | X | 716,444 | 6 | R09(0.3) | C>T | Intergenic | DAN4 |
| UR | X | 716,445 | 6 | R09(0.2) | C>T | Intergenic | DAN4 |
| UR | XI | 160,597 | 6 | R09(0.2) | A>T | Missense | RSM22 |
| UR | XI | 321,880 | 6 | R09(0.3) | A>T | Intergenic | YKL063C |
| UR | XI | 398,709 | 6 | R08(0.3) | C>A | Missense | MAK11 |
| UR | XII | 517,858 | 6 | R05(0.2) | G>A | Intergenic | VTA1 |
| UR | XII | 751,305 | 6 | R09(0.2) | A>T | Missense | IMH1 |
| UR | XII | 911,891 | 6 | R09(0.3) | A>T | Missense | VPS33 |
| UR | XIII | 5,743 | 6 | R05(0.4) | T>C | TFBS_Disruption | YML133C-COS3 |
| UR | XIII | 26,589 | 6 | R15(1) | A>T | TFBS_Disruption | PHO84 |
| UR | XIII | 772,685 | 6 | R05(0.2) | C>T | Intergenic | ARS1328 |
| UR | XIV | 552,709 | 6 | R09(0.3) | G>A | Intergenic | COG6 |
| UR | XIV | 782,992 | 6 | R08(0.2) | G>A | TFBS_Disruption | PAU6-YNR077C |
| UR | XVI | 17,238 | 6 | R09(0.2) | A>T | Intergenic | YPL277C |
| UR | XVI | 469,773 | 6 | R09(0.3) | G>A | TFBS_Disruption | NOP4 |
| UR | XVI | 774,801 | 6 | R09(0.2) | C>T | Missense | CLB5 |
| YP | III | 173,565 | 5 | R02(0.7) | T>C | TFBS_Disruption | RIM1 |
| YP | III | 302,896 | 5 | R08(0.3) | G>C | Synonymous | YCR101C |
| YP | IV | 163,869 | 5 | R03(0.5) | C>T | Synonymous | FAP7 |
| YP | IV | 212,555 | 5 | R08(0.3) | C>T | Intergenic | ARS409 |
| YP | IV | 838,154 | 5 | R04(0.2) | C>T | TFBS_Disruption | SLY1 |
| YP | IV | 838,155 | 5 | R04(0.2) | A>T | TFBS_Disruption | SLY1 |
| YP | IV | 838,156 | 5 | R04(0.2) | G>T | Intergenic | SLY1 |
| YP | IV | 1,523,881 | 5 | R08(0.4) | A>C | Intergenic | PAU10-YDR543C |
| YP | IX | 217,788 | 5 | R03(0.3) | A>T | TFBS_Disruption | RPN2 |
| YP | X | 524,681 | 5 | R08(0.2) | C>A | TFBS_Disruption | tS(AGA)J |
| YP | XI | 266,177 | 5 | R04(0.3) | A>T | Missense | BUD2 |
| YP | XIII | 420,846 | 5 | R03(0.3) | A>T | Intergenic | PDS5 |
| YP | XIII | 420,847 | 5 | R03(0.3) | A>T | Intergenic | PDS5 |
| YP | XIII | 865,417 | 5 | R08(0.2) | C>T | TFBS_Disruption | DYN3 |
| YP | XIV | 118,963 | 5 | R02(0.3) | G>T | Intergenic | YNL276C |
| YP | XIV | 576,646 | 5 | R03(0.2),<br>R04(0.3) | C>T | TFBS_Disruption | HHT2 |
| YP | XV | 1,048,427 | 5 | R04(0.7) | T>C | Intergenic | ATF1-AMF1 |

**Table S5.** Percent of genome with a LOD score greater than 5 per chemical

| Chemical | LOD > 5 |
| --- | --- |
| cadmium chloride | 98.6 |
| chlorpromazine | 100 |
| diamide | 90.5 |
| glacial acetic acid | 98.2 |
| sodium chloride | 92.2 |
| urea | 88.4 |
| YPD | 99.9 |

**Table S6.**
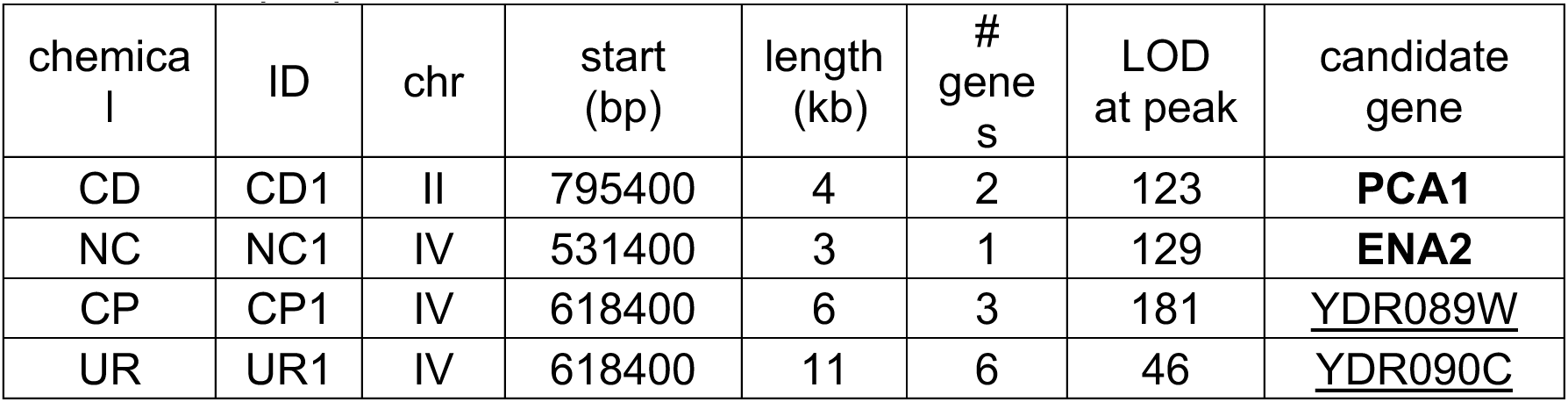
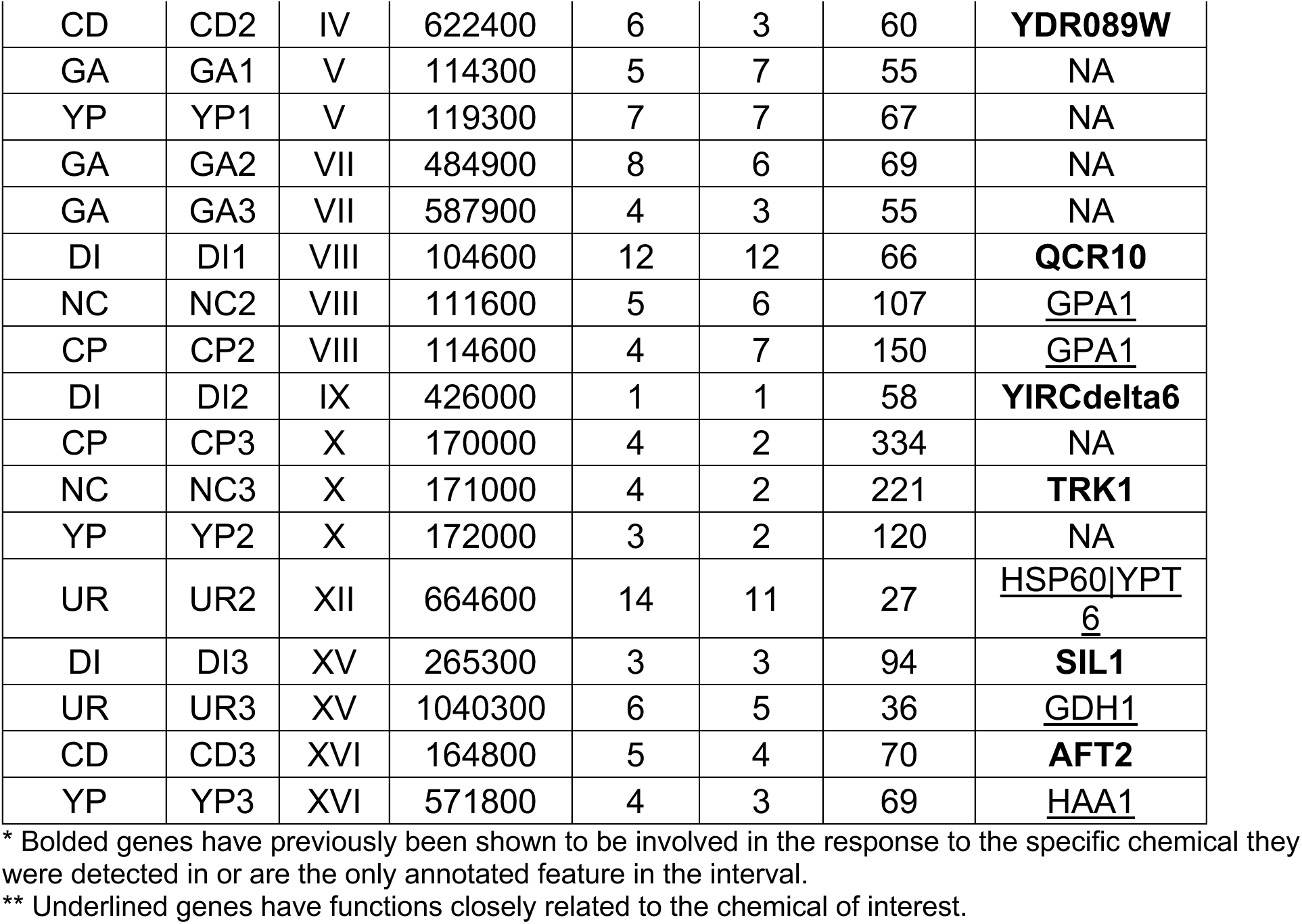
Top 3 peaks detected in each chemical treatment.

| chemical | ID | chr | start (bp) | length (kb) | # genes | LOD at peak | candidate gene |
| --- | --- | --- | --- | --- | --- | --- | --- |
| CD | CD1 | II | 795400 | 4 | 2 | 123 | **PCA1** |
| NC | NC1 | IV | 531400 | 3 | 1 | 129 | **ENA2** |
| CP | CP1 | IV | 618400 | 6 | 3 | 181 | <u>YDR089W</u> |
| UR | UR1 | IV | 618400 | 11 | 6 | 46 | <u>YDR090C</u> |
| CD | CD2 | IV | 622400 | 6 | 3 | 60 | **YDR089W** |
| GA | GA1 | V | 114300 | 5 | 7 | 55 | NA |
| YP | YP1 | V | 119300 | 7 | 7 | 67 | NA |
| GA | GA2 | VII | 484900 | 8 | 6 | 69 | NA |
| GA | GA3 | VII | 587900 | 4 | 3 | 55 | NA |
| DI | DI1 | VIII | 104600 | 12 | 12 | 66 | **QCR10** |
| NC | NC2 | VIII | 111600 | 5 | 6 | 107 | <u>GPA1</u> |
| CP | CP2 | VIII | 114600 | 4 | 7 | 150 | <u>GPA1</u> |
| DI | DI2 | IX | 426000 | 1 | 1 | 58 | **YIRCdelta6** |
| CP | CP3 | X | 170000 | 4 | 2 | 334 | NA |
| NC | NC3 | X | 171000 | 4 | 2 | 221 | **TRK1** |
| YP | YP2 | X | 172000 | 3 | 2 | 120 | NA |
| UR | UR2 | XII | 664600 | 14 | 11 | 27 | <u>HSP60\|YPT6</u> |
| DI | DI3 | XV | 265300 | 3 | 3 | 94 | **SIL1** |
| UR | UR3 | XV | 1040300 | 6 | 5 | 36 | <u>GDH1</u> |
| CD | CD3 | XVI | 164800 | 5 | 4 | 70 | **AFT2** |
| YP | YP3 | XVI | 571800 | 4 | 3 | 69 | <u>HAA1</u> |

\* Bolded genes have previously been shown to be involved in the response to the specific chemical they were detected in or are the only annotated feature in the interval.
\*\* Underlined genes have functions closely related to the chemical of interest.

**Table S7.** Potentially pleiotropic regions

| ID | chemicals | chr | start | interval (kb) | genes |
| --- | --- | --- | --- | --- | --- |
| PL1 | GA, NC | IV | 503400 | 16 | MIX14\|PST2\|ARS450\|MRH1\|LYS14\|YDRCdelta2\|YDRCdelta3\|YDRCdelta4\|YDRCTy2-1\|YDR034C-D\|YDR034C-C\|YDRCdelta5 |
| PL2 | CD, CP, DI, NC, UR | IV | 618400 | 8 | SLU7\|tR(ACG)D\|YDR089W\|YDR090C |
| PL3 | GA, YP | V | 119300 | 7 | YEL020C\|MMS21\|YEL018C-A\|EAF5\|PMP2\|GTT3\|NPP2 |
| PL4 | DI, GA, NC, YP | VII | 499900 | 3 | SWC4\|CUL3 |
| PL5 | CP, DI, NC | VIII | 101600 | 18 | LAG1\|YHL002C-A\|HSE1\|RPL14B\|CEN8\|OSH7\|QCR10\|LEU5\|TCD1\|NEM1\|GPA1\|TIM10\|tT(AGU)H\|ARS806\|YHRCdelta3\|YHRCdelta4\|STP2 |
| PL6 | CP, NC,<br>YP | X | 162000 | 17 | YJL132W\|AIM23\|URA2\|TRK1\|PBS2 |
| PL7 | CP, NC,<br>YP | XVI | 437800 | 24 | YPLCTy4-1\|YPL060C-A\|YPLCtau2\|<br>GRX5\|PDR12\|SUR1\|LCL1\|LGE1\|<br>LEE1\|ARS1633\|KTR6\|OAZ1\|ARL3\|<br>MNN9 |

## Supplemental Results

### Candidate genes and candidate causative variants for the 21 leading factors

To identify major effect candidate genes and/or potentially causative variants, we examined the top three peaks from each condition, and for each peak define a 2.5-LOD support interval. Across all ten treatments, the average size of the 2.5-LOD support interval was 6kb and contained on average 5 genes (Table S6).

Under eight of the twenty-one total peaks examined (CD1, CD2, CD3, DI1, DI2, DI3, NC1, and NC3), we identified a single gene (*PCA1, VTC5, AFT2, QCR10, YIRCdelta6, SIL1, ENA2,* and *TRK1*, respectively; bolded in Table S6) that has previously been either shown *a priori* to respond to the same chemical stressor as used in this study or to be the only annotated feature under the peak. Based on an *a posteriori* examination of the function of each gene under the remaining 13 peaks, 6 (CP1, CP2, NC2, UR1, UR3, and YP3) had a single candidate gene (*VTC5, GPA1, GPA1, YDR090C* (*ILT1*), *GDH1*, and *HAA1*, respectively) and 1 (UR2) had two candidate genes (*HSP60* and *YPT6*) with functions closely related to the chemical challenge (underlined in Table S6). The remaining six peaks did not have strong *a priori* or *a posteriori* candidates. Detailed below is evidence that makes the genes listed above strong candidates for subsequent follow-up studies as well as potentially causal variants that could be functionally validated in future work across all 21 peaks.

Cadmium chloride tolerance has previously been shown to be mediated in large part by *PCA1*, a cadmium transporter that has been detected in the same chemical treatment in a previous X-QTL study (Ehrenreich et al., 2010) and that is known to confer tolerance to cadmium chloride treatment (Shiraishi et al., 2000). A moderately sized in-frame insertion occurs in *PCA1* in four founder strains, including the Most Increased Haplotype (MIH), B8, which has a 36bp insertion towards the beginning of the protein, as does AB4 and B9, while A5 has a 72bp insertion. B8 is further distinguished from AB4 and B9 by 6 nonsynonymous SNPs, 4 of which are predicted to change the secondary structure of the protein. Previous work has shown that deletion of the candidate gene *YDR089W* (*VTC5*) leads to increased cadmium chloride resistance through an unknown mechanism (Ruotolo et al., 2008). *VTC5* may influence heavy metal tolerance through its’ regulation of the production of polyphosphate molecules (Desfougères et al., 2016), which have been found to be involved in resistance to manganese and cadmium and are present in the vacuole of *S. cerevisiae* (Andreeva et al., 2013; Trilisenko et al., 2017) and can increase tolerance to cadmium in bacteria (Keasling & Hupf, 1996). AB4, the MIH for this region, has a single nonsynonymous SNP in a highly conserved region predicted to change the secondary structure of the protein. This SNP is shared by founder A7- however, A7 is at the lower limit of our ability to detect haplotypes and may not be present at this locus. Deletion of *AFT2,* an iron-regulated transcription factor, has been shown to sensitize yeast to cadmium exposure, possibly due to cadmium interfering with iron homeostasis (Ruotolo et al., 2008; Thorsen et al., 2009). A variant private to the MIH establishes a *YAP1* binding site upstream of the TSS in *AFT2* (Figure 4). *YAP1* itself has previously been shown to induce the expression of several genes involved in tolerance to cadmium chloride (Wemmie et al., 1994).

Chlorpromazine is known to have a multitude of effects in budding yeast, including inducing oxidative stress, altering membrane integrity, decreasing cellular lipid levels (Muhieddine et al., 2019), and inhibiting intracellular protein trafficking (Nierras & Warner, 1999). *VTC5* (Figure 4- figure supplement 1A) is also a strong candidate gene for this drug, as a reduced intracellular pH has been shown to interfere with chlorpromazine’s toxicity (Ahyayauch et al., 2003), and *VTC5* has a role in maintaining the pH of the vacuole (Desfougères et al., 2016). The MIH for this region is the same as for the same region detected with cadmium chloride treatment. Another strong candidate is *GPA1*, is a gene with a known role in the response to mating pheromone (Miyajima et al., 1987) as well as involvement in the MAPK pathway (Metodiev et al., 2002), which interacts with a Phosphatidylinositol 3-Kinase to increase cellular levels of phosphatidylinositol 3-Phosphate. The MIH, AB4, has a coding SNP at a highly conserved location predicted to change the secondary structure of the protein. This SNP is shared with founder B5- both founder haplotypes increase significantly in frequency at this locus, with AB4 increasing ∼16-fold and B5 increasing by ∼2.5-fold as compared to initial frequencies in the base population. CP3 contains only two annotated features, the genes *URA2* and *TRK1*. Ura2p is a bifunctional protein that catalyzes the first two steps in the *de novo* biosynthesis of pyrimidines (Lue & Kaplan, 1969; Souciet et al., 1982). The MIH, AB4 has two nonsynonymous SNPs (one of which is predicted to change the secondary structure of the protein) and three silent SNPs- all five of these coding SNPs are in highly conserved regions. *TRK1* is a high-affinity potassium transporter (Gaber et al., 1988). The MIH, AB4, has two insertions that lead to the addition of an aspartic acid and two more lysine residues in the CDS which the rest of the founders lack. AB4 also has 3 nonsynonymous SNPs, all predicted to change the secondary structure of the protein, and 2 synonymous SNPs, one of which lies in a highly conserved region.

For diamide tolerance, deletion of the candidate gene *SIL1* (Figure 4- figure supplement 1B) leads to increased resistance (Siegenthaler et al., 2017). *SIL1* has been shown to affect protein folding in the ER by reducing a known *HSP70* chaperone, *KAR2* (Wang et al., 2014). The MIH, B5, has two nonsynonymous SNPs, one of which is predicted to change the secondary structure of the protein and is in a highly conserved region. Deletion of *QCR10,* a subunit of the mitochondrial respiratory chain, has also been shown to confer sensitivity to diamide treatment (Thorpe et al., 2004). B5, the MIH, has a single private SNP upstream of the TSS that may establish a *CST6* binding site which the other founders lack. *CST6* has been shown to be involved in oxidative stress tolerance (Liu et al., 2016). The last candidate gene, *YIRCdelta6*, is a Ty1 retrotransposon that is present in all founders; however, the MIH, A5, has a SNP which deletes a downstream binding site for Stb5p. Deletion of this gene has previously been shown to specifically decrease levels of Ty1 transposition (Griffith et al., 2003).

One of the peaks detected for glacial acetic acid tolerance includes *URA3*, a gene that was knocked out in all founder strains to enable subsequent genetic manipulations. The deletion inserted a kanamycin resistance cassette at this locus. As kanamycin was used at high concentrations to prevent bacterial contamination in all cultures, we speculate that a *cis* factor or a mutation introduced during the deletion resulted in higher levels of kanamycin expression in one of founders AB2, A7, or B12 (these 3 founders are indistinguishable in this region) and perhaps higher growth rates. Alternatively, the MIH at this locus has a binding site for Msn2p just upstream of the TSS of *GEA2*- this site is missing in the remaining founders. Msn2p is a transcriptional activator known to be involved in the response to acid stress (Causton et al., 2001; Mira et al., 2010). Only a single gene under GA2, *RPN14*, has a private coding SNP- this SNP is predicted to change the secondary structure of the protein. *RPN14* is a protein folding chaperone that helps to assemble the 19S regulatory particle of the proteasome (Roelofs et al., 2009; Saeki et al., 2009). GA3 has two genes with potentially functional variation, including *TFC4*, a subunit of the RNA polymerase III transcription initiation factor complex, and *SCM4*, a mitochondrial outer membrane protein of unknown function. *SCM4* has a single synonymous SNP in a highly conserved region while *TFC4* has one nonsynonymous SNP predicted to change the secondary structure of the protein and five synonymous SNPs - the nonsynonymous SNP and three of the synonymous SNPs are in highly conserved regions.

For salt tolerance, Ena2p is a well-known P-ATPase sodium pump involved in sodium efflux (Haro et al., 1991)(Figure 4- figure supplement 2A). The *ENA* gene region contains three genes *ENA1*, *ENA2*, and *ENA5*, with our base population segregating a complex structural variant. In 10 of 18 of the founders the *ENA* locus has been inverted with only the *ENA1* gene present, while the other founders have all three known *ENA* genes present in the reference orientation. Counter-intuitively, one of the founders with only *ENA1* (AB4) is the most increased in frequency for this candidate region, so the lack of *ENA2* and *ENA5* alone is not sufficient to explain adaptation. A potential explanation lies in the fact that the AB4 founder additionally lacks a predicted binding site for the transcriptional repressor (Mig1p), which is known to repress genes in the presence of glucose (Nehlin & Ronne, 1990)- only one other founder (A5) lacks this binding site. If this hypothesis is correct the allele responding to selection is due to an epistatic interaction between two mutations. Finally, there is a nonsynonymous SNP in *ENA1* that distinguishes founders AB4 and A5, but it is unclear if this SNP in concert with the loss of a Mig1p binding site can explain the haplotype change. Another candidate gene for sodium chloride tolerance is *TRK1* (described above), deletion of which leads to increased sensitivity to sodium ions, potentially due to the ability of Trk1p to discriminate between potassium and sodium ions (Haro et al., 1993). The MIH is the same as that detected in chlorpromazine treatment. *GPA1* has also been predicted to have a role in the cellular response to high osmolarity (Hohmann, 2002) and shares the same MIH as that detected in chlorpromazine treatment.

Nitrogen catabolite repression (NCR) occurs when a preferred nitrogen source is present in the media (such as ammonium and asparagine), leading to the repression of genes involved in the utilization of non-preferred nitrogen sources (such as proline and urea). In the presence of a preferred nitrogen source, the transcription factors Gat1p and Gln3p are sequestered in the cytoplasm by Ure2p. Once a preferred nitrogen source has become limiting, Gln3p and Gat1p are dephosphorylated and localize to the nucleus, where they activate transcription of genes needed to use non-preferred nitrogen sources (reviewed in (Hofman-Bang, 1999)). Two of the urea resistance peaks (UR1 and UR3) contain genes (*ILT1* and *GDH1*, respectively) shown to have transcriptional profiles similar to other known NCR genes when exposed to different nitrogen sources (Boer et al., 2007), making them good candidates for this trait. *GDH1* (Figure 4- figure supplement 2B) is known to be involved in the ammonia assimilation cycle (Grenson et al., 1974) and the most changed haplotype, AB4, has a nonsynonymous SNP predicted to change the secondary structure of the protein that none of the other founders have. The MIH at *ILT1* has an Hsf1p binding site (Hsf1p responds to highly diverse stressors) and is missing a Gat1p binding site the other founders all have, which may be beneficial if Ilt1p, an integral membrane protein, is involved in the uptake of urea. UR2 contains a heat shock protein, *HSP60*, that may play a role in stabilizing proteins in response to the strong denaturing action of urea, as well as *YPT6*, a gene whose deletion leads to the impaired utilization of a variety of nitrogen sources (VanderSluis et al., 2014). There is a single synonymous SNP at *HSP60* private to the MIH at a highly conserved region and three downstream SNPs, one of which is also at a highly conserved sequence. *YPT6* has a single nonsynonymous SNP private to the MIH at a highly conserved region and a single downstream SNP. There is also a trinucleotide repeat (AAC) downstream of *YPT6* that is amplified by at least 6 repeats in the MIH. This particular repeat has previously been shown to be overrepresented in budding yeast ORFs (Katti et al., 2001), which hints that there may be an unannotated ORF present in this region.

For growth in rich media, there are no clear-cut candidates within YP1, although there are abundant potentially functional private variants in the MIH, including 6 nonsynonymous SNPs (all of which are predicted to change the secondary structure of their cognate protein) and 6 synonymous SNPs. YP2 contains the previously described *URA2* and *TRK1* genes, with the same MIH as detected in chlorpromazine and sodium chloride treatment. The final YPD peak has a Ty1 LTR with a single SNP private to the MIH as well as a good candidate gene *HAA1*, a transcriptional activator involved in adaptation to weak acid stress (cultures of budding yeast are known to accumulate acetic acid over time) (Fernandes et al., 2005). The MIH at *HAA1* has one nonsynonymous and one synonymous SNP, both of which are at highly conserved sites.

### Candidate genes and candidate causative variants for the regions exhibiting pleiotropy

Despite pleiotropy not being universal there were several regions of the genome that show strong across treatment correlations in LOD scores for subsets of chemicals (Figure 8). To identify the specific genomic regions exhibiting pleiotropy we calculated z-scores from the log-transformed LOD scores and regions having z-scores >=1.96 for at least two drugs were plotted (Figure 9- figure supplement 1). There are seven such strong regions genome-wide (Figure 9- figure supplement 2), including: two regions on chromosome IV (one shared between glacial acetic acid and sodium chloride, the other shared between urea, chlorpromazine, sodium chloride, diamide, and cadmium chloride), a region on chromosome V (shared between glacial acetic acid and YPD), a region on chromosome VII (shared between YPD, diamide, glacial acetic acid, and sodium chloride), a region on chromosome VIII (shared between chlorpromazine, sodium chloride, and diamide), a region on chromosome X (shared between chlorpromazine, sodium chloride, and YPD; see Figure 9), and a region on chromosome XVI (shared between YPD, sodium chloride, and chlorpromazine). For all seven potentially pleiotropic regions, Table S7 shows genes contained within the overlapping peaks shown in Figure 9- figure supplement 2.

For PL1, the MIH is different for glacial acetic acid and sodium chloride, which hints that different genes within this region may be causal. *MRH1* is a good candidate for glacial acetic acid tolerance as deletion has been shown to confer sensitivity to this stressor (Takabatake et al., 2015)- there is a single synonymous SNP private to the MIH at a highly conserved region. Alternatively, *LYS14*, which regulates lysine biosynthesis, is a good candidate for sodium chloride tolerance as the MIH has 3 nonsynonymous SNPs, 2 of which are predicted to change the secondary structure of the protein and 2 of which are highly conserved, 3 synonymous SNPs, 2 of which are also highly conserved, and a *MSN2* binding site the other founders lack. Msn2p is a transcriptional activator known to be involved in the response to multiple different stressors (Gasch et al., 2000). For PL2, diamide has a different MIH than the other four chemicals with peaks detected in this region (all of which have AB4 as the MIH). At this locus, the most likely candidate is *VTC5*, which, as described above, is a candidate gene for both cadmium chloride and chlorpromazine tolerance. For PL3, the most likely candidate is *PMP2*, a gene that previously has been shown to be involved in acetic acid tolerance (Mira et al., 2010) and that regulates the activity of Pma1p (Navarre & Goffeau, 1998), which is an H(+)- ATPase at the plasmid membrane. At PL4, the MIH for diamide and glacial acetic acid is AB3, while the MIH for sodium chloride and YPD is B9. Only two genes are contained within the detected interval- *SWC4*, a component of a histone acetyltransferase complex, and *CUL3*, an E3 ubiquitin ligase involved in RNA polymerase II degradation. In founder B9, there is a nonsynonymous SNP at a highly conserved sequence in both *SWC4* and *CUL3*, while founder AB3 has a single SNP in the promoters of both *SWC4* and *CUL3*, with a single synonymous SNP in *CUL3* at a highly conserved sequence.

For the three chemicals detected in PL5, the MIH for diamide is B5, while the MIH for chlorpromazine and sodium chloride is AB4. Good candidate genes for these drugs include *QCR10* for diamide and *GPA1* for chlorpromazine and sodium chloride, as described above. For PL6, *TRK1, URA2,* and *PBS2* are all good candidates- all harbor multiple SNPs private to AB4. As described above, Ura2p is part of the *de novo* pyrimidine biosynthesis pathway, while Trk1p is a potassium transporter. It is possible that the *de novo* synthesis of pyrimidine provides a benefit under these specific conditions, although, as *TRK1* has previously been shown to affect sensitivity to sodium ions, this may be the better candidate. Pbs2p mutants have previously been shown to have a defective cell wall and increased susceptibility to hyperosmotic stress (Gopalbhai et al., 2003), which makes it a good candidate for both chlorpromazine and sodium chloride tolerance, respectively. For the final pleiotropic peak, PL7, there are three candidate genes, including *GRX5, PDR12,* and *SUR1*. *GRX5* has previously been shown to be involved in the oxidative stress and osmotic stress response (Chabrier-Roselló et al., 2010; Chakrabortee et al., 2016), making it a good candidate for all three drugs (chlorpromazine, sodium chloride, and YPD). *PDR12* is a multidrug transporter, while *SUR1* has also been shown to be involved in the oxidative stress response (Helsen et al., 2020) and is required for the biosynthesis of sphingolipids (reviewed in (Dickson & Lester, 1999)). The MIH at this region is AB4, which, at *GRX5,* has only a single SNP in the promoter region, while at *PDR12* there are two synonymous SNPs, both in highly conserved sequences, while at *SUR1* there is a single synonymous SNP at a highly conserved sequence.

## Supplemental Notes

**Note S1.** We discovered that there were two classes of potential ‘cheaters’ that had emerged during evolution. One, the aneuploid haploids, turned out to be haploid at all but the mating type locus. Upon closer examination of sequencing coverage, we found evidence that either a nondisjunction event occurred during meiosis causing chromosomes harboring both mating type loci (a/@) to be retained in an otherwise haploid background or a recombination event occurred in which the entirety of one mating type locus inserted into the opposite mating type locus. Either situation allowed these cells to escape diploid selection, as the markers used to select diploids were tightly linked to the mating type locus. Further, these haploid cheaters seemed as resistant to lysis as actual spores; we later found that only treatment with ether (which cannot be done safely in high throughput) can kill these cells without killing off all the spores as well (treatment with heat shock did not seem to work either, which would have been more amenable to high throughput). The second type of cheater looks like a single heterozygous diploid clone had come to dominate the population. However, we found that these populations do sporulate (unpublished results) and observed haploid spores (using fluorescence microscopy) that predominate after sporulation. Thus, it may be that these populations in fact represent instances in which one or two beneficial haplotypes fixed across the genome. This is possible due to the highly polygenic nature of adaptation that we observed.

